# The structural heterogeneity of α-synuclein is governed by several distinct subpopulations with interconversion times slower than milliseconds

**DOI:** 10.1101/2020.11.09.374991

**Authors:** Jiaxing Chen, Sofia Zaer, Paz Drori, Joanna Zamel, Khalil Joron, Nir Kalisman, Eitan Lerner, Nikolay V. Dokholyan

**Author notes:** Equal contributors (Jiaxing Chen, Sofia Zaer).

## Abstract

The intrinsically disordered protein, α-synuclein, implicated in synaptic vesicle homeostasis and neurotransmitter release, is also associated with several neurodegenerative diseases. The different roles of α-synuclein are characterized by distinct structural states (membrane-bound, dimer, tetramer, oligomer, and fibril), which are originated from its various monomeric conformations. The pathological states, determined by the ensemble of α-synuclein monomer conformations and dynamic pathways of interconversion between dominant states, remain elusive due to their transient nature. Here, we use inter-dye distance distributions from bulk time-resolved Förster resonance energy transfer as restraints in discrete molecular dynamics simulations to map the conformational space of the α-synuclein monomer. We further confirm the generated conformational ensemble in orthogonal experiments utilizing far-UV circular dichroism and cross-linking mass spectrometry. Single-molecule protein-induced fluorescence enhancement measurements show that within this conformational ensemble, some of the conformations of α-synuclein are surprisingly stable, exhibiting conformational transitions slower than milliseconds. Our comprehensive analysis of the conformational ensemble reveals essential structural properties and potential conformations that promote its various functions in membrane interaction or oligomer and fibril formation.

## Introduction

α-Synuclein (α-syn) is a 140-residue protein containing an N-terminal segment (residues 1~60), a hydrophobic non-amyloid-β component (NAC) segment (residues 61~95), and an acidic C-terminal segment (residues 96~140)^1^. It is encoded by the SNCA gene in humans and is mainly expressed in the presynaptic terminals of neurons^2^. Endogenous α-syn regulates neurotransmitter release and synaptic vesicle homeostasis through the interaction with synaptic vesicle membranes^3–6^. An imbalance of this α-syn-membrane interaction may induce the formation of α-syn fibrils^6^. These ordered aggregates of α-syn serve as a precursor in Lewy body formation, which is a hallmark of several neurodegenerative diseases (e.g. Parkinson’s disease (PD), PD dementia, dementia with Lewy bodies, and multiple system atrophy)^7–11^. The different roles of α-syn in normal synaptic activity and as a factor in the advancement of PD may stem from different manifestations of its structure. As an intrinsically disordered protein (IDP), α-syn is characterized by a heterogeneous ensemble of dynamic conformations^12^. Although monomeric α-syn is structurally flexible, it can form ordered structures under different conditions. In the membrane-bound state, α-syn adopts two dynamically interconverting^13^ conformations: a broken antiparallel α-helix structure (residues 3~37 and residues 45~92)^14^, and an extended helix (residues 1~97)^15^. Physiologically, α-syn appears as a helically folded tetramer (α-helices shown in the initial 100 residues), which is stable and resistant to protein aggregation^16–19^. These two conformational states of α-syn mainly composed of α-helices are essential for physiological functions^20^ and disease prevention^18^. On the other hand, α-syn oligomers and fibrils, which are toxic to neuronal cells, are characterized by β-sheets rather than α-helices^21–24^. These observations link the different roles of α-syn to its various conformational states. Can the structures of the α-syn subunit in these complexes be identified also as conformational states in the structural ensemble of the α-syn monomer? If yes, a detailed structural characterization of the α-syn monomer conformations would shed light on the specific mechanisms that lead α-syn to different functional routes.

Characterizing the structurally heterogeneous ensemble of α-syn conformations is particularly challenging because it is an IDP. Experimental methods such as nuclear magnetic resonance (NMR), small-angle X-ray scattering (SAXS), and circular dichroism (CD) frequently report the conformational ensemble average^25^, which is a superposition of multiple distinct states. Due to the high concentration of proteins required in some experiments, the structural information provided might be of oligomers rather than solely of the monomer^26^. Finally, these methods (except for NMR) cannot provide atomic-resolution structural models of the conformations. Molecular dynamics (MD) simulations have been used to investigate the details of α-syn conformations. Traditional force fields used in MD, however, are designed for folded proteins^27^. Simulations of IDPs using these force fields usually generate structures that are too compact and contradict experimental data^28–31^. Moreover, computational exploration of the structural ensembles of IDPs is often prohibitive due to the vast allowable conformational space. To overcome the force-field limitations and enhance sampling, experimental data from NMR^32^, SAXS^33^, and cross-linking mass spectrometry (XL-MS)^26^ have been integrated with MD simulations^34^. These experimental methods, however, either report ensemble averages over structurally heterogeneous states or are limited by the length of crosslinkers^34^. Time-resolved fluorescence resonance energy transfer (trFRET) reports a distribution of inter-residue distances rather than the single average of the ensemble^34^. Therefore, we use data previously generated from trFRET experiment^35^ to guide discrete molecular dynamics (DMD) simulations^36–38^ of the α-syn monomer.

Previous studies using experimentally-restrained simulations to characterize the α-syn structural ensemble provided insights into the long-range interactions and secondary structures that stabilize α-syn or stimulate early stages of aggregation^26,32,33^. The long-range interaction between the N- and C-terminal segments was proposed to expose the hydrophobic NAC segment and therefore, promote protein aggregation^33^. However, recent studies showed that such interaction protects α-syn from aggregation^39,40^. Whether this interaction protects or exposes the aggregation-prone NAC segment needs to be further investigated. A study based on paramagnetic relaxation enhancement (PRE) measurement showed that the multiple conformations of the α-syn monomer interconvert in the time range of nanoseconds to microseconds^39^. However, such rapid conformational dynamics and the outcoming short-lived conformations may decrease the probability of ligand binding to a specific conformation. Knowing the correct timescale of conformational transitions is important for understanding the functional fate of the α-syn monomer. Furthermore, although the conformational ensemble of α-syn has been studied^26,32,33,41–44^, the molecular mechanisms of forming diverse conformational states (e.g. membrane-bound, dimer, tetramer, oligomer, and fibril) from the monomer ensemble are still obscure. The representative structural subpopulations of the monomer that are associated with specific cellular events (e.g. membrane binding, nucleating fibrillization), have not been fully characterized yet. Identifying these subpopulations or representative monomeric structures is significant as the concrete structural features could be used to develop small molecules that regulate α-syn functions or interfere with disease progression.

Here, we use existing trFRET data as experimental restraints in DMD simulations to map the conformational ensemble of α-syn. We confirm the predicted conformational ensemble using far-UV CD spectra and XL-MS experimental data. The results of single-molecule protein-induced fluorescence enhancement (smPIFE) measurements suggest that some of the structural sub-populations, or groups of them, are stable enough to have interconversion transitions to occur in milliseconds. This finding suggests that the α-syn monomer has stable local structures. The simulations reveal that structural subpopulations of the α-syn monomer include structures that are similar to the structures of the α-syn subunit in several well-documented complexes. Therefore, these structures might promote α-syn complexation including membrane binding and the formation of various oligomers and fibrils. Taken together, we propose stable α-syn local structures are promoting specific complex formations, and, thus, link to a structure-function relationship.

## Methods

### Determination and optimization of discretized potential energy functions used as restraints

The workflow of designing discretized potential energy functions used as restraints in DMD simulations of α-syn is illustrated in Fig. S1. First, we retrieve the distance distributions of 8 residue pairs from analysis results of bulk trFRET data previously recorded by Grupi and Haas^35^. We then convert these distributions to potential energy curves in the reaction coordinate between the residues, using the approach introduced by Haas and Steinberg^45^. We discretize the resulting potential curves to produce multi-step square-well functions^36–38^. Usually, most populated distances have lower potential energies, while distances with low probabilities have less attractions. We add the designed multistep square-well functions into the *Medusa* force field for DMD simulations^46^. The addition of these eight potentials restrains the positions of the corresponding residue pairs during simulations and reduces the degrees of freedom of the molecule. After simulation, we calculate the eight inter-residue distances for all the structures of system equilibration and plot distance distributions for each residue pair (defined here as computational/simulated distance distributions). Next, we compare the computational and experimental distance distributions. DMD simulations using this designed multi-step square-well potential generate structures with much shorter distances as compared to the distance distribution curve characterized by trFRET. To match the long-distance distribution, we modify the previous square-well functions using the following rules: (i) if the simulated distance distribution has a higher probability than the experimental one at a given range of inter-residue distances, we increase the potential energy at these distances, which then reduces the attractions in simulations; (ii) if the simulated distance distribution has a lower probability than the experimental one, we decrease the potential energy at these distances, which in turn allows more attractions. Then, we use the newly designed multi-step square-well functions for DMD simulations. After several iterations of optimization, our designed potential converges to a set of square-well functions (Fig. S2) that corresponds to an ensemble with the desired distance distributions (Fig. S3). The distance distributions of the eight residue pairs of the simulated structures do not perfectly match the experimental data (Fig. S3). This mismatch could be attributed to the skewed Gaussian distribution of the distances assumed by the trFRET experiments^35,47,48^. The simulations recapitulate most of the experimental distance distributions and therefore, provide structural insights of α-syn conformational ensemble.

### Discrete molecular dynamics simulations of α-synuclein

Discrete step function potentials are used in DMD to define inter-atomic interactions, rather than continuous potentials widely adopted in traditional MD simulations^36–38^. This application greatly reduces calculations and therefore, enhances sampling and allows explorations of the dominant conformational space in a practical timescale^26^. During DMD simulations, we use the designed multi-step square-well functions as constraints for each residue pair^26,50,51^ by integrating them with the *Medusa* force field^36–38,46,49^. We use a fully extended structure of α-syn to initiate simulations. We perform the all-atom replica exchange DMD simulations with 26 replicas, which include temperatures evenly distributed from 0.35 to 0.60 kcal/(mol·k_B_)^37^. The simulation temperature of each replicate exchanges based on the *Metropolis* algorithm, which allows structures trapped at local stable states to bypass enthalpic barriers, and hence, enhances conformational sampling^38,52,53^. We perform four million DMD time steps (~50 ps each)^38^ for each simulation. After simulation, we calculate the exchange rates between adjacent replicas to ensure appropriate replica spacing and sampling^38^. The rates in the range of 0.25 and 0.7 (Table S1) mean that the temperature exchange is sufficient for adequate sampling. We plot the system’s energy distribution specific heat curve using the weighted histogram analysis method to indicate the convergence of the system (Fig. S4)^54^.

### Simulation data analysis

We discard the first 0.5 × 10^6^ time steps of simulations during equilibration. Then, we calculate the radius of gyration (R_g_) for all the remaining structures in the 26 trajectories. We extract the highly populated structures and cluster them using the agglomerative hierarchical clustering method based on the *ward* linkage algorithm from TTClust^55^. *Ward*’s method determines a linkage between two clusters when the integration of these clusters causes the least increase in overall within-cluster variance^56^. We choose the clustering cutoff following the suggestions by Offutt *et al.*^57^: (i) the total number of clusters is less than 40; (ii) 90% of the structures are included in less than 7 clusters; and (iii) minimal number of clusters have only one structure. Specifically, first we rule out the cutoffs that generate too many clusters (more than 40), and then we increase the cutoff until we find one in which 90% of the structures are included in fewer clusters (~7). After structural clustering, TTClust also generates a clustering dendrogram, a bar plot showing number of structures within each cluster, and a table including pairwise root-mean-square deviation (RMSD) between clusters^55^. We utilize GROMACS’ *sham* program to plot the Gibbs free energy landscape by using two of the three components including RMSD to the average structure of the ensemble, the R_g_, and the potential energy. Specifically, GROMACS projects all the conformations onto a two-dimensional plane in which the axes correspond to the two selected features. Then, the program counts the number of structures occupied by each cell. It assigns the bin (in the two-dimensional plane) populated by the largest number of conformations as the reference bin and sets the free energy to zero. GROMACS calculates the free energies for the other bins using the equation: ΔG = −k_B_Tln[P(x,y)/P_max_], where P(x,y) represents the probability of structures in a bin with respect to all the structures, P_max_ is the probability of the reference bin, k_B_ is the Boltzmann constant, and T is the temperature. We compute the secondary structure composition of the predicted conformational ensemble of α-syn using GROMACS’ *DSSP* program^58,59^. We calculate the solvent accessible surface area (SASA) for each residue using the *SASA* algorithm in GROMACS^60^. We characterize the contact frequency maps for each cluster by Contact Map Explorer (https://contact-map.readthedocs.io/en/latest/examples/nb/contact_map.html).

### Expression and purification of α-synuclein

We follow the protocol introduced by Grupi & Haas^35^ for the expression and production of recombinant α-syn variants (WT and different single cysteine mutants V26C, Y39C or A56C). Briefly, we clone α-syn variants into pT7-7 vectors. Then, we transform the resulting plasmids verified by sequencing into BL21(DE3) cells (Novagen) for protein expression. We grow BL21(DE3) cells in Luria-Bertani media in the presence of 0.1 mg/mL ampicillin. We induce protein expression by adding 1 mM isopropyl β-d-1-thiogalactopyranoside (Sigma) when the cell density reaches an OD_λ=600nm_ of 0.6 ~ 0.7. After induction, we culture the cells at 37 °C for 5 h and harvest the cells by centrifugation at 6000 rpm. We resuspend the cell pellet in lysis buffer (30 mM Tris–HCl, 2 mM ethylenediaminetetraacetic acid (EDTA), 2 mM dithiothreitol (DTT), pH 8.0 - also called buffer A - and 40% sucrose) and stir it for 20 minutes at room temperature. Then, we transfer the solution to 50 mL tubes and centrifuge for 0.5 hour at 11,000 rpm. Finally, we resuspend the cell pellet in dissolution buffer (37 μL saturated MgCl_2_ solution and 90 mL of buffer A) to perform osmotic shock.

We precipitate DNA by adding streptomycin sulfate (Sigma) to a final concentration of 10 mg/mL. We stir the mixture for 20 minutes at room temperature and centrifuge at 12,000 rpm for 0.5 hour. We collect the supernatant and precipitate proteins by adding 300 mg/mL ammonium sulfate (Sigma). We stir the solution for 30 minutes at room temperature and centrifuge again at 12,000 rpm for 0.5 hour. We collect the protein precipitate and resuspend in buffer A. To remove α-syn aggregates and large oligomers, we filter the solution through a 100 kDa molecular weight cutoff (MWCO) Amicon tube. We dialyze the filtered solution using 3.5 kDa MWCO dialysis bags at 4 °C overnight against buffer A. Then, we load the solution on a 1 mL MonoQ column (GE Healthcare) using an FPLC system (Äkta Explorer). We elute α-syn with a salt gradient from 0 to 500 mM NaCl. We verify the presence of α-syn in the fractions by measuring absorption spectrum in the wavelength range of 260-350 nm and running the samples in a 12% SDS-PAGE Gel. We stain the gels by fast seeBand (Gene Bio-application). We unify relevant fractions and dialyze it with 3.5 kDa MWCO dialysis bags at 4 °C overnight against buffer A. We verify the molecular mass of the recombinant α-syn by intact protein mass determination (next section). We evaluate the purity of α-syn by running a 12% SDS-PAGE gel. We use a 12% native PAGE gel to probe the presence of α-syn dimers. We store protein samples at −20 °C.

### Intact protein mass determination

We dissolve proteins in 40% acetonitrile, 0.3% formic acid (all solvents are MS-grade) at a concentration of 2-5 mg/mL. Then, we inject the dissolved proteins directly via a HESI-II ion source into a Q Exactive Plus (Thermo Fisher Scientific) mass spectrometer and obtain a minimum of three scans lasting 30 seconds. The scan parameters are: scan range 1,800 to 3,000 m/z without fragmentation; resolution 140,000; positive polarity; AGC target 3×106; maximum injection time 50 ms; spray voltage 4.0 kV; capillary temperature 275 °C; S-lens RF level^61^. We perform scan deconvolution using Mag Tran version 1.0.3.0 (Amgen).

### Cross-linking mass spectrometry of α-synuclein

We dissolve BS^3^ powder (Sigma aldrich) in HEPES buffer (pH 7.18) to a concentration of 10 mM. We add the prepared BS^3^ solution to α-syn to a final concentration of 1 mM and incubate at 4 °C for 1.5 h with shaking at 600 rpm. We quench the cross-linking reaction by the addition of 20 mM ammonium bicarbonate for 20 min^62,63^. We prepare samples for MS and MS data analysis as described by Slavin *et al*^64^. We combine the XL-MS results obtained here using BS^3^ crosslinker (performed as a triplicate of experiments) with those based on other crosslinkers (ABAS, SDA, TATA, CBDPS, and EDC) from the work of Brodie *et al*^26^. To use XL-MS data to validate our predicted conformational ensemble, we calculate the distances between the crosslinked residues for all the simulated structures in each cluster and compare these distance distributions with the maximum length of the corresponding crosslinker. Structures with distances smaller than the maximum crosslinker length are validated by the corresponding crosslinker and we calculate the proportions of these validated structures in each cluster.

### Far-UV circular dichroism of α-synuclein

We measure the far-UV CD spectrum of 10 μM wild type α-syn in HEPES buffer (pH 7) at 25 °C in a CD spectrometer (J-1100ST, Jasco, Japan). We perform secondary structure estimation using Spectra Manager 2 (Jasco, Japan) in triplicate.

### Single-molecule protein-induced fluorescence enhancement

#### Experimental setup

We perform single-molecule protein-induced fluorescence enhancement (smPIFE)^61,65–69^ experiments using a confocal-based setup (ISS™, USA) assembled on top of an Olympus IX73 inverted microscope stand. We use a pulsed picosecond fiber laser (λ=532 nm, pulse width of 100 ps FWHM, operating at 20 MHz repetition rate and 150 μW measured at the back aperture of the objective lens) for exciting the Cy3 dye (FL-532-PICO, CNI, China). The laser beam pass through a polarization-maintaining optical fiber and is then further shaped by a linear polarizer and a half-wave plate. A dichroic beam splitter with high reflectivity at 532 nm (ZT532/640rpc, Chroma, USA) reflect the light through the optical path to a high numerical aperture (NA) super Apo-chromatic objective (60X, NA=1.2, water immersion, Olympus, Japan), which focuses the light onto a small confocal volume. The microscope collects the fluorescence from the excited molecules through the same objective, and focuses it with an achromatic lens (f = 100 mm) onto a 100 μm diameter pinhole (variable pinhole, motorized, tunable from 20 μm to 1 mm), and then re-collimates it with an achromatic lens (f = 100 mm). We further filter the fluorescence from other light sources (transmitted scattering) with a 585/40 nm band-pass filter (FF01-585/40-25, Semrock Rochester NY, USA) and detect it using a hybrid photomultiplier (Model R10467U-40, Hamamatsu, Japan), routed through a 4-to-1 router to a time-correlated single photon counting (TCSPC) module (SPC-150, Becker & Hickl, GmbH) as its START signal (the STOP signal is routed from the laser controller). We perform data acquisition using the VistaVision software (version 4.2.095, 64-bit, ISS™, USA) in the time-tagged time-resolved (TTTR) file format. After acquiring the data, we transform it into the photon HDF5 file format^70^ for easy dissemination of raw data to the public.

#### Preparation of Cy3-labeled α-synuclein

Sulfo-Cy3 (Invitrogen) linked to the maleimide thiol-reactive group, is coupled specifically to single cysteine residues in α-syn mutants via thiol coupling reaction. We reduce the thiols of cysteine residues in α-syn mutants by 2 mM DTT (Sigma) for 1 hour at room temperature. Then, we remove DTT with two rounds of dialysis, first round against buffer A and the second one against 50 mM HEPES pH7.2, 2 mM EDTA, using dialysis bags with 3 kDa MWCO. We further activate the cysteine thiols by 50 μM Tris(2-carboxyethyl)phosphine hydrochloride (TCEP, Sigma) for an hour at room temperature. Then, we add the Cy3 dye to the protein at a molar ratio of 5:1 (dye:protein). We stir the reaction mixtures for five hours at room temperature in the dark. Then, we terminate the reaction by adding 2 mM DTT. We remove excess dyes from the solution via three rounds of dialysis against buffer A using dialysis bags with 3 kDa MWCO. Afterwards, we further purify the dye-labeled α-syn using a size exclusion column HiTrap Desalting 5 mL x 5. We assess the concentration of pure labeling products by measuring the absorption of the dye (Cy3 with an absorption coefficient of 162,000 M^−1^cm^−1^). Then, we further characterize the labeling products by 12 % SDS-PAGE gels based on molecular mass.

#### Measurement

We add 100 μL samples containing 25 pM Cy3-labeled α-syn in measurement buffer (10 mM Tris, 1 mM EDTA, pH 8, 50 mM NaCl, 10 mM cysteamine, and 1 mM (±)-6-Hydroxy-2,5,7,8-tetramethylchromane-2-carboxylic acid, TROLOX) to a microscopy well (μ-Slide 18 Well, Ibidi, GmbH), first rinsed with 1 mg/mL BSA, and then sealed with the microscopy well cover. Data acquisition lasts 1-2 hours per sample. The measured samples include α-syn V26C, Y39C, or A56C mutants labeled with Cy3, as well as Cy3-labeled ssDNA and dsDNA (the lacCONS promoter with its template labeled with Cy3 linked to either register −1, +2 or +6)^68,69^.

#### Single-molecule burst analysis

We analyze all photon HDF5 files carrying photon timestamps using the FRETbursts software^71^, by first estimating the background rate per each 30 second of the measurement. Then, by using the sliding window algorithm^72–74^, we identify single-molecule photon bursts as having instantaneous photon rates larger than F (=6) times the background rate. We further select the identified bursts for having more than 20 photons. We store the burst data (e.g. burst sizes, widths/durations, separation times between consecutive bursts, background counts). We use the photon detection times relative to the moment of excitation (the photon nanotimes) of each single-molecule burst, to calculate the mean nanotimes of single-molecule bursts, which is equivalent to the average intrinsic fluorescence lifetime, and is referred to in the text as the burst average lifetime. We calculate it as the algebraic average of all photon nanotimes in the burst, that are higher than a minimal time threshold (the time beyond which the impulse response function, IRF, ends, and the fluorescence decay becomes exponential), and subtracted from that minimal time threshold. We use histograms of burst average lifetime to exhibit the subpopulations based on the fluorescence lifetime of Cy3.

We use the separation times between consecutive bursts, termed the burst separation times, to distinguish between consecutive bursts of two different molecules (times inversely dependent on the concentration; seconds in these measurements), and consecutive bursts of the same molecule recurring back in the confocal spot (times independent of concentration and depend only on the diffusion coefficient; milliseconds in these measurements). Inspired by the concepts behind burst recurrence analysis used in assessing conformational dynamics in single-molecule Förster resonance energy transfer (smFRET) experiments^75^, we report on dynamics of the Cy3 fluorescence lifetime in the following manner: the burst average lifetimes of pairs of consecutive bursts separated by less than 100 ms are compared, to qualitatively report on transitions that occur between lifetime subpopulations within 100 ms. We deposit all smPIFE analyses and raw data in *Zenodo* (DOI: 10.5281/zenodo.4081424).

## Results

### The conformational ensemble of α-synuclein predicted by trFRET-guided DMD (trFRET-DMD) simulations

We construct the conformational ensemble of the α-syn monomer by trFRET-DMD (*Methods*). We select the most populated structures (Fig. 1a) and perform hierarchical clustering, which results in the division of the conformational ensemble of α-syn into eight distinct clusters (Fig. S5 and Fig. S6; further discussion in *Methods*). The average pairwise RMSD between the clusters is 16.7 Å (Table S2) indicating the heterogeneity of the conformational ensemble. The heterogenous conformations are also demonstrated by the centroid structures (the structure with the lowest overall RMSD to the other structures in a cluster) of the eight clusters. Some of these centroids are compact while others are more expanded and disordered, as judged by R_g_ (Fig. 1a). The average pairwise RMSD within each cluster (Table S3) shows that most of the clusters are structurally heterogeneous to different degrees, and the structures within these clusters can deviate relative to their centroids. However, clusters 3 and 7 tend to be less heterogeneous and include structures that do not deviate much from their centroids. To visualize the relation of the eight clusters to the physical states they represent, we calculate Gibbs free energy maps (*Methods*) using pairs of the following parameters: (i) the level of compactness, as represented by R_g_, (ii) the RMSD relative to the average structure of the ensemble, and (iii) the potential energy from DMD (Fig. 1b). The centroids of clusters 1 and 3 have similar levels of compactness relative to the RMSD and hence are not separate in the free energy map (R_g_ vs. RMSD; Fig. 1b, left panel). However, when the free energy is calculated using the potential energy and the level of compactness (potential energy vs. R_g_) or using the potential energy and the RMSD (potential energy vs. RMSD), the centroids of clusters 1 and 3 are close yet separate in the free energy maps (Fig. 1b, middle and right panels, respectively). This feature has occurred also between the centroids of clusters 2 and 8, when comparing the different two-dimensional representations of the free energy maps. This observation is due to reduced dimensionality in our representation of the reaction coordinates, represented here by three measures (R_g_, RMSD, and the potential energy) out of many other possible representations. Indeed, visual inspection of the structures of centroids 1 and 3 as well as centroids 2 and 8 show they have markedly different secondary structures and three-dimensional organizations (Fig. 1a). The centroids of other clusters, however, are distributed at different R_g_, RMSD and potential energy positions on the free energy maps (Fig. 1b). Therefore, these well-separated clusters represent structurally distinct conformations as judged by their levels of compactness and by the RMSD. The centroid structure of cluster 6 locates at regions with higher Gibbs free energy and the structure has a low level of compactness (R_g_ ≈ 21.1 Å). This expanded structure might contribute to its high free energy and therefore, might appear less frequently than the other centroids in the ensemble. The centroid of cluster 7 is found to be located near the area with the lowest Gibbs free energy. It contains a β-sheet structure, which might stabilize the conformation. Indeed, this centroid is compact (R_g_ ≈ 15.4 Å). After performing this computational procedure guided by trFRET-derived distance distributions, we add a validation layer against additional experimental information, which is summarized in the next section.

**Fig. 1:**
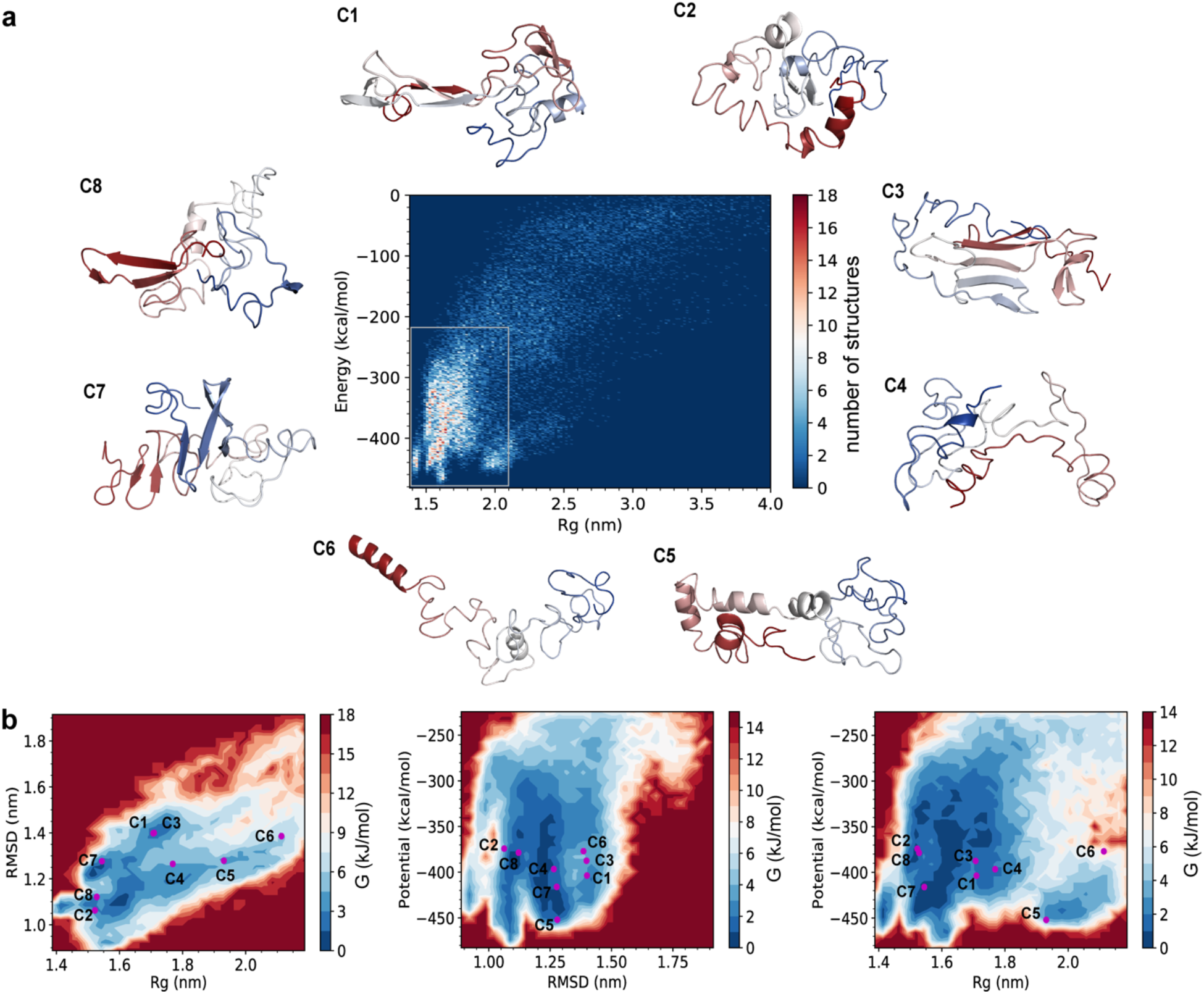
Eight distinct clusters are computationally resolved. **a**, Energy versus radius of gyration (R_g_) for the structural ensemble and the centroid structures of the eight clusters (C1-8). The structures are colored from red (N-terminus) to blue (C-terminus). **b**, The energy landscapes of α-syn based on RMSD, R_g_, and/or potential. RMSD indicates the distance between each structure and the average structure of the conformational ensemble.

### Experimental validation of the predicted conformational ensemble

To validate the simulated ensemble structure of α-syn, we conduct orthogonal experiments using CD and XL-MS and compare the computational results with experimental data in terms of secondary structure composition and intramolecular proximities. Analysis of the far-UV CD spectrum shows that monomeric α-syn in solution consists of 6.1% α-helix, 34.1% β-strand (β-sheet and β-bridge), 17.5% turns, and 42.3% other structures (random coil, bends, 5-helix and 3-helix; Fig. 2, panels a & b). To compare the secondary structure content of predicted conformational ensemble with CD data, we quantify secondary structures for every structure in the 8 clusters. The ensemble average of the secondary structures includes 7.9±5.9% α-helix, 36.3±9.3% β-strand, 20.8±6.0% turns, and 34.9±6.4% other structures (Fig. 2b). Overall, the secondary structure content of our predicted conformational ensemble agrees with the secondary structure content measured by far-UV CD experiments.

**Fig. 2:**
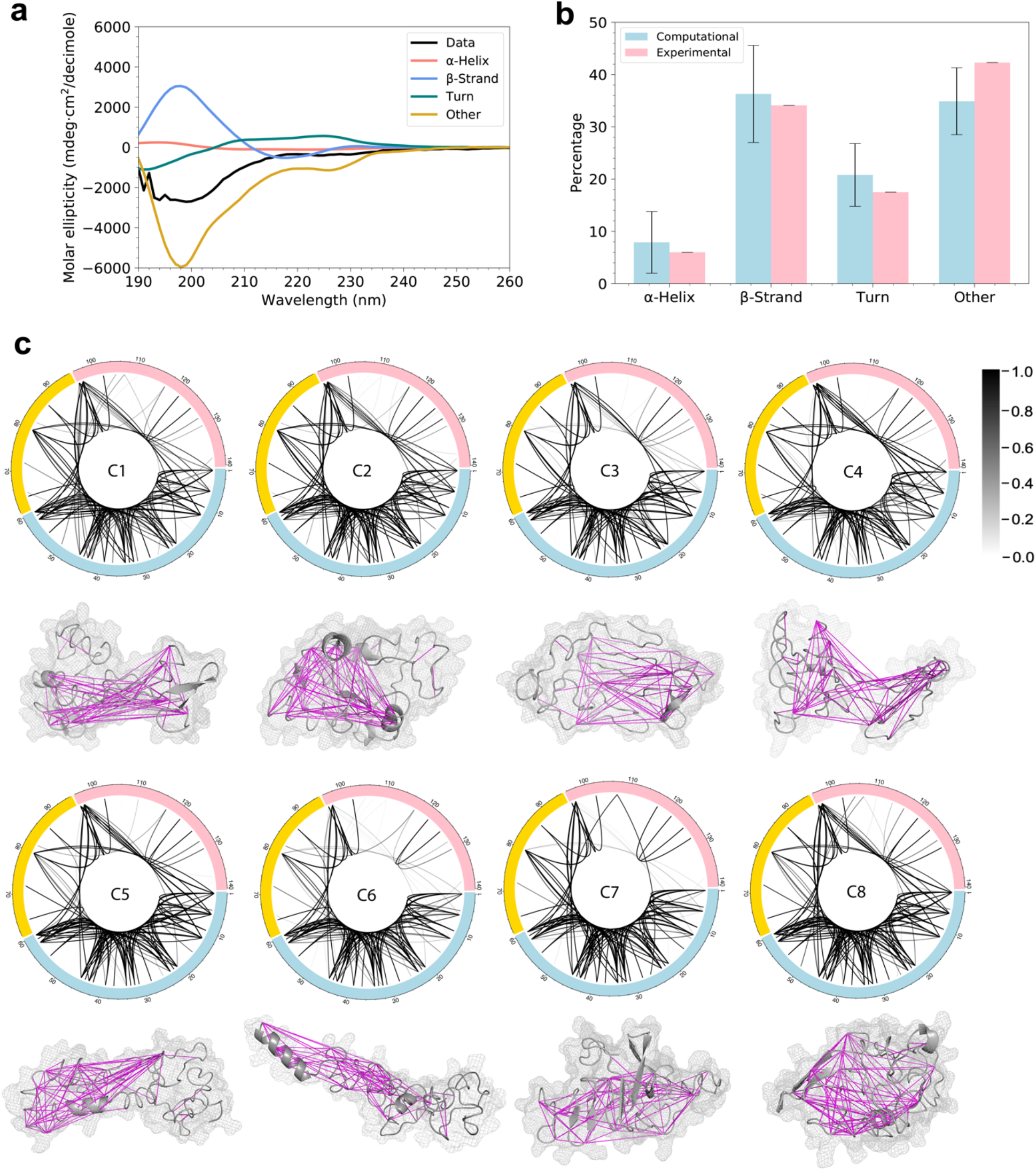
Predicted conformational ensemble is validated by far-UV CD and XL-MS experiments. **a**, The far-UV CD data showing the secondary structure content. **b**, The comparison of secondary structure content calculated from the simulated structures and from the analysis of the far-UV CD spectrum. Error bars indicate standard deviation. **c**, XL-MS connection maps of the eight clusters. Each line indicates a crosslink and the grayscale level indicates the probability of structures that satisfy cross-linking constraints within each cluster. The centroid structure of each cluster is shown under the connection map. Magenta lines indicate that these cross-linking constraints are satisfied by over 50% of the structures within each cluster. The calculated C_α_-C_α_ distance of each pair of amino acids satisfies the cross-linking constraint if it is smaller than a certain threshold: ABAS < 17 Å^76^, EDC < 10 Å^26^, SDA < 15 Å^26^, TATA < 15 Å^77^, CBDPS < 28 Å^78^, BS^3^ < 30 Å^79^.

XL-MS is often used to study local proximity of protein structures in three-dimensional space^64^. Previously, Nicholas *et al.* combined cross-linking data with DMD simulations to improve computational prediction of protein structures^26,50^. Here, we take advantage of these data together with additional XL-MS data that we collect in order to validate the characterized conformational ensemble of α-syn. A total of 125 pairs of amino acids that are crosslinked with various crosslinkers are summarized in Table S4 (*Methods*). To compare these cross-linking results with the modeled structures from DMD simulations, we calculate the distances between the 125 residue pairs and plot distance distribution for all the structures in each cluster. The maximal cross-linking length of each crosslinker is also displayed and compared with the computational distance distribution (Figs. S7-S14). The cross-linking constraints are determined to be satisfied if the maximal length falls within the computational distance distribution, since we expect the ensemble structure of α-syn includes multiple conformations and only part of the conformations are in accordance with the constraints. As shown in the figures (Figs. S7-S14), all the 125 constraints are satisfied by at least one of the eight clusters indicating that our predicted monomeric structures of α-syn agree with the XL-MS data. The probabilities of structures within each cluster that are validated by the cross-linking data are also calculated and shown in Figure 2c. Most of the cross-linking constraints are satisfied by a high proportion of structures within each cluster. The N-terminal segment is highly connected intra-segmentally and constraints within this segment are satisfied by most structures in all the 8 clusters. The difference between the clusters comes in the probability of structures that satisfy the crosslinks between amino acids of different segments as well as those within the C-terminal segment, indicating that long-range interactions between the three segments vary for each cluster and the C-terminal segment is more flexible than the N-terminal. The crosslinking arrests multiple conformational states of a protein, and therefore, not all crosslinks agree with one particular structure. Using our trFRET-DMD analysis, we deconvolute the contributions of predominant conformations to overall conformational ensemble of α-syn.

### Structural properties of α-synuclein conformational ensemble

The conformational characteristics of our predicted α-syn ensemble may provide structural insight into the molecular mechanisms that stabilize the monomer, as well as trigger protein aggregation^80^. Therefore, we analyze the structural features including secondary structures, surface exposure, and long-range interactions and compare our results to previously published data.

We compute the secondary structures and calculate the probability of each residue to adopt an α-helix, β-strand, turn, or other structure in each cluster (Fig. S15). The N-terminal and NAC segments have a higher propensity to form α-helices than the C-terminal segment, which is also captured by other studies^81,82^. Folded α-helices in these two segments are proposed to decrease the fibrillation rate of α-syn^83,84^. Overall, the formation of α-helix occupies a much lower probability than the other secondary structures. On the other hand, β-strands tend to form more frequently across the whole protein compared to the other structures. In clusters 1, 2, and 6, the central segment of α-syn has a higher propensity to form β-strand than the N- and C-terminal segments. Previous studies suggest that β-strand formation within the hydrophobic central segment may lead to oligomerization and fibril formation^26,85–89^. The average secondary structure content for each cluster (Table S5) shows that the eight clusters can be separated into two groups, one group (clusters 2, 5, 6, 8) with a higher α-helix propensity and another group (clusters 1, 3, 4, 7) with a higher β-strand propensity. Clusters with a high propensity of forming β-strands are also confirmed by their contact frequency maps (Fig. S16), exhibiting lines vertical to the diagonal. The membrane-bound state and the tetramer of α-syn form stable α-helices in the N-terminal and NAC segments, while the oligomers and fibrils have an increasing amount of β-strand. The two distinct groups with different α-helix and β-strand propensity may indicate their diverse roles in physiological functions and pathogenesis of PD.

Long-range interaction between the C-terminal and NAC segments, as well as the interaction between C- and N-terminal segments of native α-syn have been detected and confirmed by several PRE experiments^32,39,90,91^. These experiments measure the interactions between an attached paramagnetic nitroxide radical group and protons of a distance within 25 Å^92^. To determine if our conformations agree with these observations, we calculate the distances between the centers of mass of the N- and C-terminal segments, as well as the NAC and the C-terminal segments, respectively. A long-range contact is defined to occur if the distance is less than 25 Å. The distance distribution between NAC and C-terminal segments for all the structures in each cluster shows that the majority of our predicted structures have this long-range interaction (Fig. S17), which is consistent with the PRE experimental data^32^. This NAC and C-terminal contact is hypothesized to sequester the hydrophobic NAC segment from exposing in the solvent and therefore, is critical to retard protein aggregation^32,43^. The long-range interaction between the N- and C-terminal segments was also proposed to protect monomeric α-syn from aggregating by hiding the hydrophobic core^39^. A recent study showed that calcium interaction with the negatively charged C-terminal segment disrupts the N- and C-terminal interaction causing the exposure of the N-terminal, which consequently increases aggregation propensity^40^. Some studies, however, found that this long-range interaction exposes the hydrophobic NAC segment, which could induce protein aggregation rather than inhibit it^33^. To solve these two contradictory statements, first, we plot the distance distribution between the N- and C-terminal segments to confirm if this long-range interaction exists in our data. Our analysis shows that over 70% of the structures in clusters 1 (90.46%), 2 (83.69%), 3 (84.04%), 4 (73.33%), 7 (99.54%), 8 (79.35%) possess the N- and C-terminal long-range interaction, while less then 35% of the structures in clusters 5 (30.94%) and 6 (15.25%) contain this interaction (Fig. S18). Then, we extract the protein structures that have this long-range interaction from each cluster and compute the average solvent accessible surface area (SASA) for each residue. Since different amino acids have different sizes, we calculate the relative solvent accessibility by dividing its average SASA by the maximum surface area of that type of residue. A cutoff of 40% is used to define the two states, buried or exposed, as this cutoff has been proved to be useful by other studies^33,93^. Our data (Fig. S19) show that most residues in the NAC segment tend to be buried except residues in cluster 7, where 57% of the residues in the hydrophobic segment are exposed. These results indicate that for most of the monomeric conformations of α-syn, N- and C-terminal contact covers the hydrophobic NAC segment, which stabilizes the protein and protects it from aggregation. However, this long-range contact also causes the NAC segment to be exposed in a small fraction of the structures that are aggregation-prone.

### Millisecond transitions between some of the structural subpopulations observed by single-molecule protein-induced fluorescence enhancement

We have shown that the ensemble structure of α-syn can be divided into eight distinct clusters, describing the underlying structural subpopulations of α-syn. DMD simulations in the present study, however, do not capture the transition between structural subpopulations. The ensemble structure of an IDP may include several distinct conformational states with low activation barriers between them, in which transitions may occur as slow as a few microseconds. If this is the case, fluorescence measurements of freely-diffusing single molecules that yield photon bursts with durations of a few milliseconds would result in a single burst population that is averaged out (up to thousands of transitions between conformational states occurring within the time a single α-syn traverses the confocal spot). However, if the transitions between some of α-syn’s structural subpopulations occur slower than the millisecond timescale of bursts, such measurements would result in burst histograms including more than a single population. In order to test the conformational dynamics of α-syn monomer, we perform smPIFE experiments^61,65–69^.

We conjugate the dye Cy3 to a cysteine in three different single cysteine α-syn mutants (at either residues 26, 39 or 56; *Methods*). Cy3 exhibits excited-state isomerization between a bright *trans* isomer and a dark *cis* isomer, leading to an overall low fluorescence quantum yield. However, if the excited-state isomerization of Cy3 is slowed down relative to its excited-state lifetime, Cy3 dwells in the *trans* isomer longer, leading to an increase in its fluorescence quantum yield. The PIFE effect tracks that change in excited-state isomerization caused by nearby segments of the protein, which impose steric obstruction on Cy3 and limit its capability to isomerize. In PIFE measurements the increase in fluorescence quantum yield, can be observed via the increase in the average fluorescence lifetime. We measure the Cy3-labeled α-syn constructs at a low concentration, in which it is found as a monomer (25 pM), where the average fluorescence lifetime is measured one α-syn molecule at a time (when it crosses a tightly focused laser beam in a few milliseconds). Importantly, the durations of the recorded single-molecule bursts are in the range of 1-8 ms.

The resulting average fluorescence lifetimes of single-molecule bursts are collected in histograms (Fig. 3 and Fig. S20). The results clearly show the average lifetime values of α-syn monomers (Fig. 3) can be grouped into more than a single central population. The sub-population with low average lifetime values represents α-syn species with minimal intramolecular interactions in the vicinity of Cy3-labeled residue, while the sub-populations with larger average lifetime values represent species with intramolecular interactions in the vicinity of Cy3-labeled residue, sterically interfering with the free isomerization of the exited Cy3. As a control of the smPIFE measurements, we perform the same measurements on Cy3 labeling DNA at either one of three different bases along the sequence, either in double- or single-stranded form (Fig. S20). These measurements result in a single population of average lifetime values, showing that smPIFE yields a single population of values either in the case of a single rigid structure, such as in dsDNA, or in the case of a very flexible structure with rapid structural transitions, such as in ssDNA, that average-out in millisecond. Therefore, we show that at least some of the proposed eight clusters or structural subpopulations, interconvert in 1-8 ms or slower which is surprisingly far from what is expected for an IDP^13,39,42,47,94–98^.

**Fig. 3:**
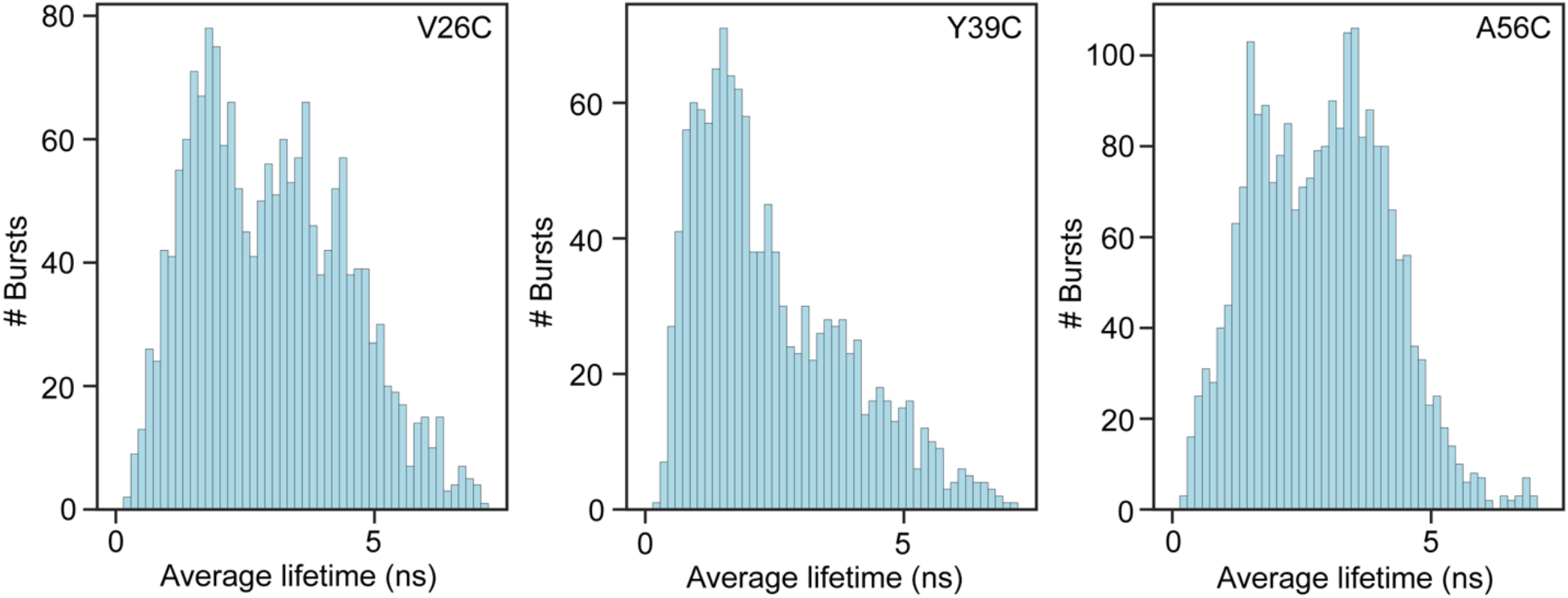
The ensemble of α-syn includes different subpopulations with various degrees of side chain accessible volume that interconvert slower than a few milliseconds (slower than burst durations). The average lifetime histograms from smPIFE measurements of Cy3-labelled α-syn (conjugated at Cys26, Cys39, or Cys56 in the V26C, Y39C or A56C mutants, respectively).

While the appearance of distinct burst-wise subpopulations is a marker of dynamics occurring slower than the burst durations, this evidence is insufficient to find the timescale of these slow transitions. To answer this question, at least qualitatively, we perform burst recurrence analysis^75^ based on the values of burst average lifetimes (Fig. S21). Briefly, there are two types of pairs of consecutive bursts: (i) different molecules, where the pair of bursts separated by longer separation times, t_sep_, the lower is the concentration. The separation times of such pairs of consecutive bursts in single-molecule experiments are at least seconds; (ii) a recurring molecule that produces a burst when crossing through the observation volume, proceeds to move outside the observation time, and then recurs back into the observation volume to produce another burst. Such pair of consecutive bursts occurs rarely, with burst separation times similar to the burst durations (few ms) that are independent of the concentration. If the two types of burst separation times are well-separated (Fig. S21, panel a), one can select pairs of consecutive bursts of recurring molecules and compare their burst properties. Out of such pairs of consecutive bursts, separated by less than a given t_sep_ (Fig. S21, panel a, orange shade), if the first burst has an average lifetime belonging to one subpopulation of lifetimes (Fig. S21, panels c, d) and the second burst has an average lifetime belonging to the other subpopulation of lifetimes (Fig. S21, panels c, d), then a transition occurred between these two subpopulations, within t_sep_.

For the smPIFE measurements of each Cy3-labeled α-syn, we define ranges of burst average lifetimes that are either within low or high values (Fig. S21, yellow and green shades, respectively). Then, we seek pairs of consecutive bursts separated by < t_sep_ (Fig. S21, panel a, orange shade), defined as 80 ms when Cy3 is conjugated to Cys26 in the V26C mutant, or 100 ms when Cy3 is conjugated to Cys39 or Cys 56, in the Y39C or A56C mutants, respectively. Out of these burst pairs, we also seek ones where the average lifetime of the first burst is within the low or high average lifetime subpopulations, and subsequently report on the lifetime of the second burst, for all these burst pairs (Fig. S21, panels c and d, respectively). In all three Cy3-labeled positions, a fraction of burst pairs undergoes transitions between low and high average lifetime subpopulations (Fig. S21, panel c, bursts outside the yellow shade) and vice versa (Fig. S21, panel d, bursts outside the green shade). Nevertheless, the amount of these selected recurring molecule bursts is low and we cannot further use this data to estimate the exact transition rates.

Overall, our results show that α-syn undergoes structural transitions influencing the degree by which some amino acid residues in the N-terminal segment (residues 26, 39 & 56) experience steric restriction from other parts of the protein. These transitions occur within tens of milliseconds between some of the structural subpopulations of α-syn.

## Discussion

The size of the α-syn monomer in aqueous solution was studied extensively by various experimental techniques including PRE, SAXS, and smFRET^41,99–102^. These experiments indicate that the average structural ensemble of α-syn is more compact than a random coil. However, the R_g_ values differ significantly between methods ranging from 22.6 to 50 Å^41,99–102^. This disagreement and the large R_g_ may be caused by a mixture of monomeric as well as multimeric states of α-syn under specific experimental conditions^101,103^. In addition, techniques such as SAXS produce larger errors when the R_g_ is large^100^, because the highly extended conformations contribute more to the scattering intensity^101^. Our simulation data shows that most of the monomeric structures (99.6%) have R_g_ values ranging from 14 to 50 Å, most of which (66.6%) are populated within the range of 14 and 21 Å. The ensemble structure retrieved from our simulation captures more compact structures, which is intriguing. These compact structural subpopulations may exist and be overlooked by experimental studies due to the above-mentioned issues. Therefore, we study these compact subpopulations in the attempt to understand their potential roles as the precursors to form diverse α-syn complexes and oligomers. The ensemble structure is split into eight clusters with distinct conformations by the agglomerative hierarchical clustering method (*Methods*). Our smPIFE data suggests that some of these structural subpopulations transition within tens of milliseconds, which is much slower than the reported nanosecond and microsecond timescales^39^. The slow dynamics support the notion that to carry out one of its multiple functions via ligand binding and complexation, the α-syn monomer has to stay in a specific conformation long enough to facilitate ligand binding. We suggest that the heterogenous conformations of α-syn follow a hierarchy of transition dynamics, in which 10-100 ms dynamics represents transitions between some of the structural subpopulations, while other transitions occur in microseconds or faster. PRE^92,104^, bulk time-resolved fluorescence^42,47,98^, fluorescence correlation spectroscopy^97^, smFRET^13,94–96^, and MD simulations^105^ all point to structural dynamics as slow as microseconds but lack identification of millisecond dynamics. Here, using smPIFE measurements, we identify the millisecond or slower dynamics of structural changes between some of α-syn’s structural subpopulations. This finding is unique, especially when compared to previous smFRET works, since PIFE identifies subpopulations on the basis of steric obstructions to Cy3 that range shorter (<3 nm) than the distance range covered by smFRET (3-10 nm)^61^. By that, one can assume that the slow dynamics were not identified by smFRET, because they were local, and hence did not introduce FRET subpopulations with significantly different mean values. Indeed, smFRET studies of the α-syn monomer report on a single wide FRET population when in solution^13,106–108^. Interestingly, in the presence of the kosmotropic osmolyte trimethylamine N-oxide (TMAO), the mean FRET efficiency changes, however within a single FRET population^107^. Since TMAO stabilizes compact conformations out of an ensemble of structures, these findings come to show that multiple conformations exist, but are reported within a single millisecond-averaged population. smPIFE, however, exhibited these conformational subpopulations on the basis of a reporter that captures features that are more local than what smFRET could capture, the microenvironment of Cy3 conjugated to a given amino acid residue. Bulk trFRET can track FRET between UV-excitable dyes in shorter inter-dye distances (1-5 nm)^35,42^. However, in bulk trFRET measurements, conformational subpopulations cannot be distinguished as in smFRET (only in exceptional cases)^109^. Therefore, the local nature of the reported information in smPIFE, and the fact that it can be employed on single molecules rendered it useful in uncovering the multiple structural subpopulations of α-syn and the slow dynamics between them. Taken together, these results point to a possible conformational energy landscape with a hierarchy of activation barrier heights. What is still unknown to us is which pairs of conformations/clusters undergo rapid interconversions and which do not.

The monomeric structures of α-syn may serve as the building block of its specific interactions with other ligands and proteins, and hence dictate which of its various functions it will follow. By that, the conformation of α-syn may determine its role in maintaining normal cellular functions as well as initiating protein aggregation that is detrimental to neuronal cells and promotes PD progression^110–113^. A deep investigation of α-syn structural ensemble characterized by trFRET-DMD reveals transient structures that may trigger membrane binding and protein fibrillization.

α-Syn forms two well-ordered α-helices in an anti-parallel arrangement in the N-terminal and NAC segments, followed by a disordered C-terminal segment when it interacts with micellar membranes (PDB 1XQ8^14^). To find out if our predicted conformational ensemble of the α-syn monomer contains structures similar to the membrane-bound state of α-syn, we calculated the RMSD between the pairs of C_α_ of our simulated structures in the eight clusters and the structure of the micelle-bound α-syn. We find that several structures in cluster 5 are the most similar to this micelle-bound state, based on the RMSD distribution (Fig. S22). The cluster 5 structure with the lowest RMSD (30.2 Å, p < 2.2e-16) relative to 1XQ8 displays partially folded helices in the N-terminal as well as the NAC segments (Fig. 4 and Fig. S27). These two helices are arranged in an anti-parallel direction connected by the linker at the same region as that in 1XQ8. The C-terminal segment is also characterized by the highly disordered loops (Fig. 4 and Fig. S27). Structural alignment of the 10 structures with the lowest RMSD in cluster 5 shows that a number of structures are closely related to the membrane-bound state of α-syn. The structures with the similar secondary structural composition and topology in cluster 5 might represent the monomeric precursor that interacts with the surface of a membrane and therefore, might be important for the synaptic functions. The organization of the N-terminal and NAC segments in these structures is more globular than the elongated structure of the end-product bound to the membrane. Gambin *et al.*^114^ performed smFRET measurements to track the kinetics of the conformational change upon mixing α-syn with a membrane, as viewed from the distance between the edges of the N-terminal and NAC segments, from the α-syn monomer ensemble to the membrane bound helical hairpin structure. In that study, they identified the kinetics includes an intermediate state. We propose that the first kinetic transition is the stabilization of the α-syn conformation represented by the structures in cluster 5, and the depopulation of the structures in the other clusters. Therefore, the second kinetic transition is caused by the stabilization of the elongated helical hairpin conformation by the membrane. In summary, this proposed mechanism highlights the possibility that certain structures in the structural subpopulations we identified in the α-syn monomer ensemble, serve as the preexisting intermediates for membrane binding.

**Fig. 4:**
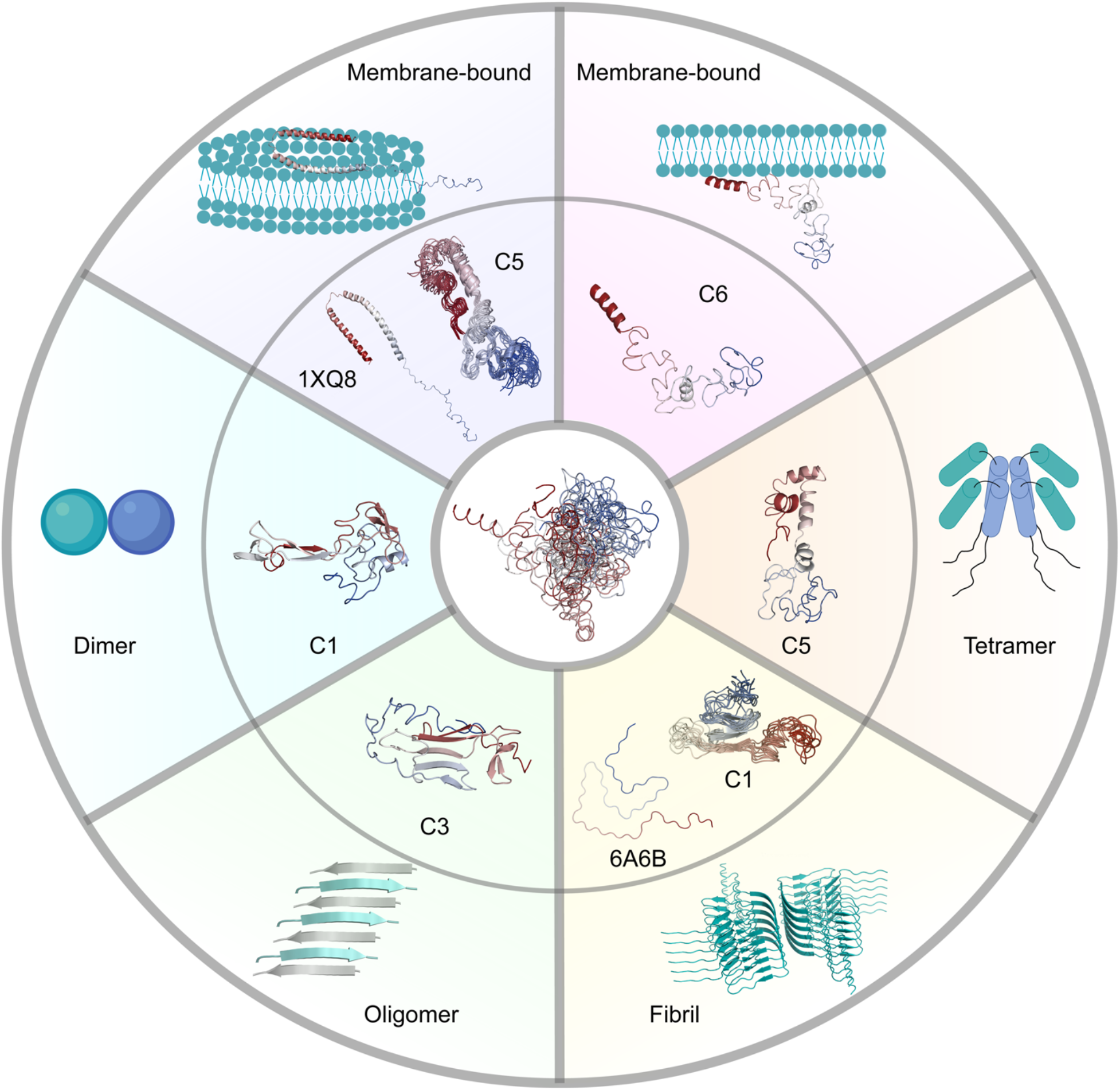
Conformational ensemble of α-syn includes dynamic structures that potentially serve as structural precursors of its various physiological and pathogenic functions. Membrane interaction and dimer, tetramer, and oligomer formation might be stimulated by clusters 6, 1, 5, and 3 (C6, C1, C5, and C3), respectively. The membrane-bound state of α-syn with broken anti-parallel α-helices might be derived from structures in cluster 5 (C5). α-Syn fibrils might be formed by structures in cluster 1 (C1). The outer panels present the structures of the α-syn subunit in several complexes or existing models in other complexes. The inner panels present the centroid structures of specific clusters or the structural alignment of the top 10 structures with the lowest RMSD to 1XQ8 or 6A6B. The central panel describes the structural ensemble of α-syn. Structures in the figure are colored from red (N-terminus) to blue (C-terminus).

Besides the helical hairpin structure (broken-helix structure) characterized by the NMR study^14^, an extended helix conformation associated with the membrane of a curvature larger than that of a micelle was also reported^15^. Based on a solid-state and solution NMR spectroscopy study^115^, the N-terminal residues (first 25 residues) are found to adopt a well-defined α-helical secondary structure that targets and anchors α-syn to the membrane of synaptic vesicles. The central segment of the protein (residues 26~97), however, is more flexible in solution and forms stable α-helix when it binds to the membrane surface^115^. The C-terminus (residues 99~140) is characterized to be extremely dynamic and unstructured^115^. By comparing these experimental data with our predicted structural subpopulations, we find that the centroid structure of cluster 6 explains these essential features (Fig. 4). Our results show that the first 15 residues form an α-helix followed by dynamic and partially folded α-helices in the NAC segment and a completely unstructured C-terminal segment at the end. The extended structure favors targeting of the N-terminal segment to the membrane and then brings the NAC segment close to the membrane, which further induces α-helical elongation. This proposed binding mechanism with membrane might result in the extended helix conformation rather than the broken-helix structure. Therefore, structures in cluster 6 might interact with membrane in an extended helix conformation. An overall analysis of the secondary structure composition in cluster 6 shows a high probability (>40%) of forming α-helices in the N-terminal and central segments (Fig. S15). This observation further suggests that cluster 6 might represent the structural subpopulation that is important for membrane binding. The low propensity of α-helical formation in some other clusters (clusters 3, 4, 7) implicates the heterogeneity of the conformational ensemble and provides evidence that the α-syn monomer includes structural subpopulations that are prone to membrane bound and unbound states.

The α-syn tetramer was reported to be the native state in eukaryotic cells, preventing it from aggregation and reducing its pathogenicity^18^. According to the results from electron microscopy and PRE experiments, a model of the α-syn tetramer formed by hydrophobic interactions of the α-helix in the central segment (residues 50~103) was proposed^16^. An α-helix in the N-terminal segment (residues 1~43) locates at the opposite direction of the hydrophobic core and forms an antiparallel arrangement with the α-helix in the NAC segment. By scrutinizing our conformational ensemble, we find that the centroid structure of cluster 5 might serve as the structural precursor of the tetramer (Fig. 4). We observe discontinuous α-helices at both N-terminal and the NAC segments. An antiparallel arrangement between these two segments is also displayed. Moreover, the C-terminal segment described as disordered in the literature is also unstructured in our prediction^16^. The overall hairpin-like structure exposes the hydrophobic residues in the NAC segment, which might facilitate oligomerization and benefit the formation of the tetramer hydrophobic core. Although the α-helices at the N-terminal and NAC segments are discontinuous, which is different from those in the tetramer, this can be easily explained by the monomeric state of α-syn. The structures of α-syn monomer are highly flexible in solution, while the α-helices in oligomers are elongated and stabilized by the hydrophobic interactions^16^. The whole analysis of α-helix composition for all the structures in cluster 5 also reveals high probabilities of forming discontinuous α-helices at the N-terminal and NAC segments (Fig. S15). We propose that these discontinuous and transient α-helices as well as the exposure of the hydrophobic NAC segment promote tetramer formation. While some structures in cluster 5 interact with membranes as discussed before, other cluster 5 structures might be involved in tetramer formation *in vivo* and protection from neurodegenerative disorders.

α-Syn oligomers and fibrils have been implicated in PD and other synucleinopathies. Understanding how the α-syn monomer forms oligomers and fibrils is critical to demystify the pathogenesis of PD, as well as, to assist in developing effective therapies. Dimer formation is considered to be the first step of oligomerization^116^. A high-speed atomic force microscopy (HS-AFM) study displayed two types of dimer conformations^81^. Type I is composed of two compact monomers, while type II consists of a compact monomer and a monomer with an extended protrusion. From the centroid structures of the 8 clusters, we find structures that are more globular and structures with extended tails. Because of the low-resolution of HS-AFM experiments, we have no information about the secondary structure composition of the dimers. Therefore, we propose that the conformational ensemble includes both compact and extended structures that are necessary for the formation of the α-syn dimer. Further studies are required to further clarify which cluster might promote dimer formation. Previous simulations suggested that the β-strand-rich hydrophobic NAC segment is important for dimerization^1^. Since the centroid structure of cluster 1 has an exposed hydrophobic β-sheet that is extended, cluster 1 might be the potential structural subpopulation that promotes dimerization (Fig. 4).

α-Syn oligomers and fibrils with varying morphologies have been reported in a large number of studies^117^. Oligomer species are mainly divided into two groups: (i) on-pathway oligomers that can grow into fibrils, and (ii) off-pathway oligomers that are resistant to fibrillation^118^. Infrared spectroscopy studies showed that the prefibrillar α-syn oligomers adopt antiparallel β-sheet structures, which is different from the parallel β-sheet structure in fibrils characterized by x-ray crystallography^22^. It was proposed that on-pathway oligomers must undergo structural rearrangements to produce fibrils. Experimental evidence has already established the connection between oligomers and fibrils. If so, how does the α-syn monomer accumulate and form various oligomers? Can the α-syn monomer directly aggregate and generate fibrils? To answer these questions, we examine the centroid structures of our eight clusters and compare various fibril structures characterized by NMR and cryo-EM with the simulated structures. We identify antiparallel β-sheet structures formed in the aggregation-prone segment (NAC) of the centroid structure in cluster 3 (Fig. 4). In addition, long-range interactions between the N- and C-terminal segments expose the NAC segment, which was also discovered in other studies and speculated to enhance protein aggregation^33^. A comprehensive analysis of the secondary structure composition reveals that cluster 3 has a higher proportion of β-strands (Fig. S15). Therefore, we deduce that cluster 3 might represent the subpopulation of forming oligomers (Fig. 4), some of which are toxic to neuronal cells and promote neurodegenerative diseases.

To understand if α-syn ensemble includes structures prone to fibrillation, we compare our simulated structures with the structures of the α-syn subunit in various fibrils based on RMSD of C_α_s. The comparison with the cryo-EM structure of α-syn fibril (PDB 6A6B^119^) shows that clusters 1, 4, 6, and 8 contain structures with lower RMSD than the other clusters (Fig. S23). We extract the structures with the lowest RMSD in these four clusters and compare them with the subunit of 6A6B. We find that the structure in cluster 1 has the lowest RMSD (11.1 Å, p < 2.2e-16) and the most similar topology compared to 6A6B (Fig. S27). The comparison of simulated structures with the other fibril structures (PDB 2N0A^120^ and 6XYP^121^) also indicates that structures in cluster 1 are the most similar to the subunits in these fibrils (Fig. S24, S25, S26, and S27). Structural alignment of the top 10 structures with the lowest RMSD also implicates that a number of structures are similar to the subunit of the fibrils (Fig. S27). Therefore, structures of cluster 1 might play a crucial role in fibril formation more than the other clusters and therefore, might represent the subpopulation that stimulates fibrillation.

Based on our predicted conformational ensemble of the α-syn monomer, we suggest potential structures that have a high propensity to bind membranes, form different oligomers, or aggregate into fibrils, which are supported by various experimental data. Experimental conditions or mutations that stabilize a specific conformational state could be explored in the future in order to verify our hypothesis. The concrete structures provided by our study may also be used in designing small molecule regulators aiming to prevent protein aggregation and interfere with the development of neurodegenerative diseases.

## Acknowledgements

We would like to thank Dr. Asaf Grupi, Dr. Dan Amir & Dr. Elisha Haas from the Mina & Everard Goodman Faculty of Life Sciences in Bar Ilan University for sharing the plasmids of α-syn bearing single cysteine mutations. We would also like to thank Dr. Asaf Grupi for fruitful discussions regarding α-syn. We acknowledge support from the National Institutes for Health 1R35 GM134864 and the Passan Foundation (to N.D.). The project described was also supported by the National Center for Advancing Translational Sciences, National Institutes of Health, through Grant UL1 TR002014. The content is solely the responsibility of the authors and does not necessarily represent the official views of the NIH (to N.D.). In addition, this project was supported by the Israel Science Foundation (grant 1768/15 to N.K.), the National Institutes of Health (NIH, grant R01 GM130942 to E.L. as a subaward), by the Milner Fund (to E.L.) and by the Hebrew University of Jerusalem (start-up funds to E.L.).

## Author contributions

J.C. & N.D. performed trFRET-restrained DMD simulations; S.Z., P.D. & K.J. prepared recombinant α-syn proteins and labeled single cysteine α-syn mutants with Cy3; S.Z. & E.L. performed smPIFE experiments and data analyses; J.Z. & N.K. performed BS3-based XL-MS experiments and data analyses; E.L. recorded far-UV CD spectrum of wild type α-syn and analyzed the secondary structure content; J.C. & E.L. devised and J.C. performed the probabilistic verification of the DMD simulation results against XL-MS experimental results; J.C. & E.L. devised and J.C. performed the verification of the DMD simulation results against far-UV CD secondary structure experimental results; E.L. provided consultation about bulk trFRET data; N.D. provided consultation about DMD simulations; J.C. wrote the paper and all co-authors assisted in refining it.

