## Supplementary information for "The structural heterogeneity of α-synuclein is governed by several distinct subpopulations with interconversion times slower than milliseconds"

^#^ Equal contributors

* Corresponding authors

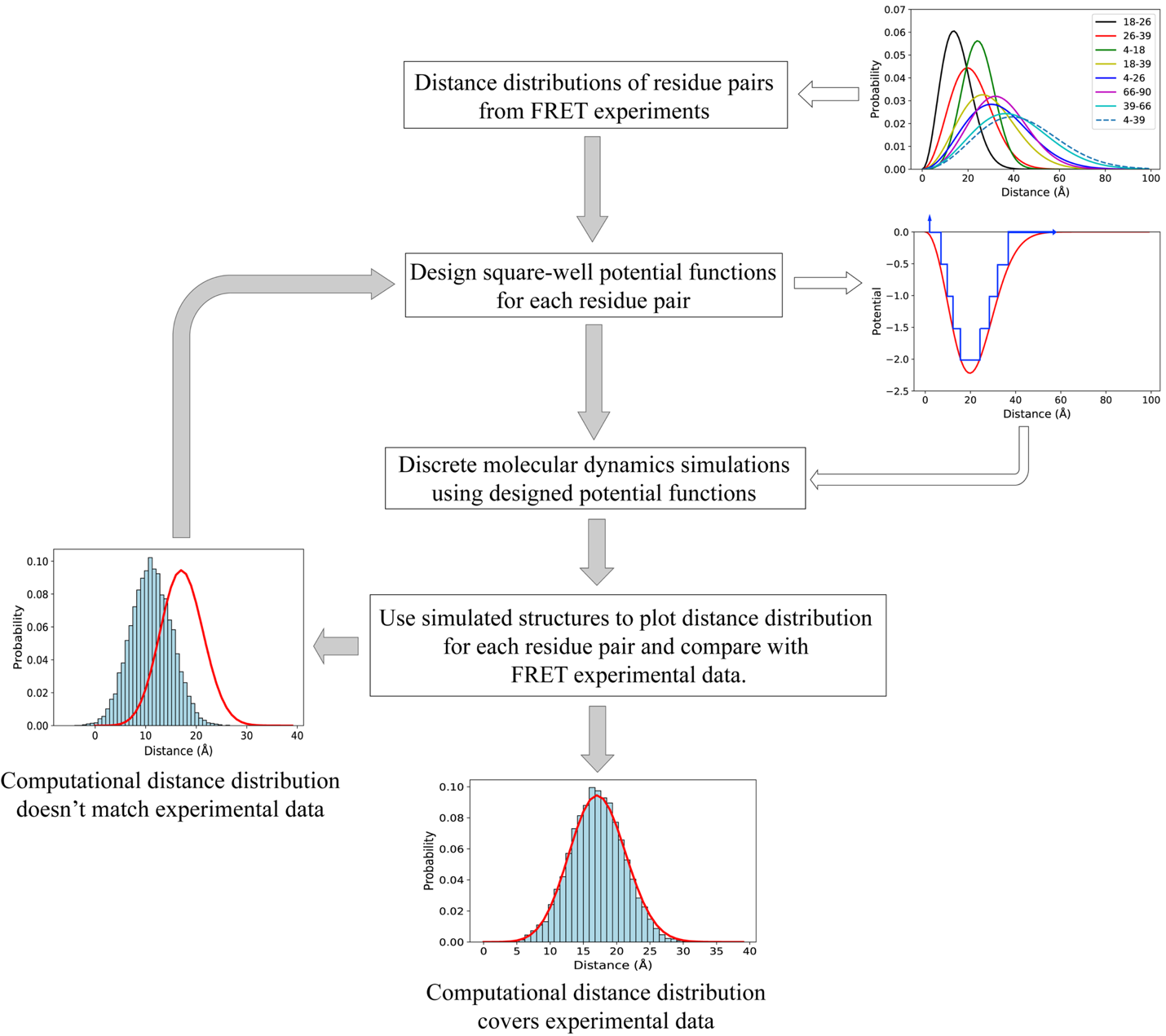

**Fig. S1.** **The schematic workflow of predicting protein conformations by the combination of trFRET and DMD simulations.**

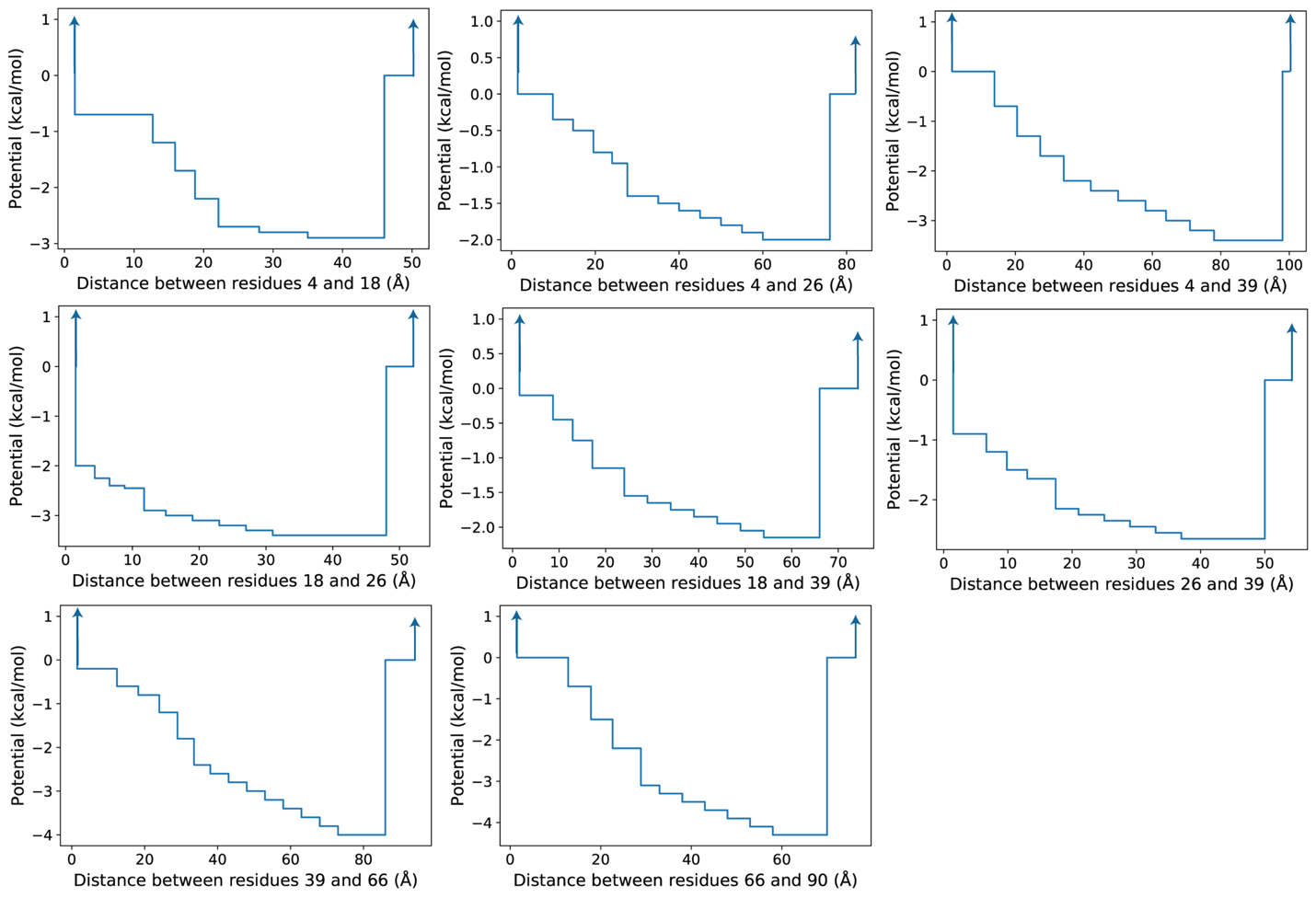

**Fig. S2. The discrete step function potential used as constraints for DMD simulations.**

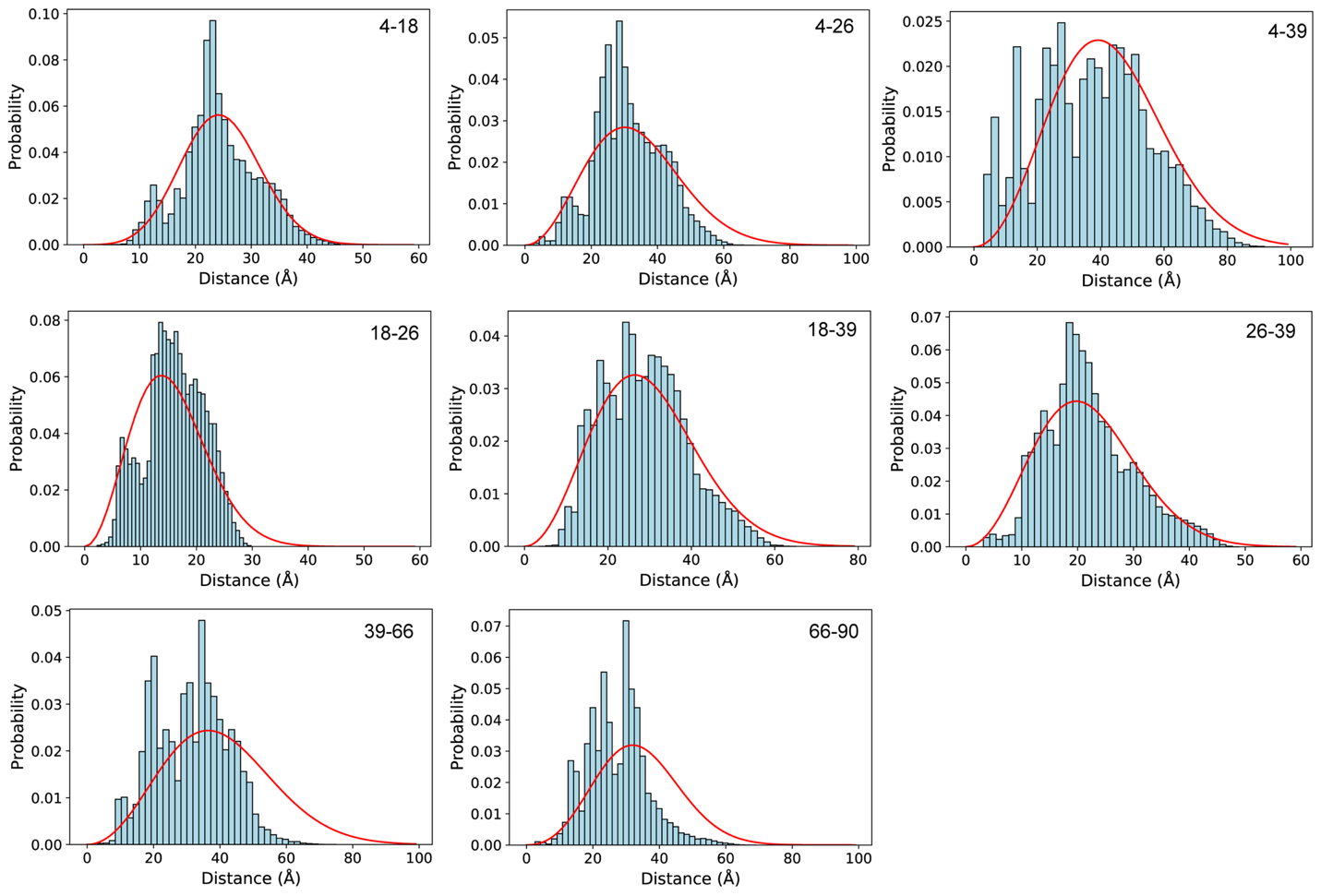

**Fig. S3. Comparison of computational (blue histogram) and experimental (red line) distance distributions**. The pair of residues are labeled in each panel.

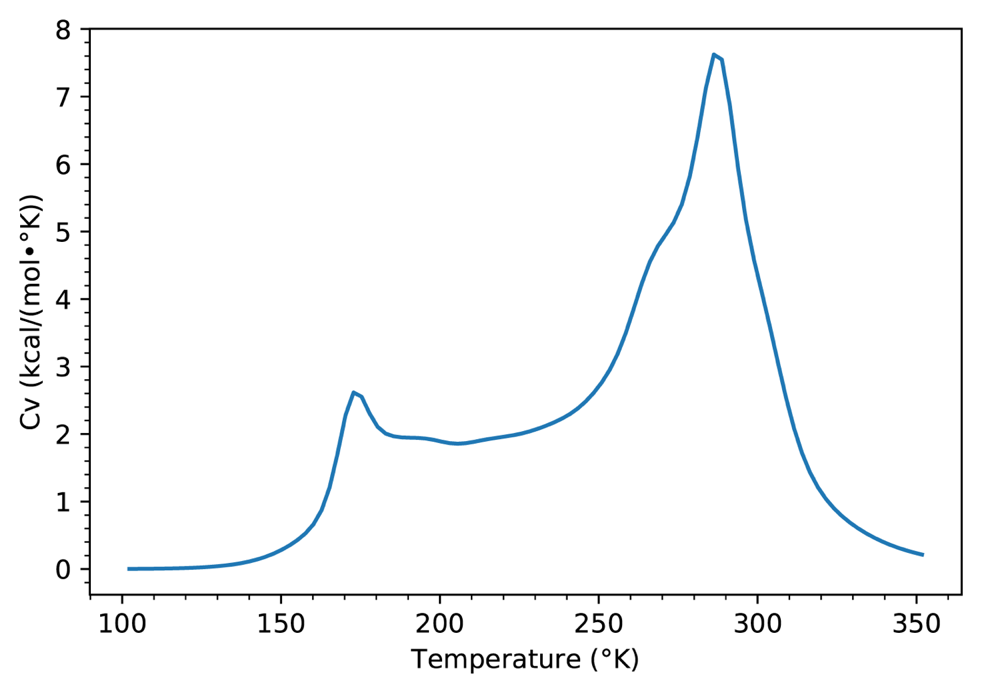

**Fig. S4. The heat capacity curve of α-syn simulations within time steps of 0.5x10^6^ ~ 4x10^6^.**

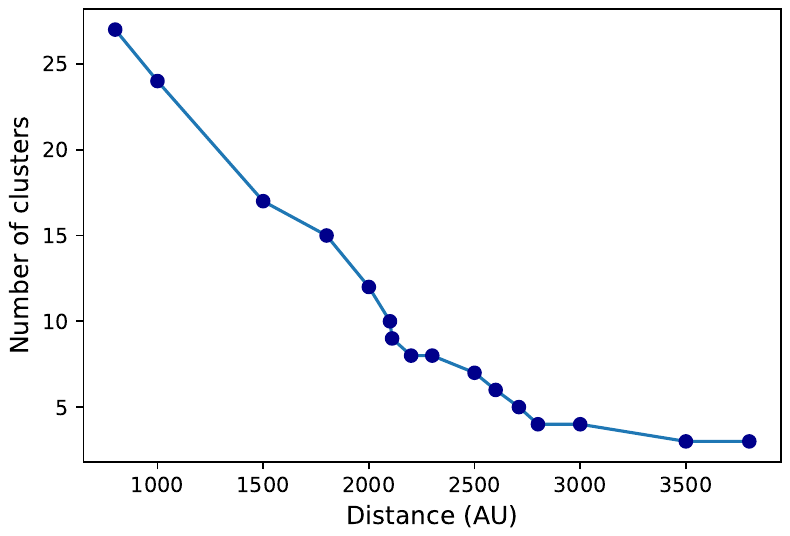

**Fig. S5. Optimization of distance cutoffs for the hierarchical clustering.**

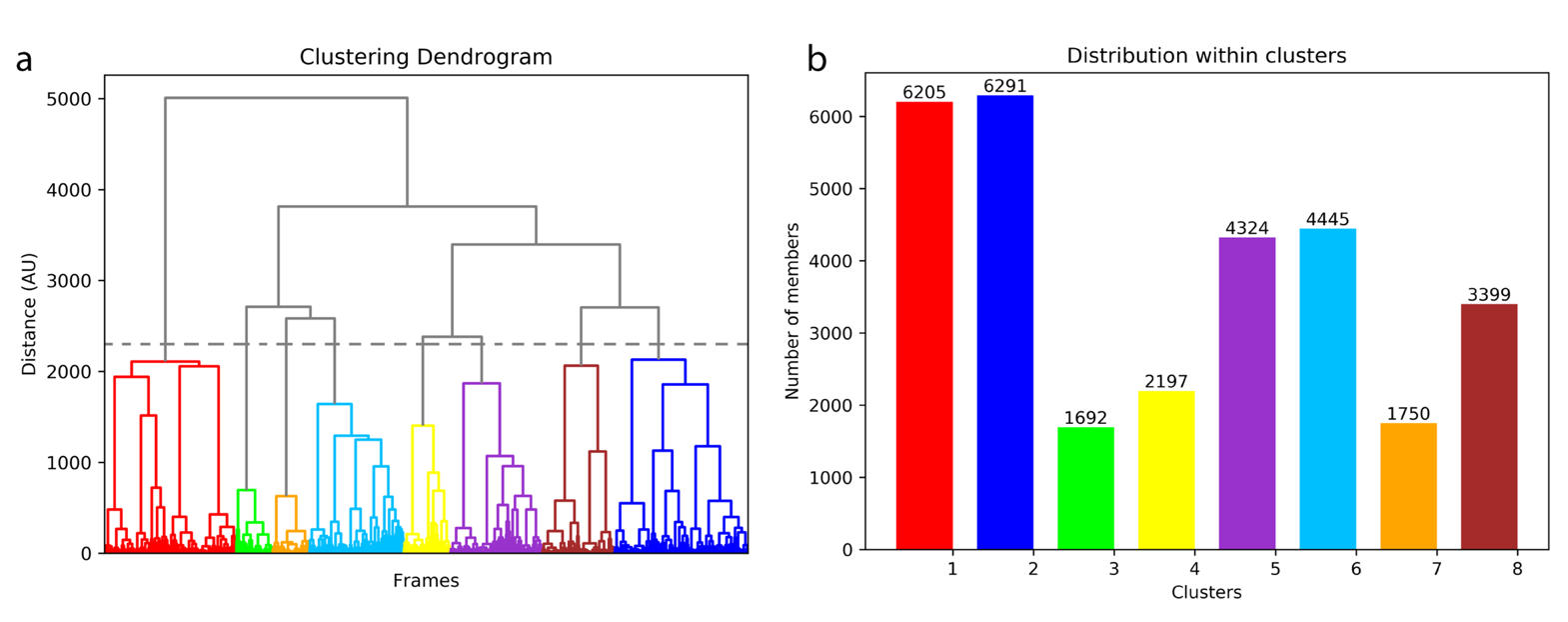

**Fig. S6. The clustering dendrogram (a) and the number of structures within each cluster (b).**

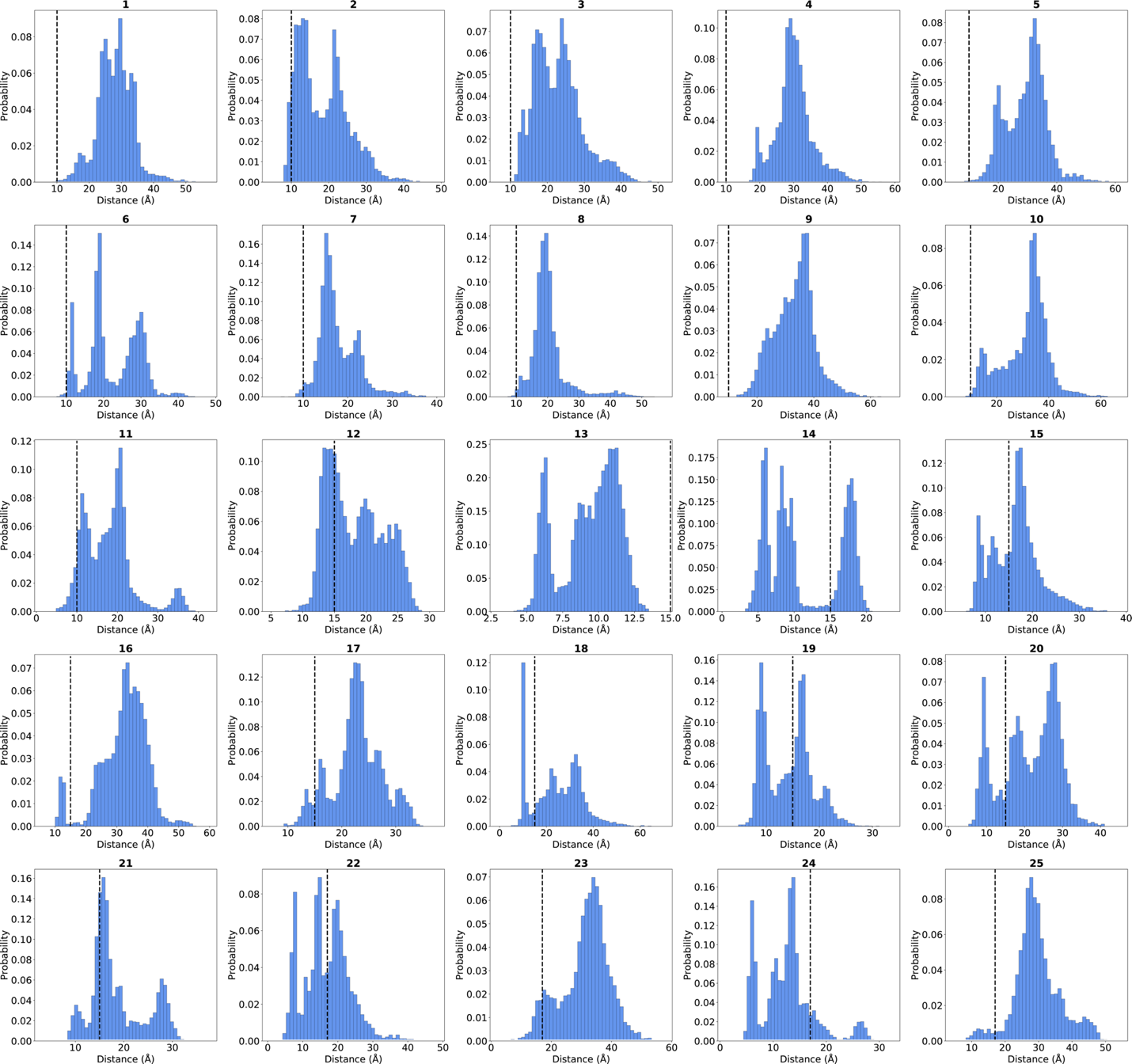

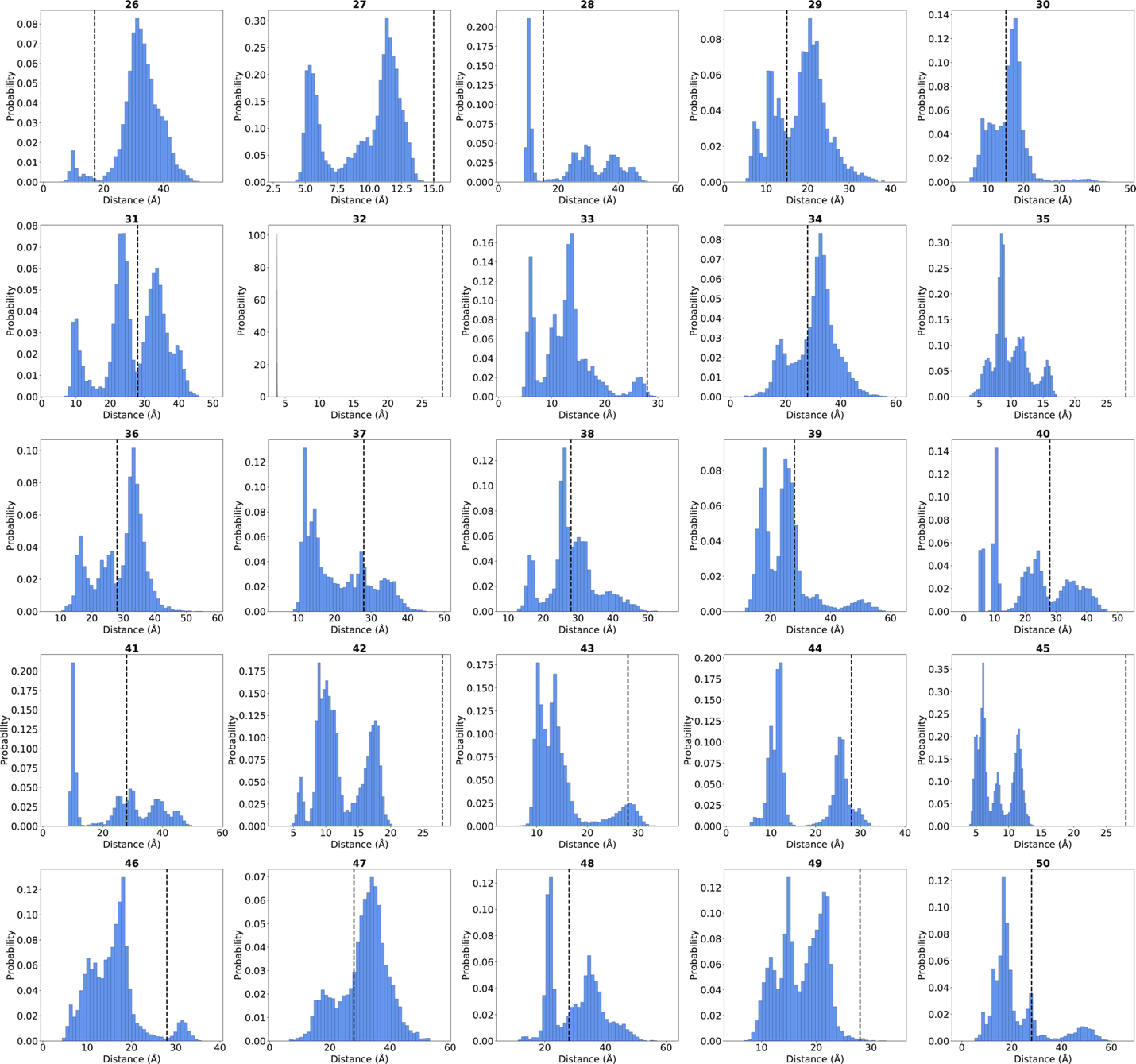

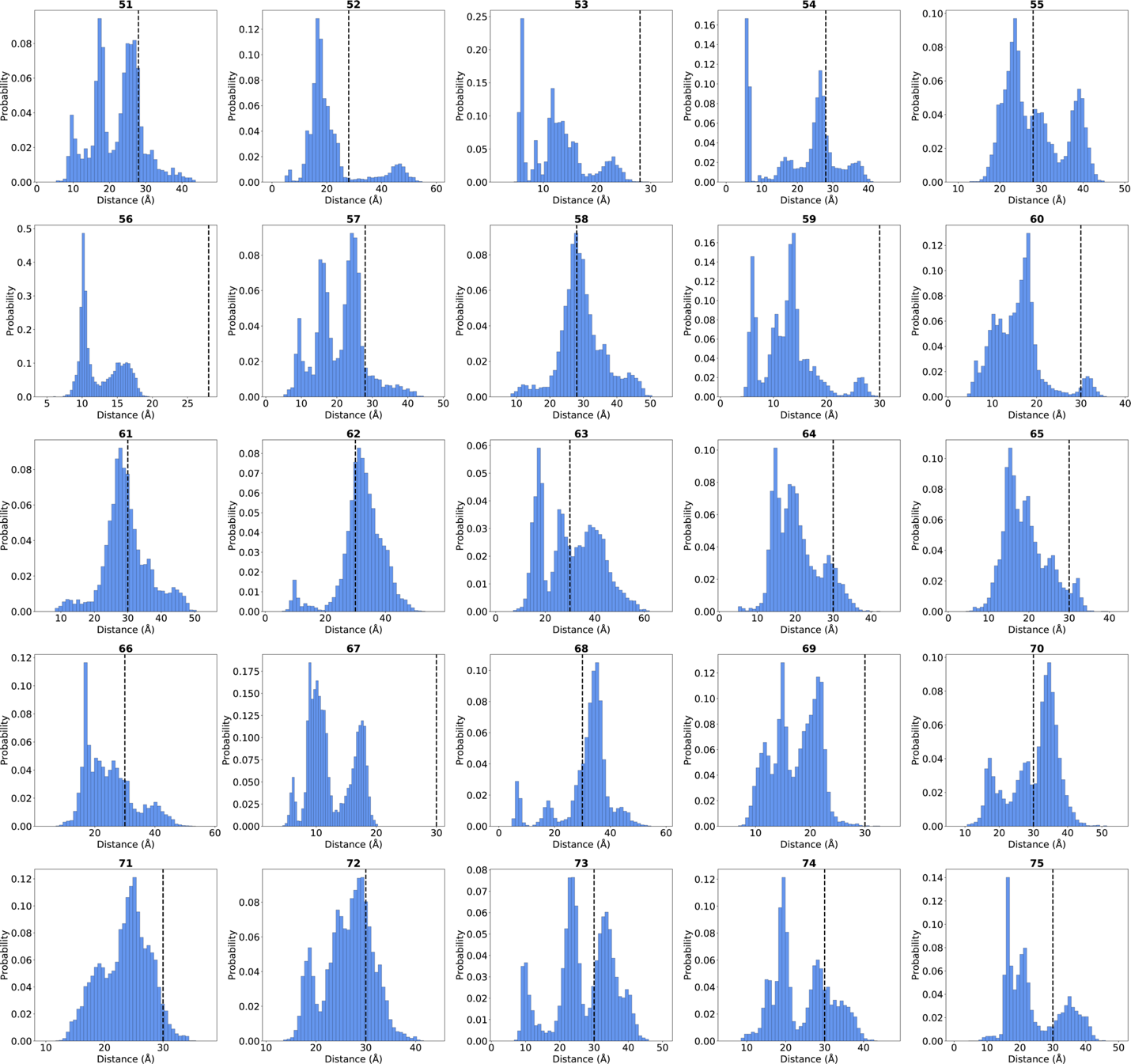

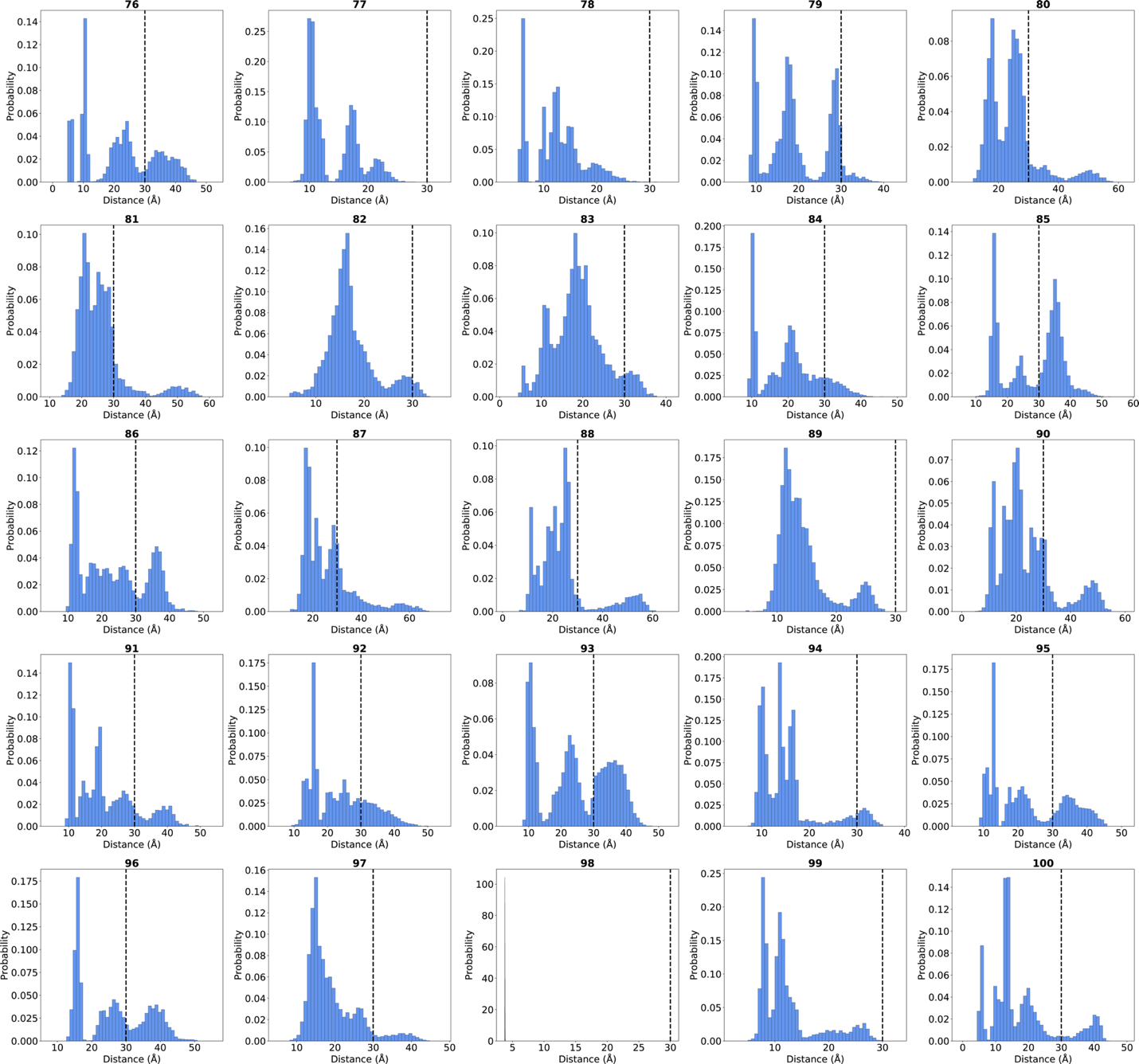

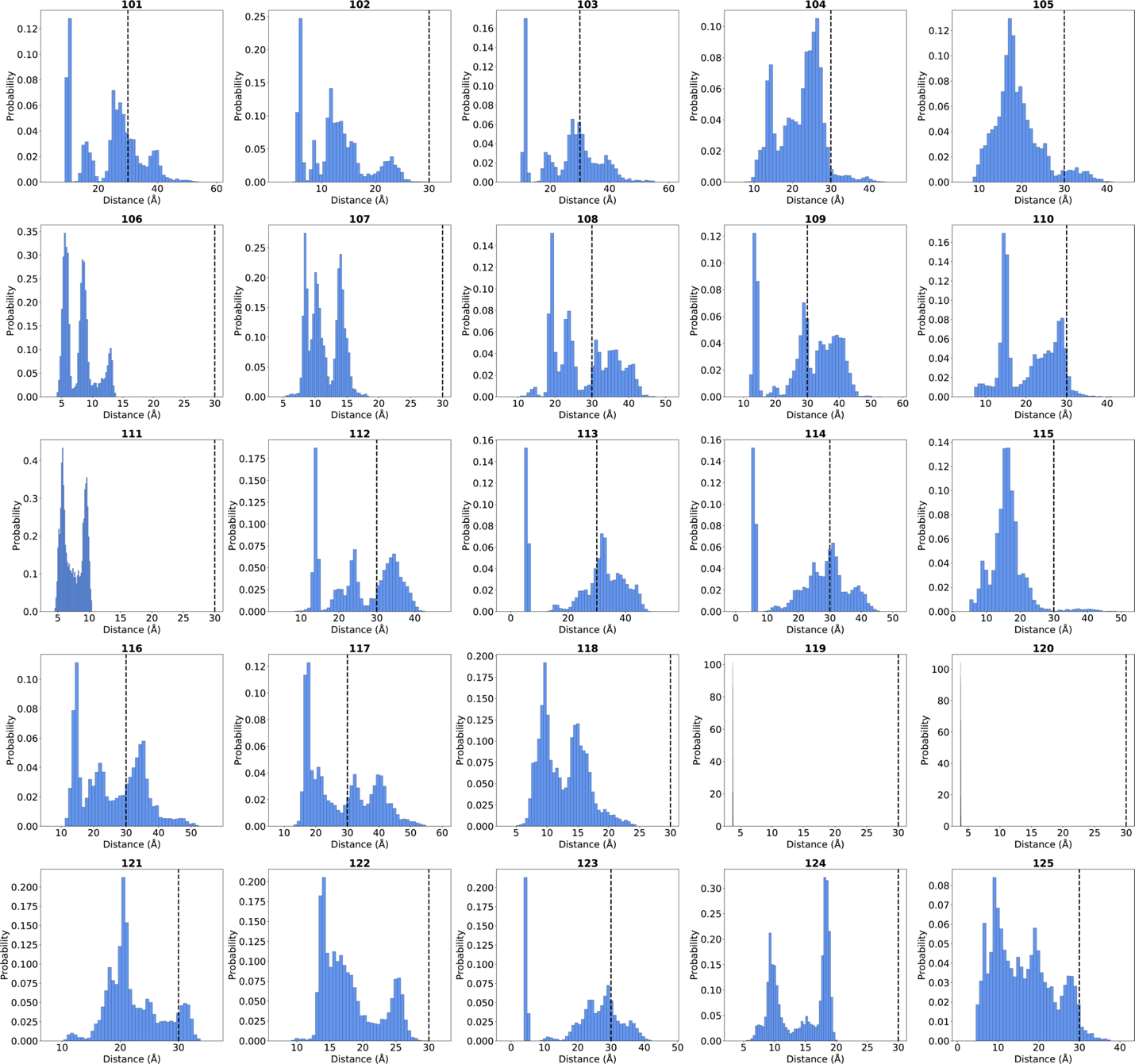

**Fig. S7. Computational distance distributions of the 125 residue pairs in Cluster 1.** The numbers above each subplot are corresponding to the numbers in Table S4 indicating each residue pair. The black dashed line indicates the maximal cross-linking length for each crosslinker: ABAS < 17 Å, EDC < 10 Å, SDA < 15 Å, TATA < 15 Å, CBDPS < 28 Å, and BS3 < 30 Å.

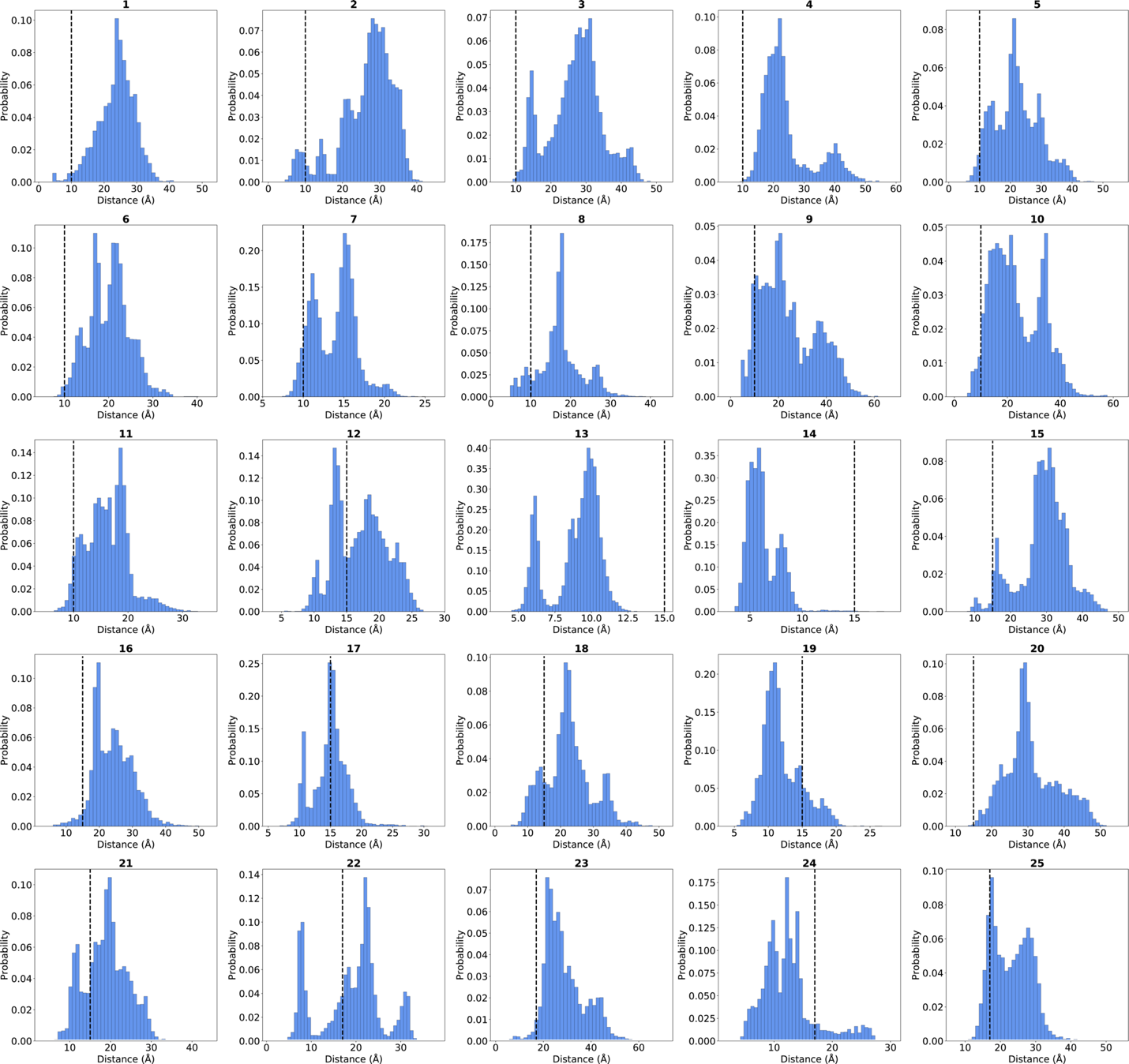

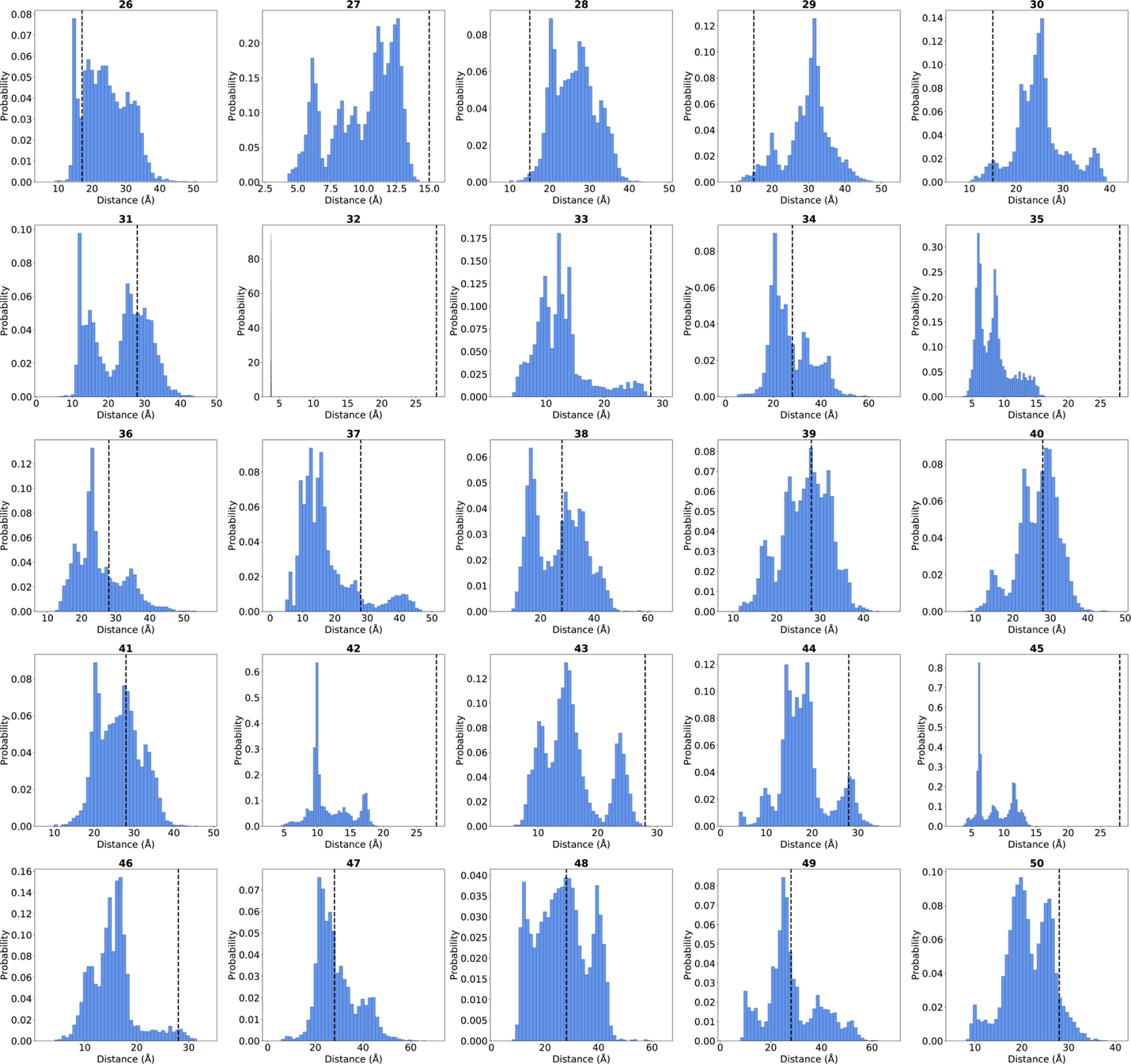

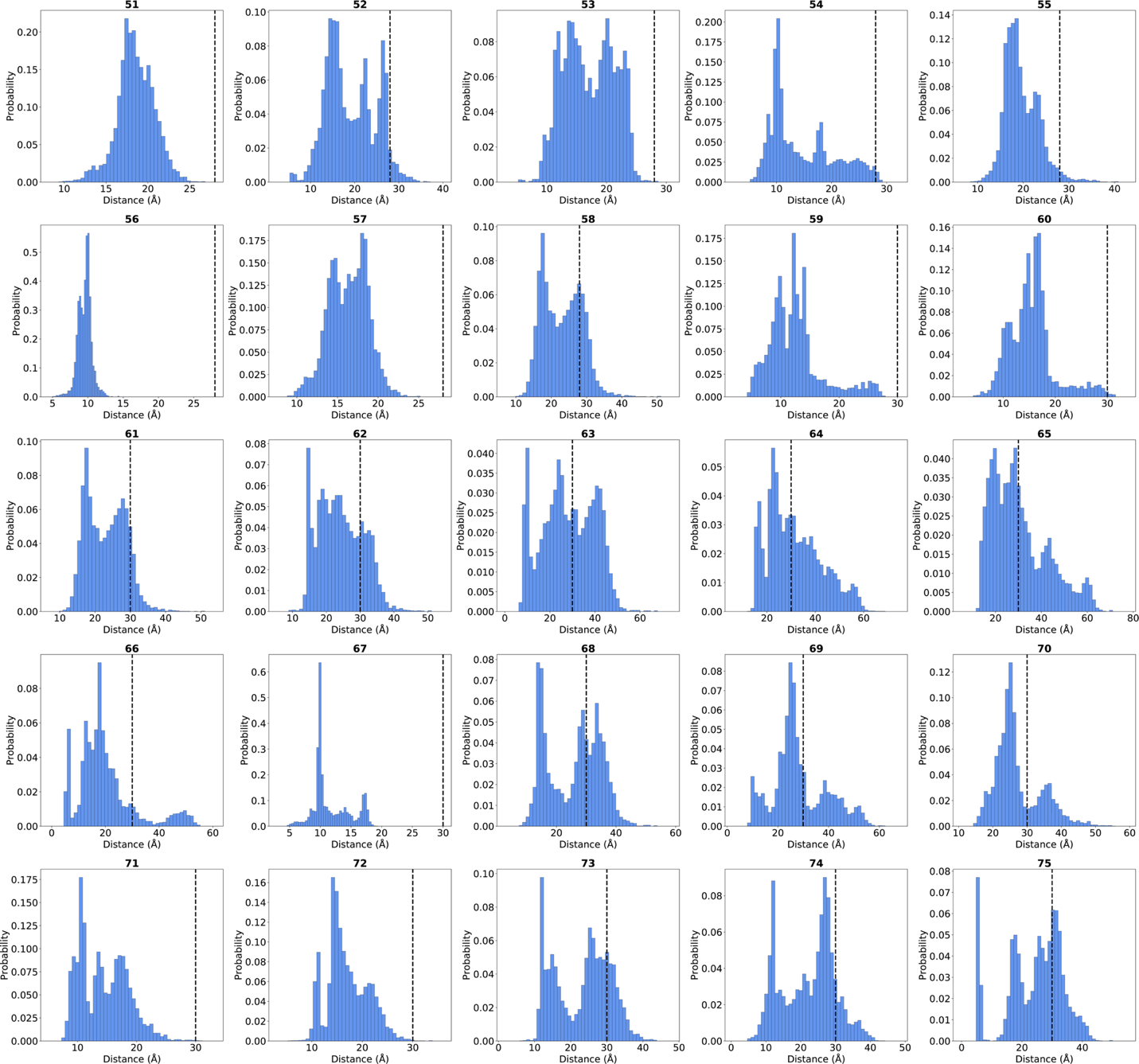

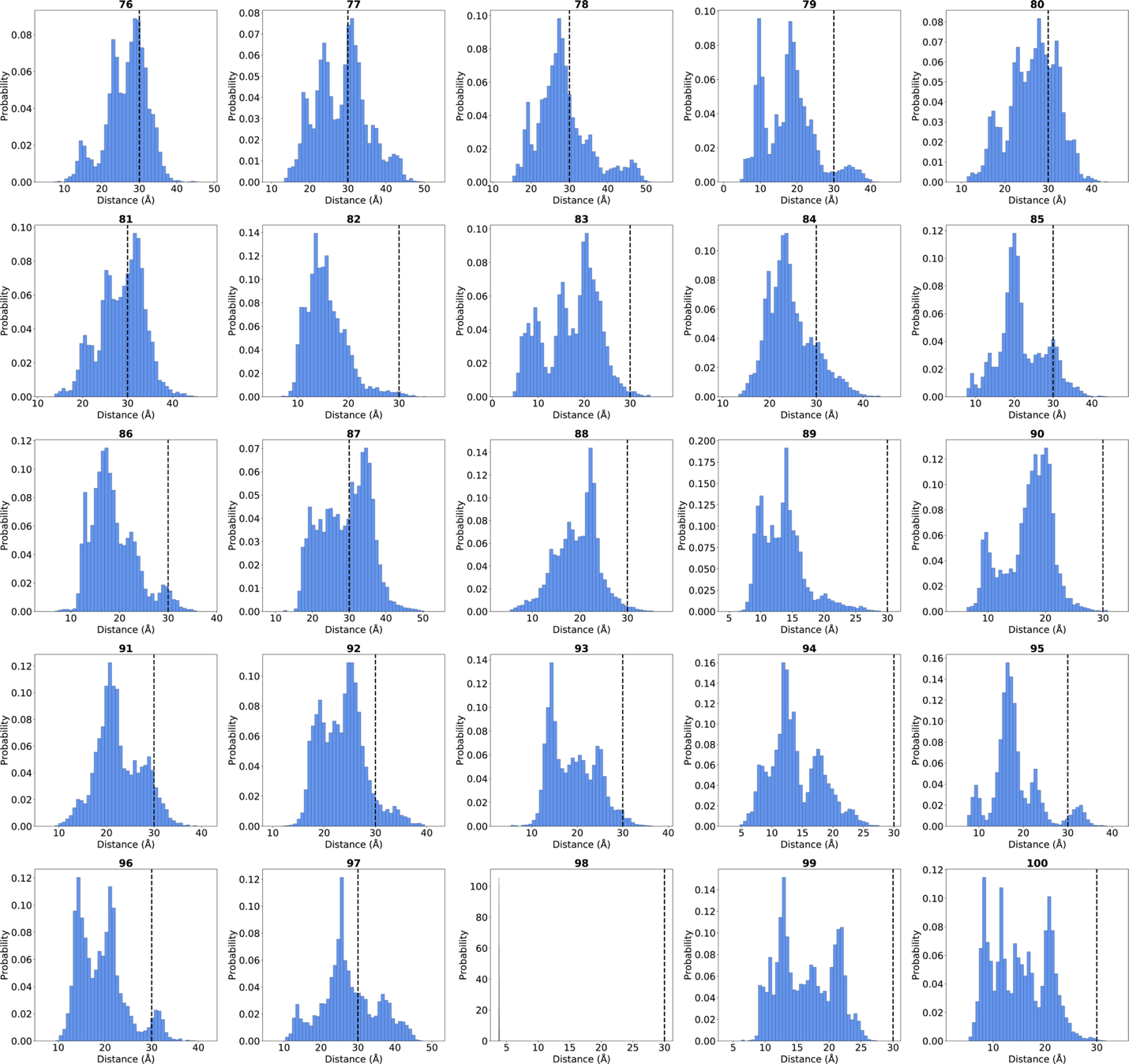

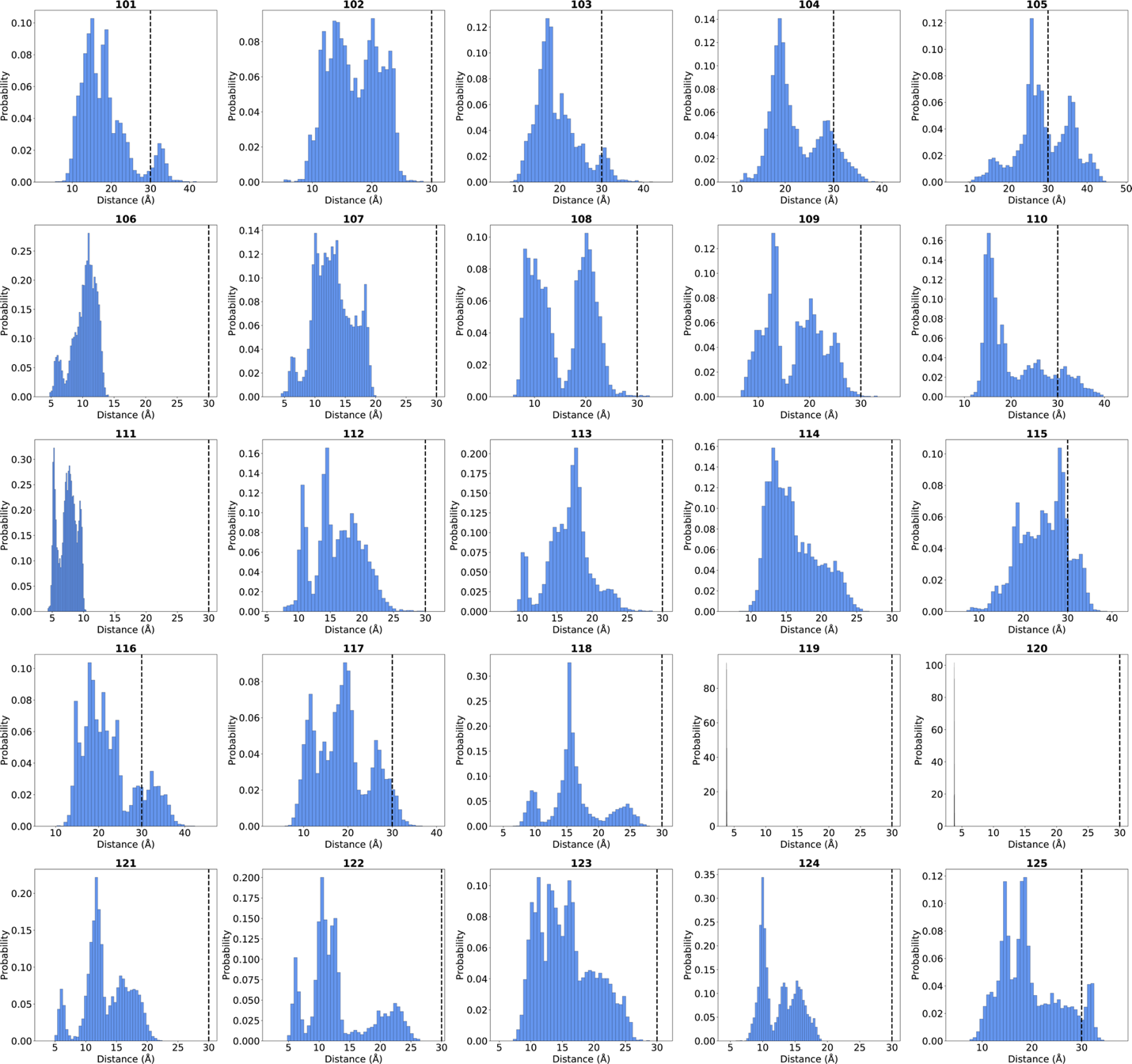

**Fig. S8. Computational distance distributions of the 125 residue pairs in Cluster 2.** The numbers above each subplot are corresponding to the numbers in Table S4 indicating each residue pair. The black dashed line indicates the maximal cross-linking length for each crosslinker: ABAS < 17 Å, EDC < 10 Å, SDA < 15 Å, TATA < 15 Å, CBDPS < 28 Å, and BS3 < 30 Å.

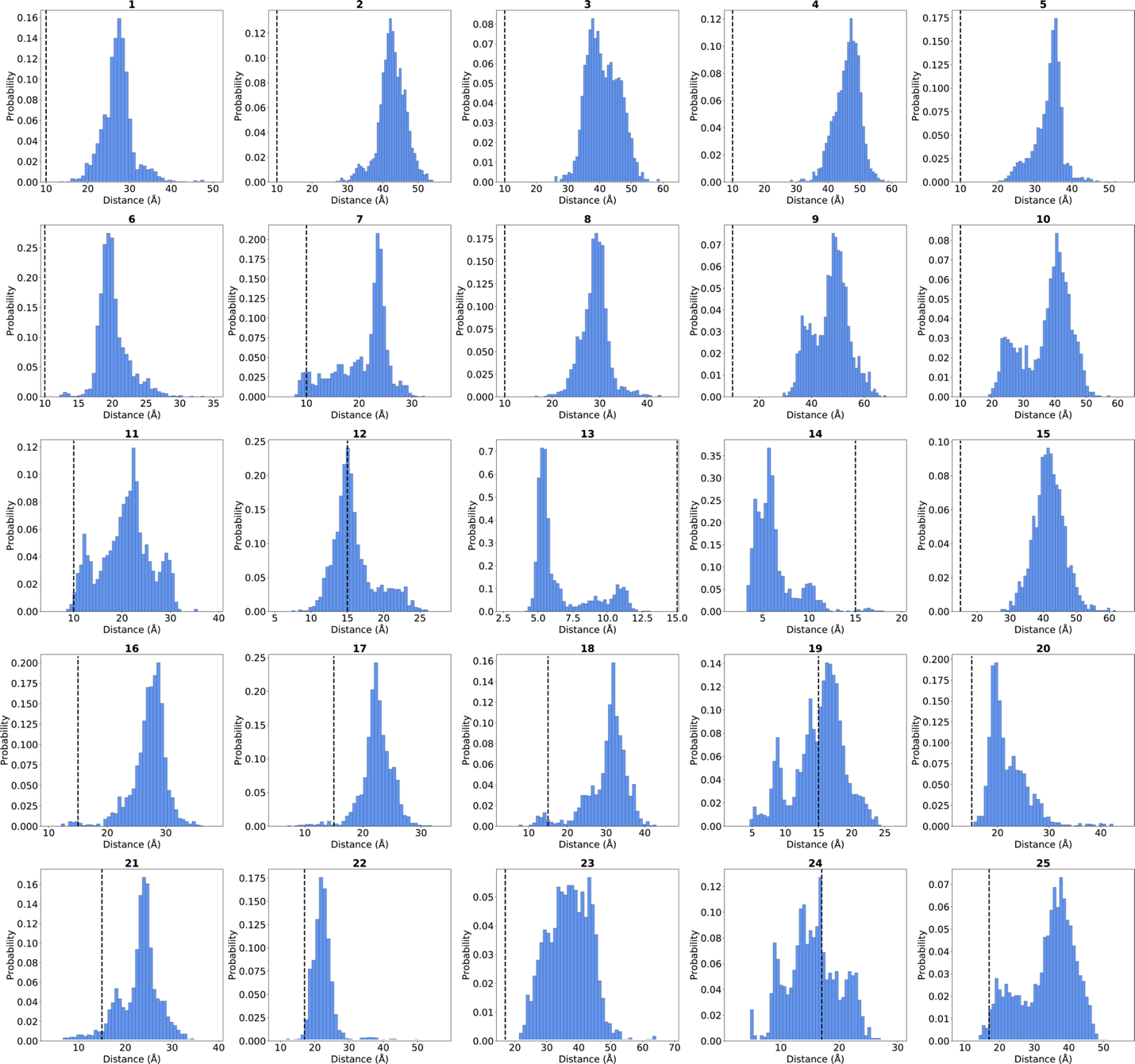

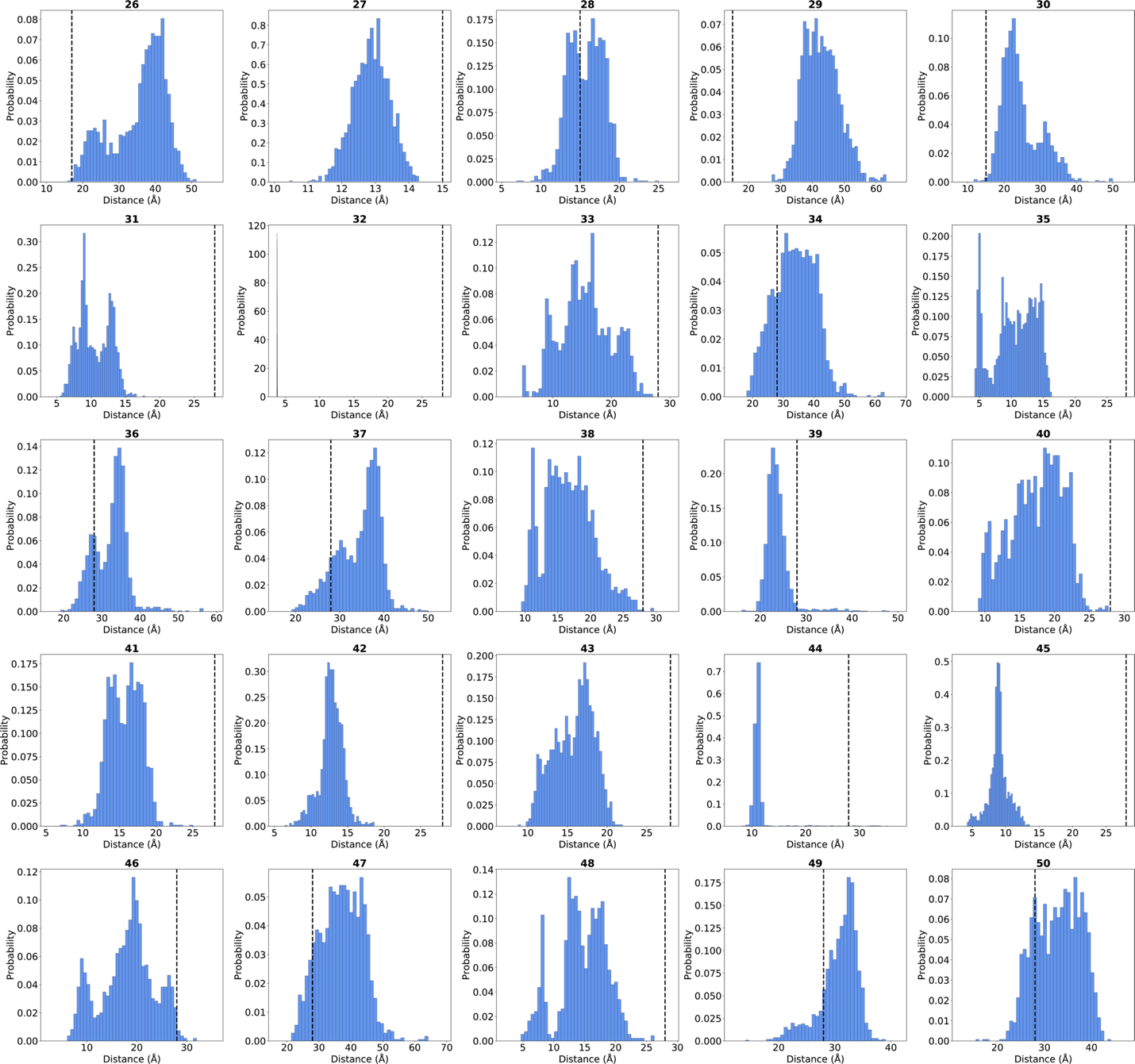

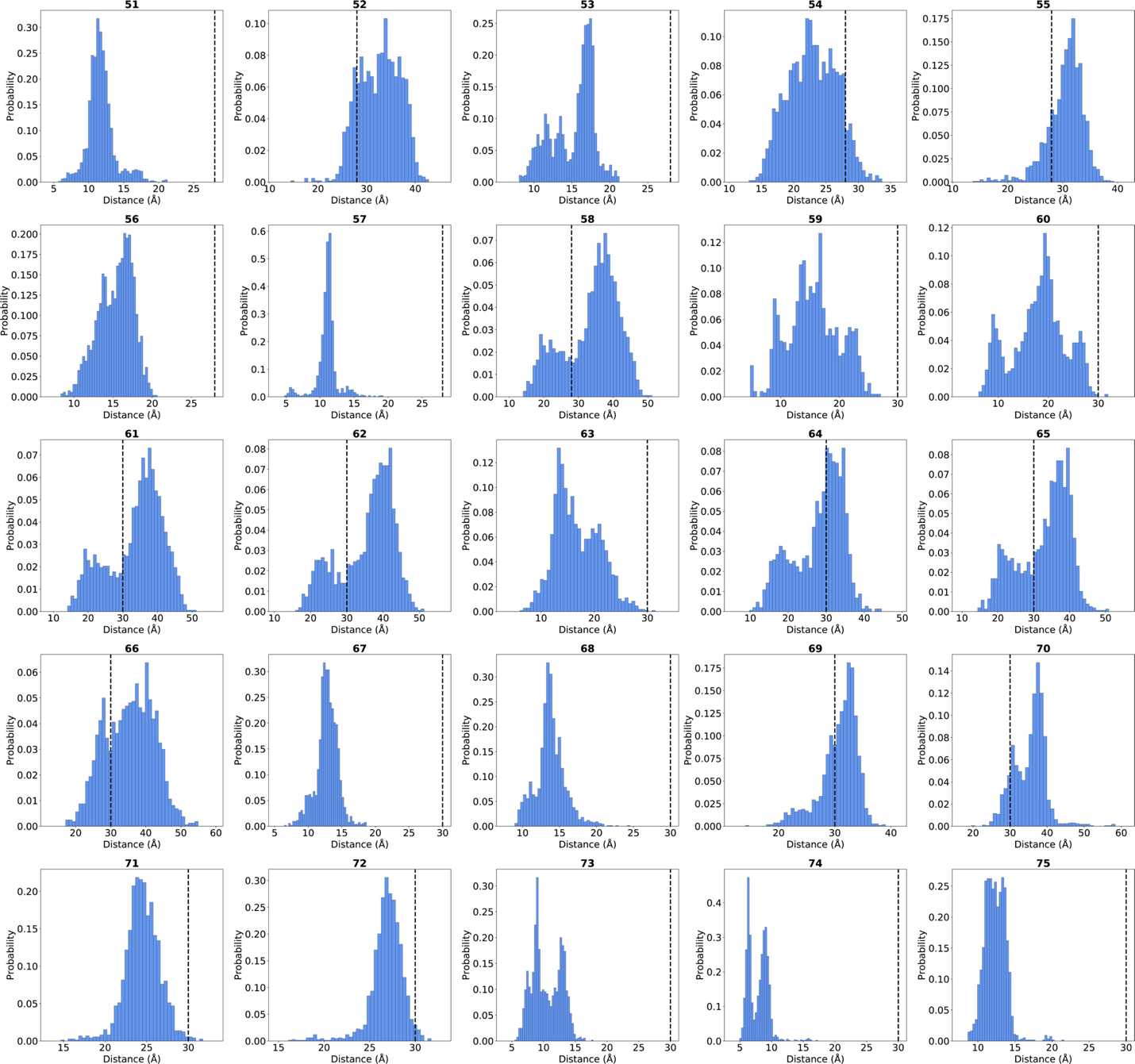

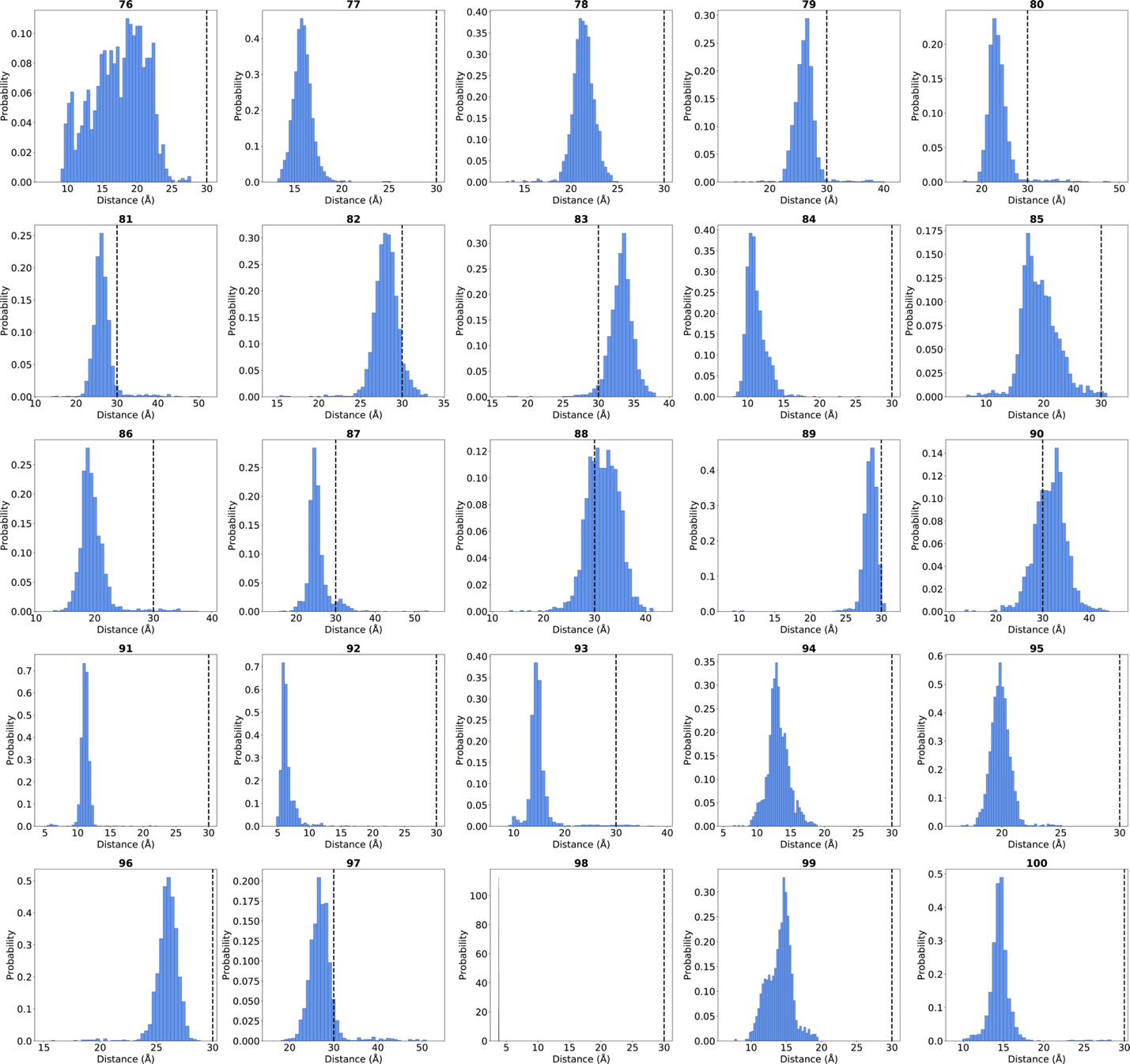

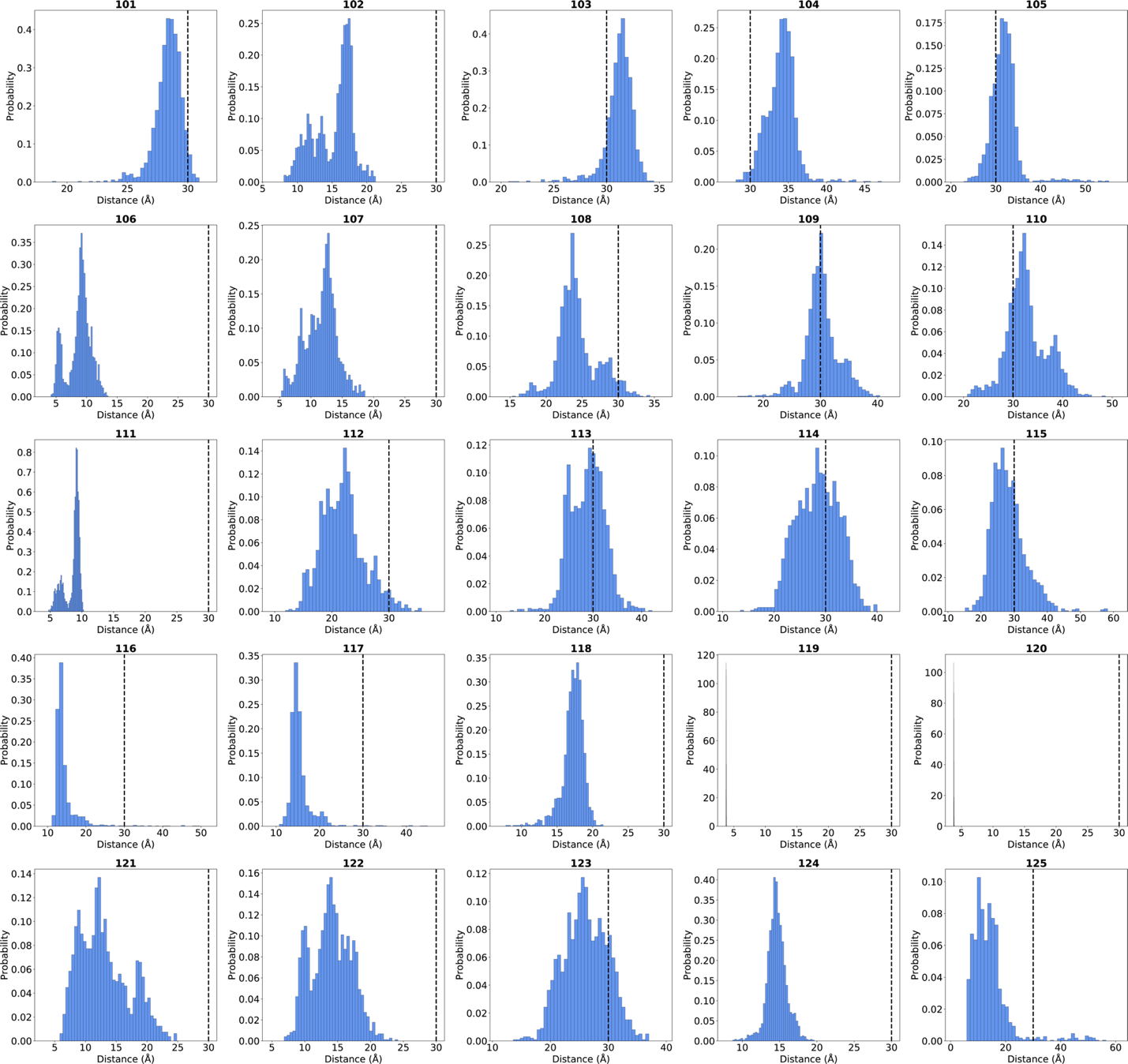

**Fig. S9. Computational distance distributions of the 125 residue pairs in Cluster 3.** The numbers above each subplot are corresponding to the numbers in Table S4 indicating each residue pair. The black dashed line indicates the maximal cross-linking length for each crosslinker: ABAS < 17 Å, EDC < 10 Å, SDA < 15 Å, TATA < 15 Å, CBDPS < 28 Å, and BS3 < 30 Å.

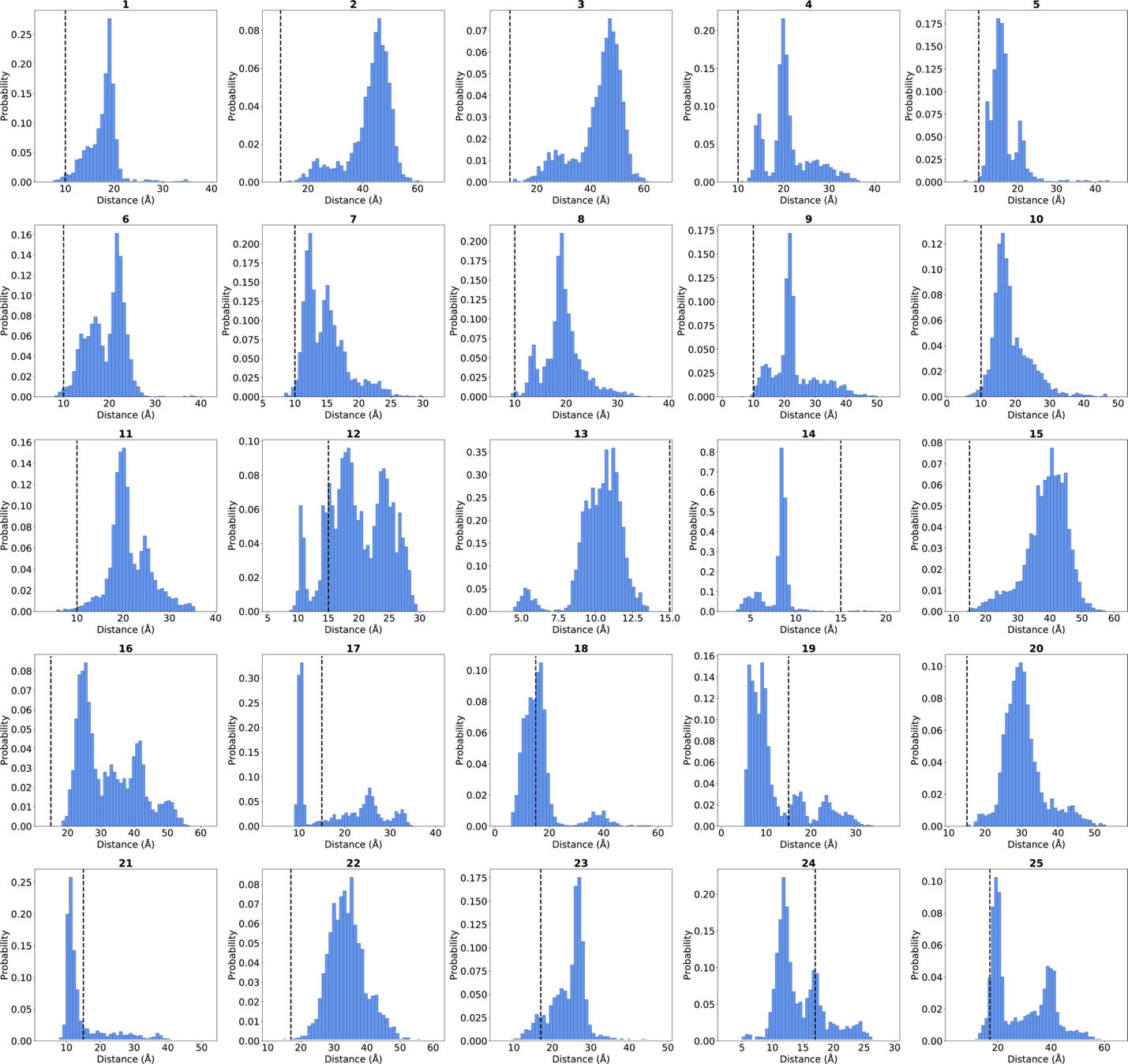

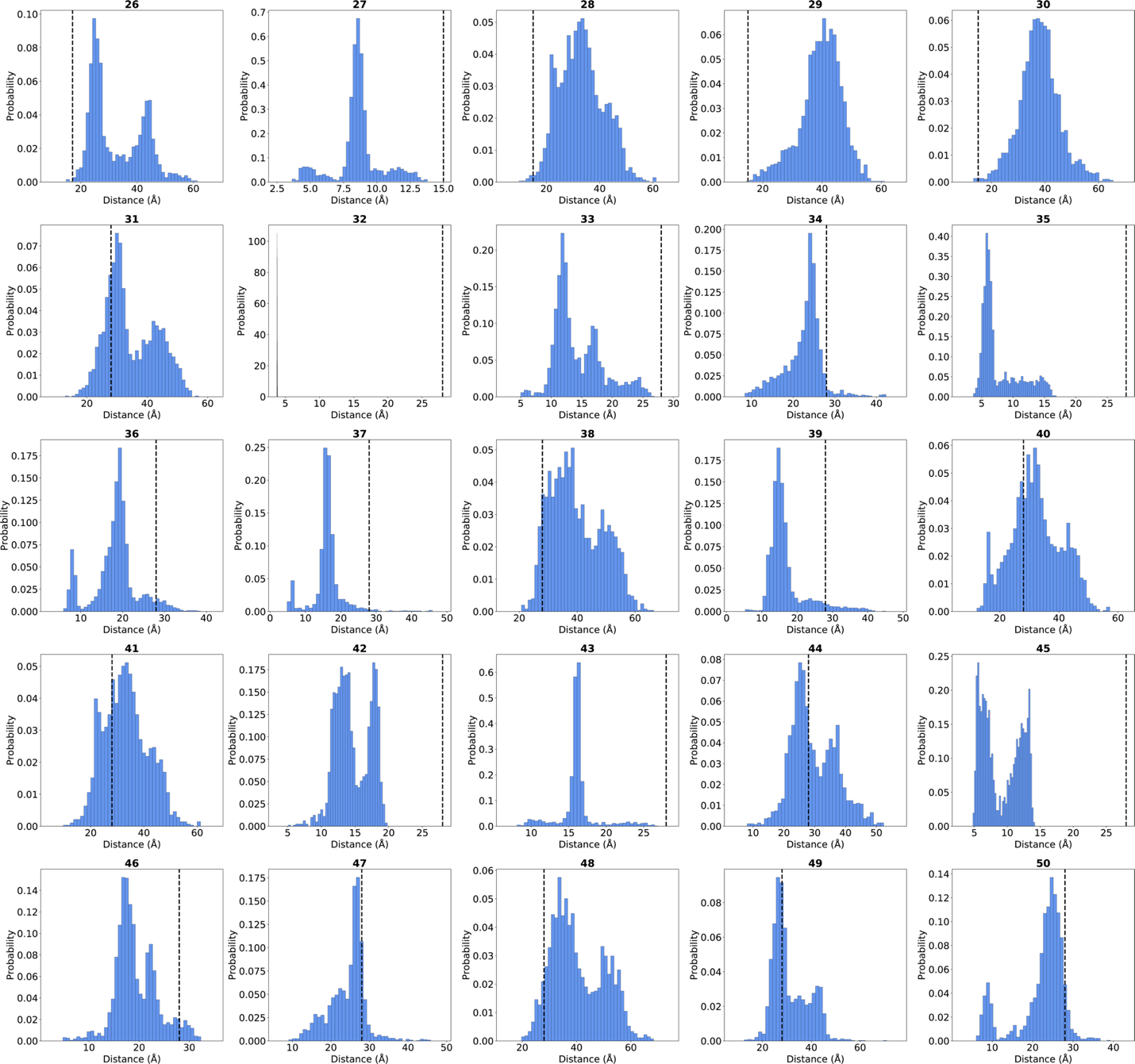

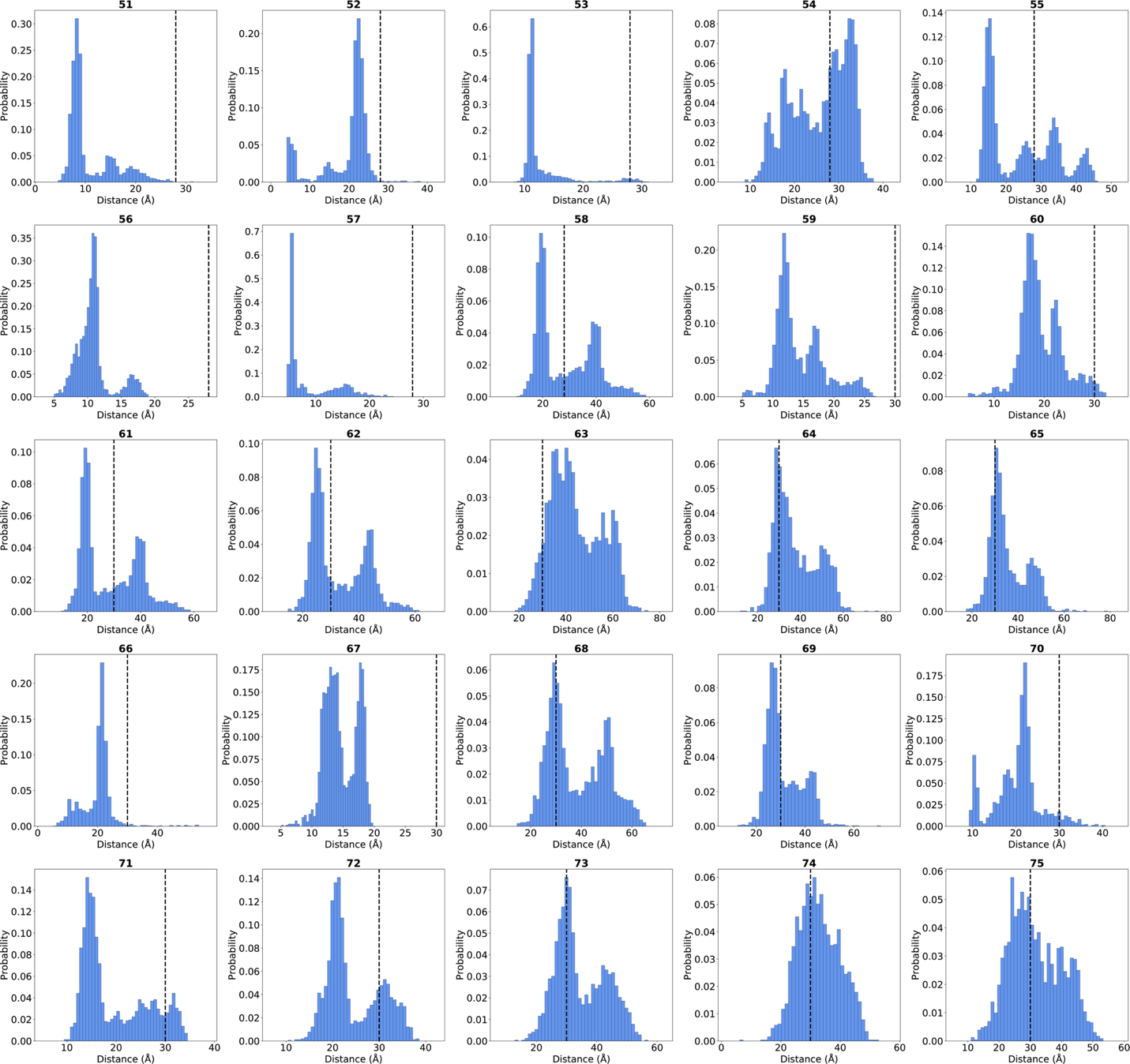

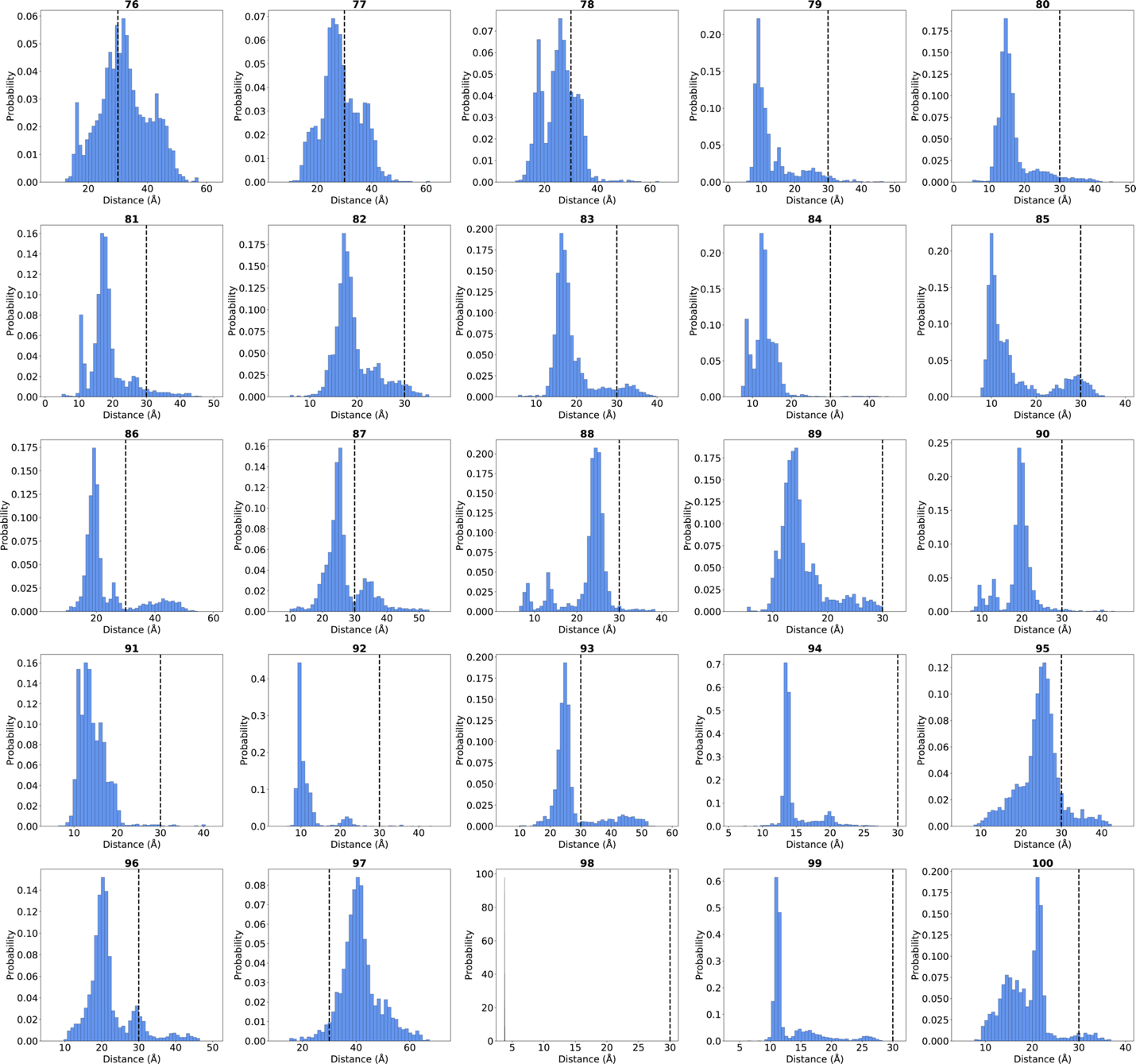

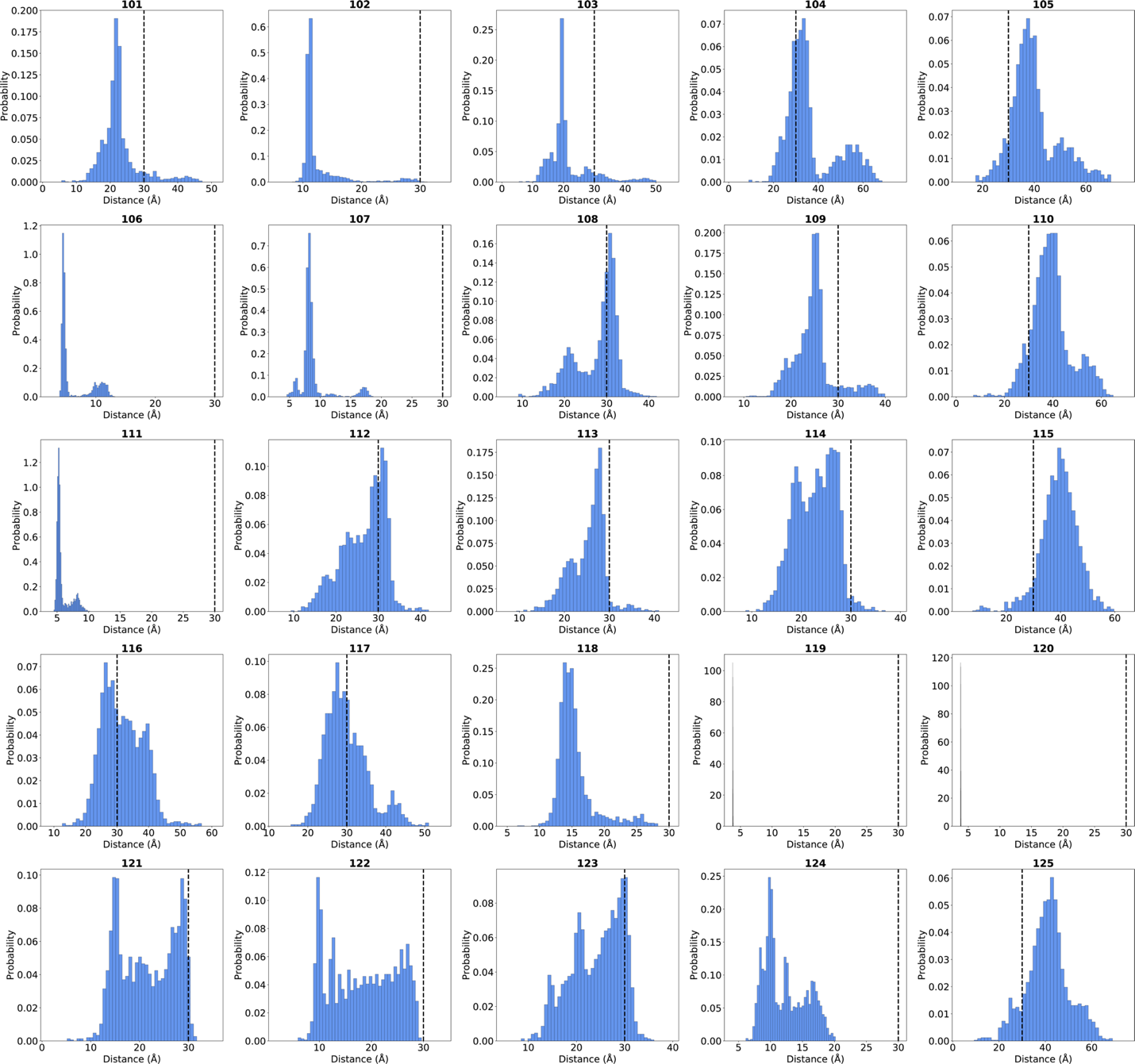

**Fig. S10. Computational distance distributions of the 125 residue pairs in Cluster 4.** The numbers above each subplot are corresponding to the numbers in Table S4 indicating each residue pair. The black dashed line indicates the maximal cross-linking length for each crosslinker: ABAS < 17 Å, EDC < 10 Å, SDA < 15 Å, TATA < 15 Å, CBDPS < 28 Å, and BS3 < 30 Å.

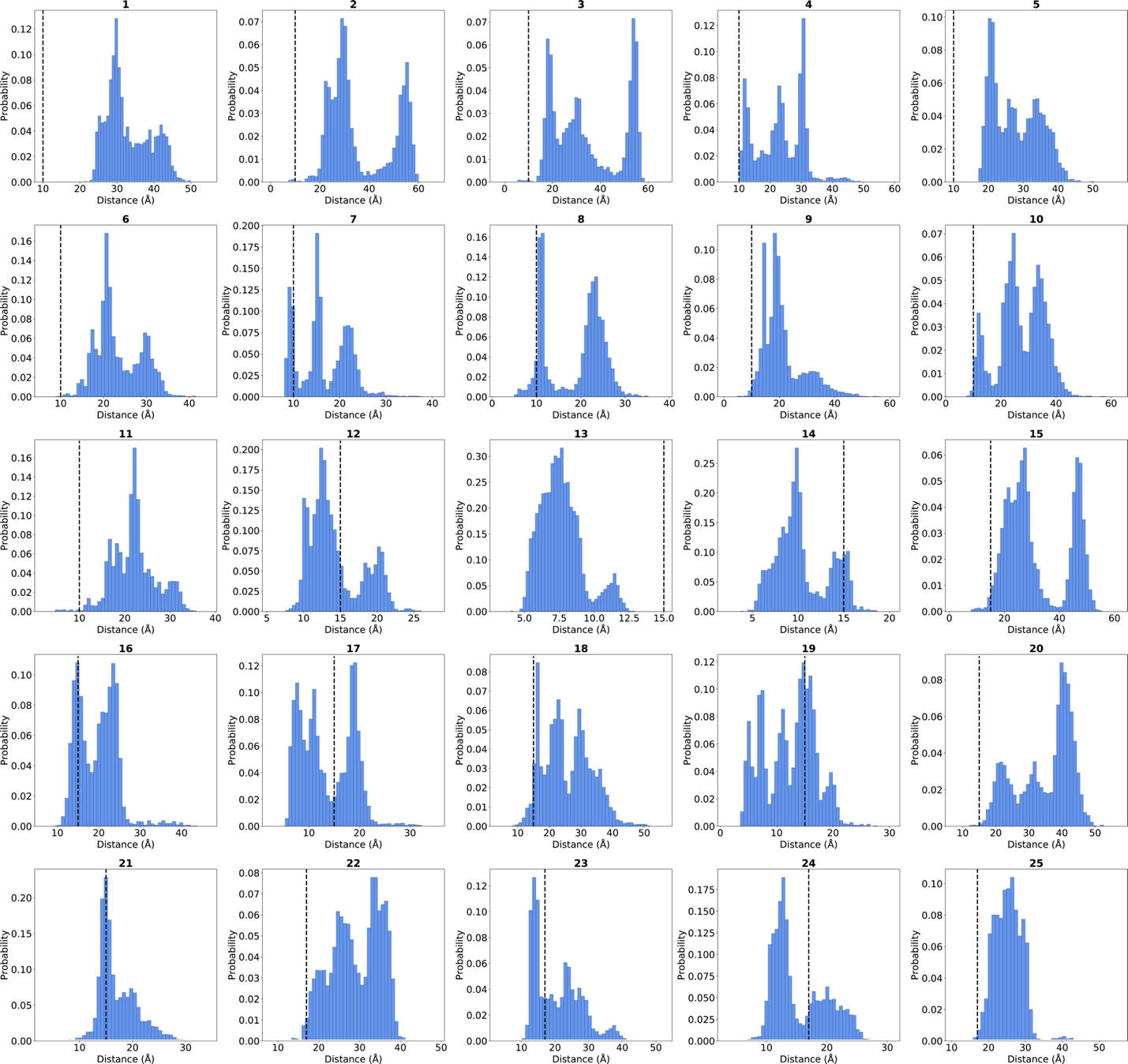

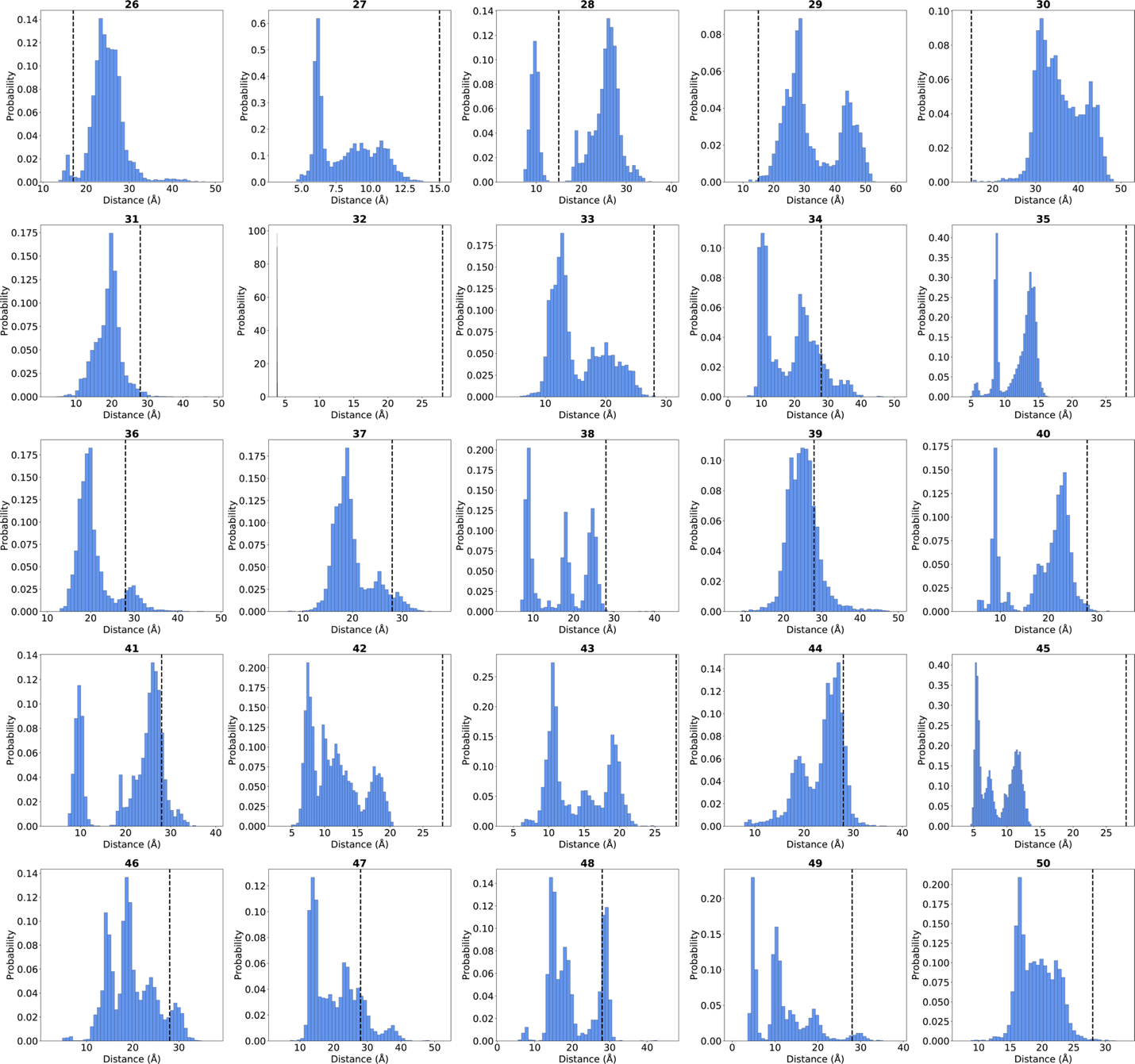

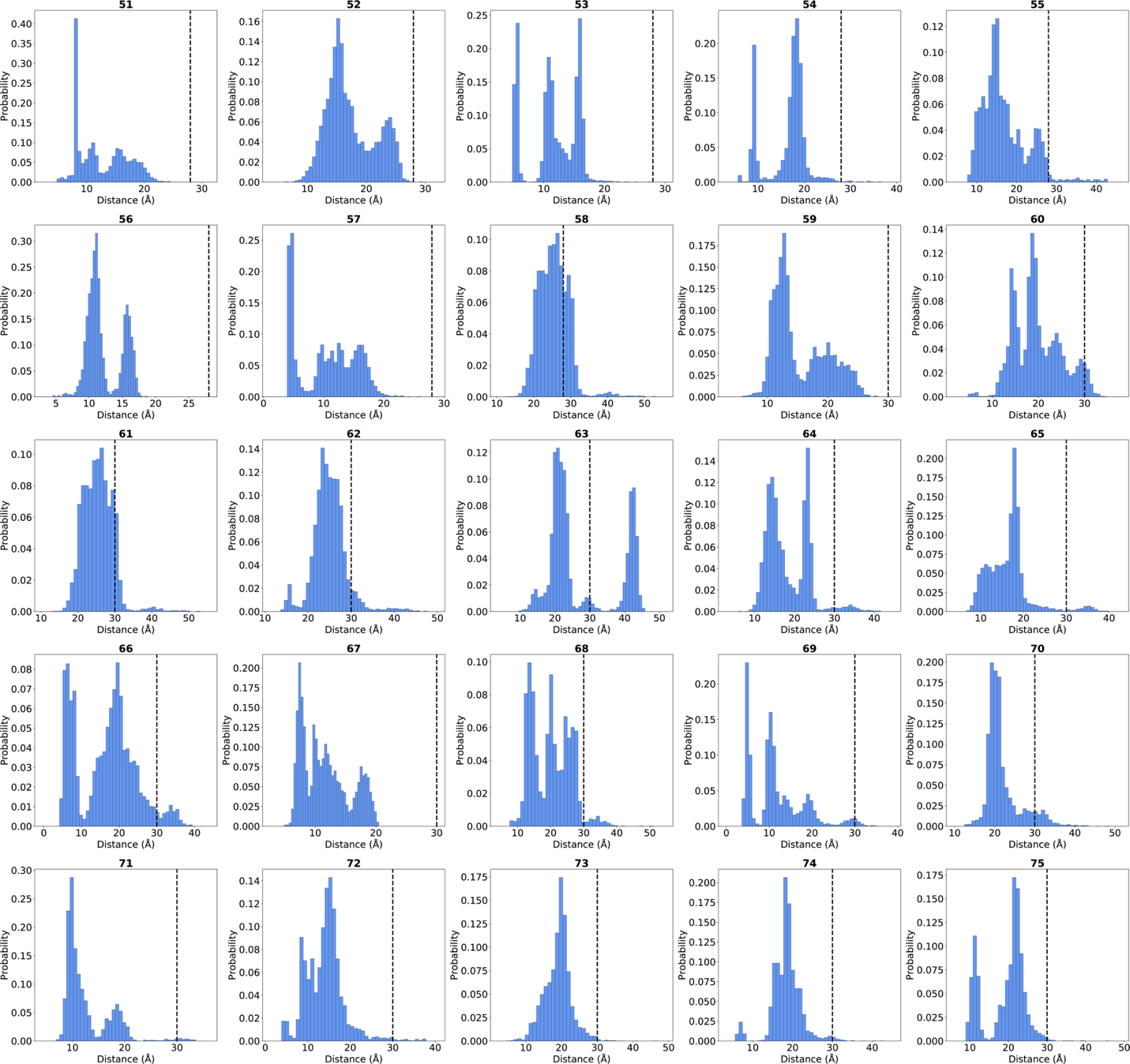

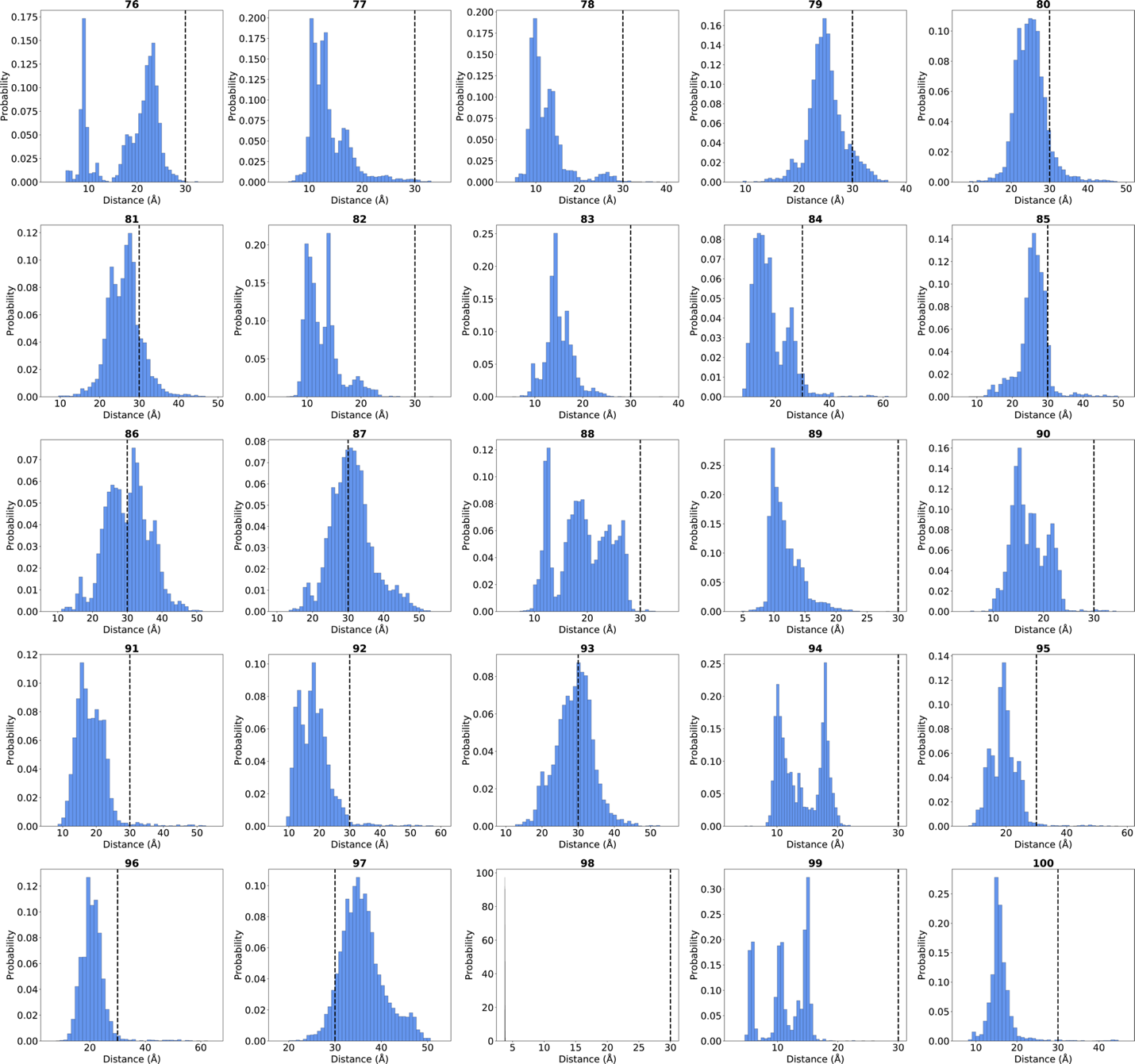

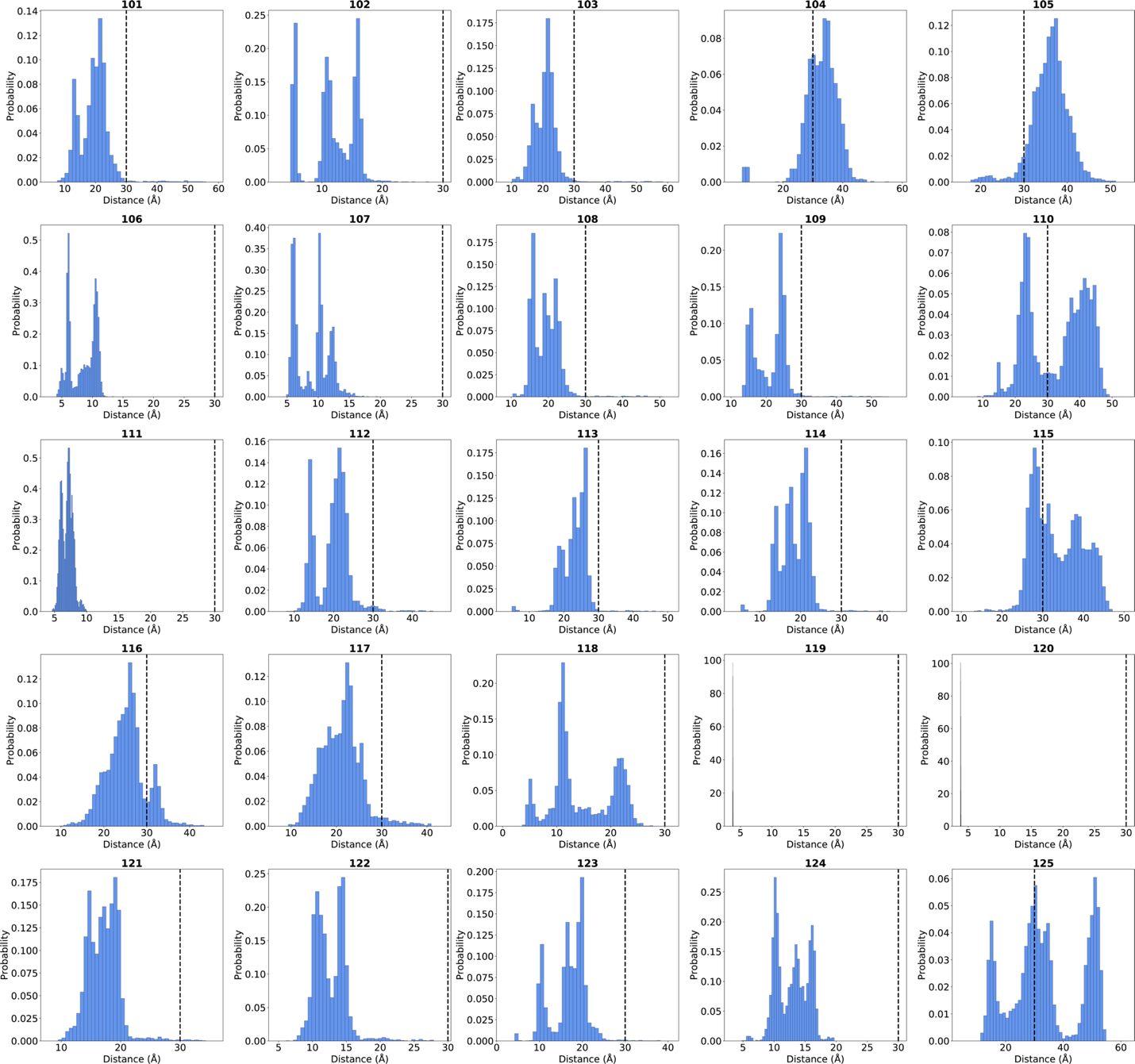

**Fig. S11. Computational distance distributions of the 125 residue pairs in Cluster 5.** The numbers above each subplot are corresponding to the numbers in Table S4 indicating each residue pair. The black dashed line indicates the maximal cross-linking length for each crosslinker: ABAS < 17 Å, EDC < 10 Å, SDA < 15 Å, TATA < 15 Å, CBDPS < 28 Å, and BS3 < 30 Å.

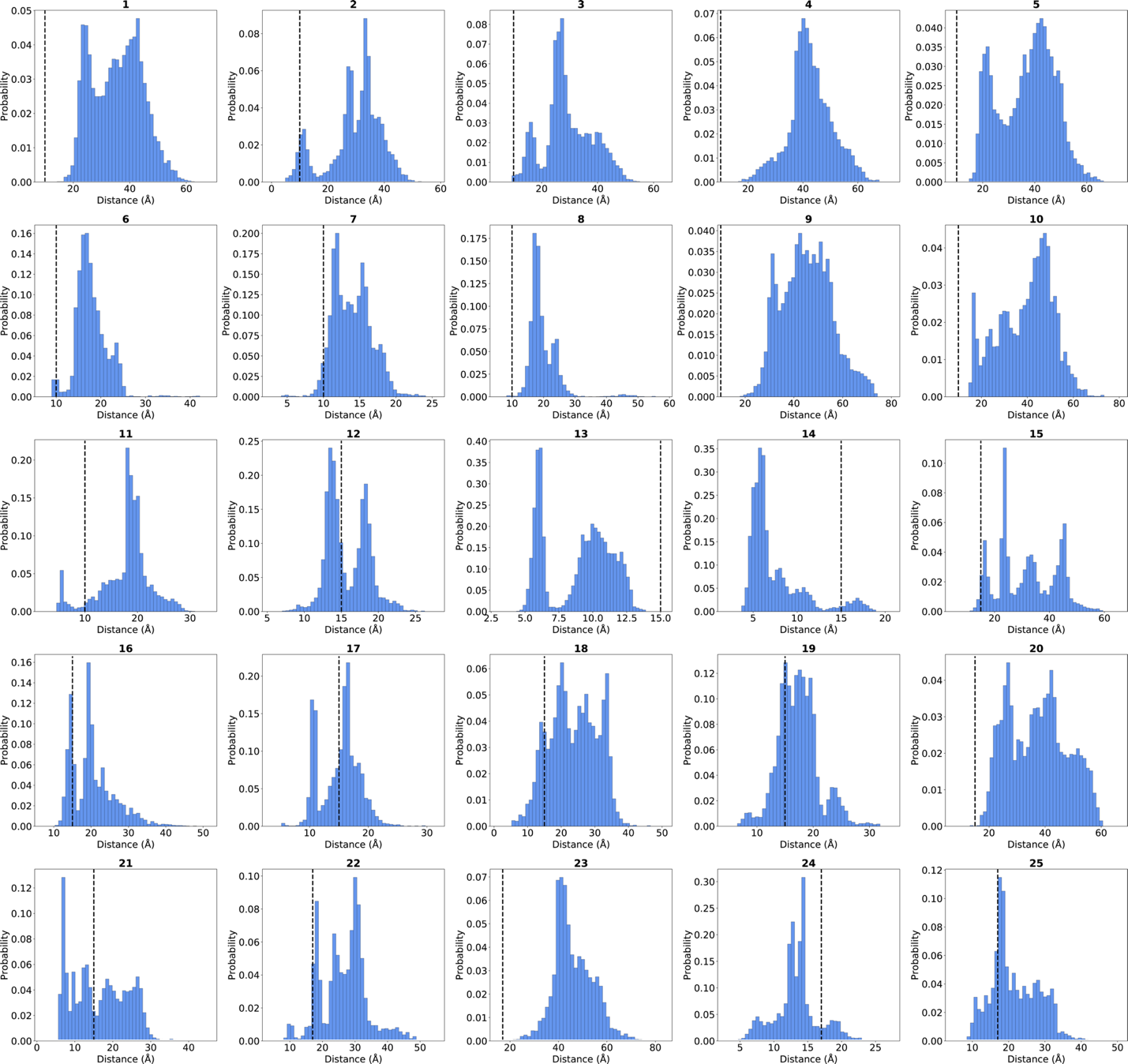

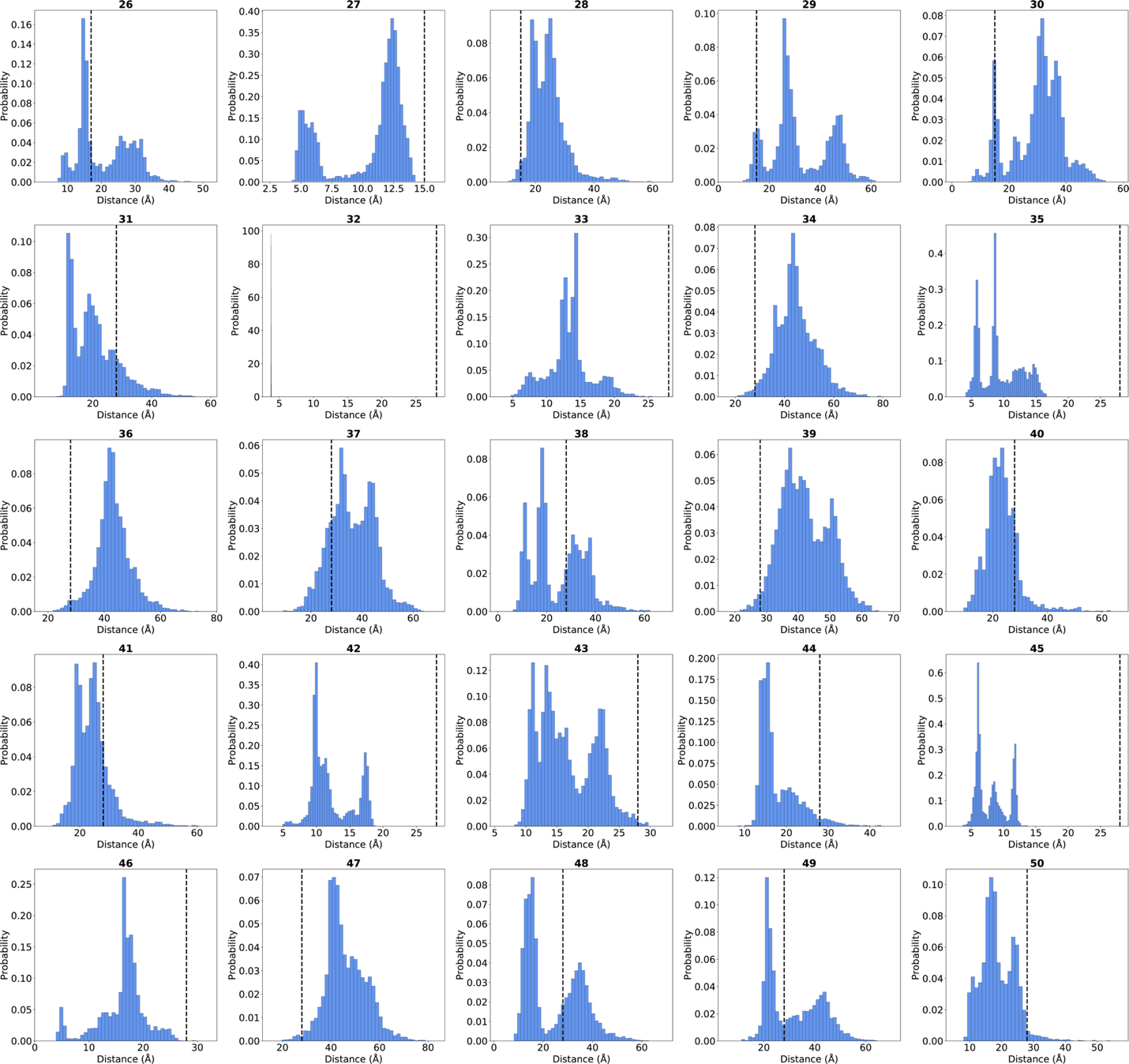

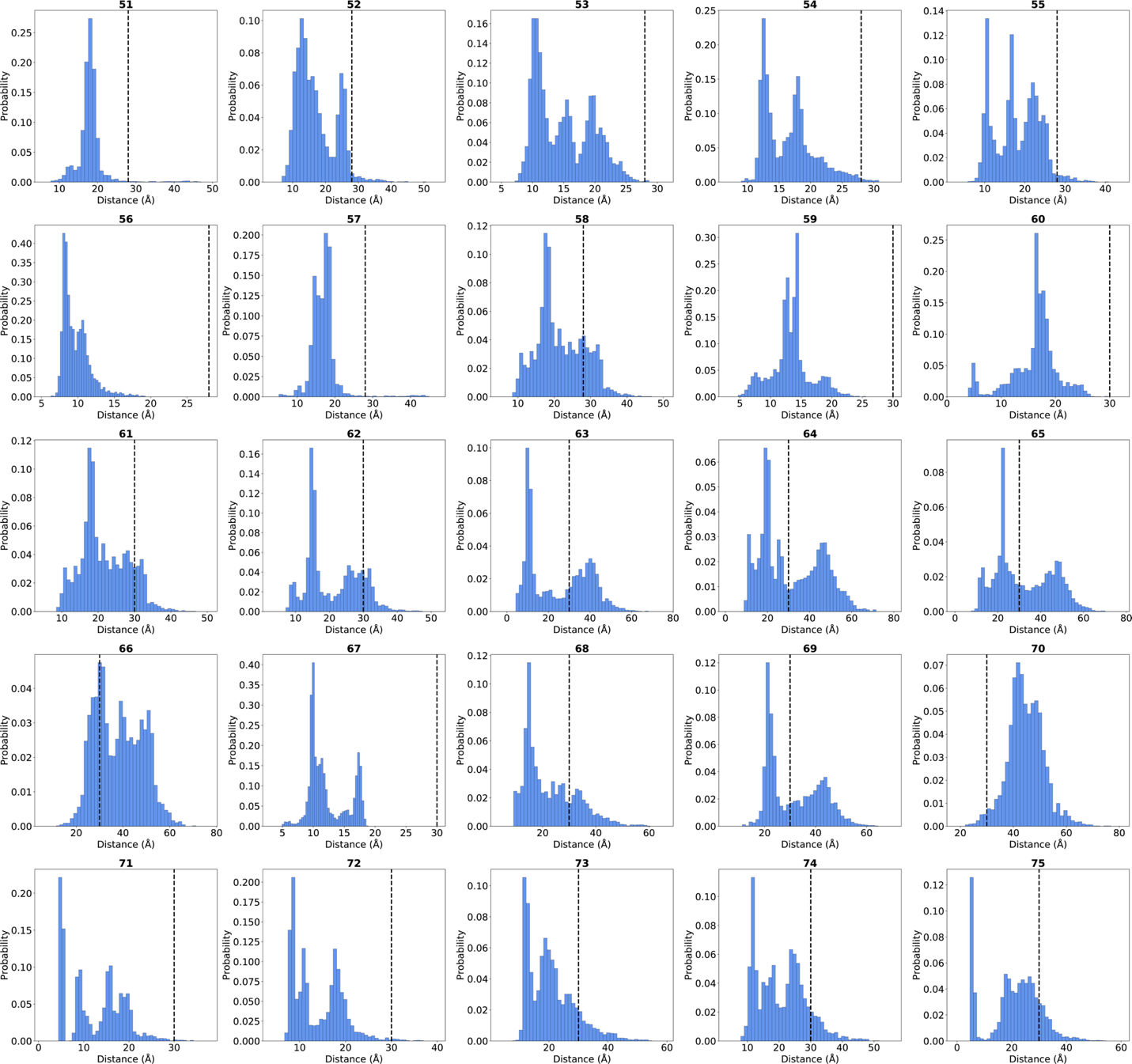

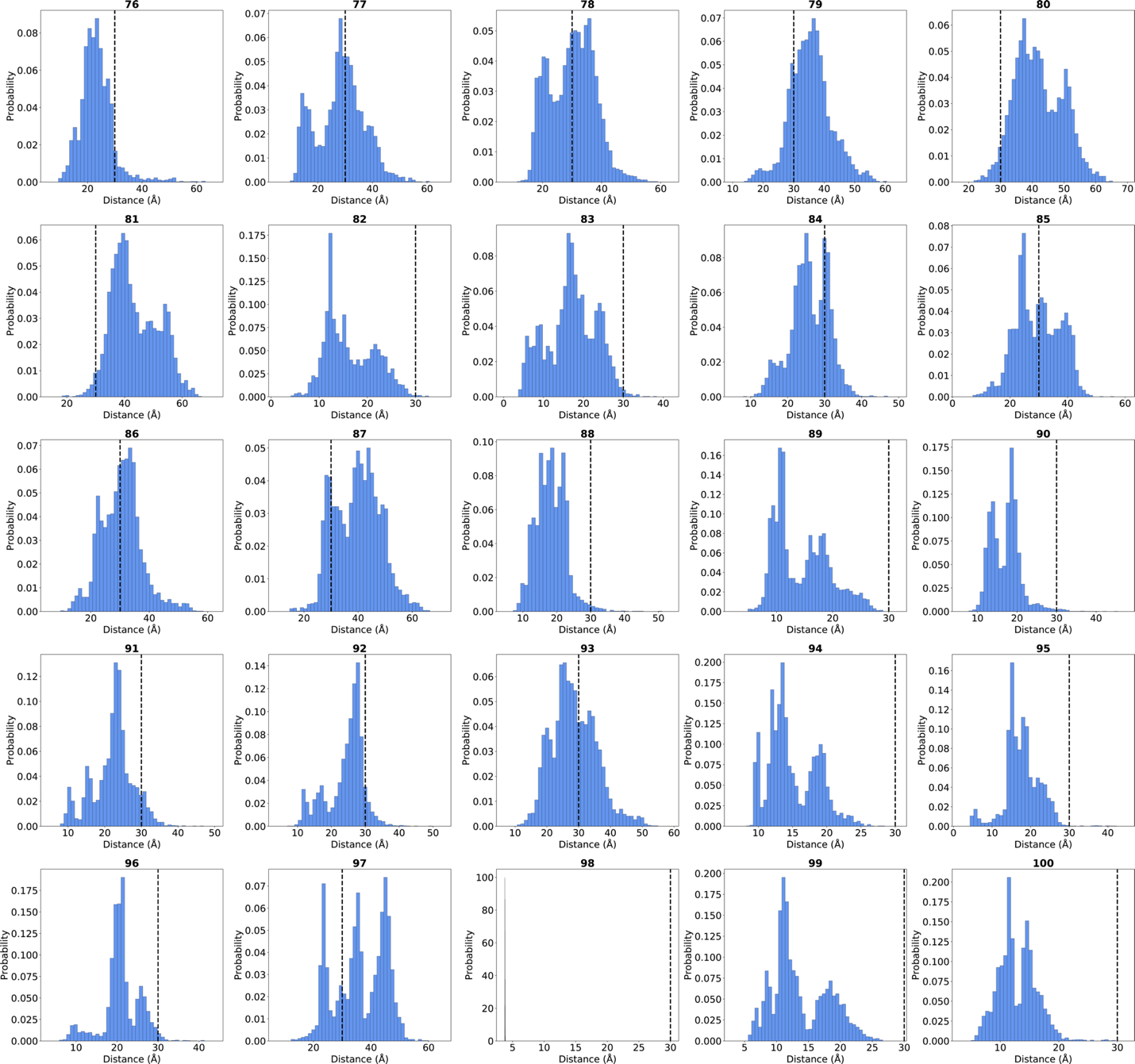

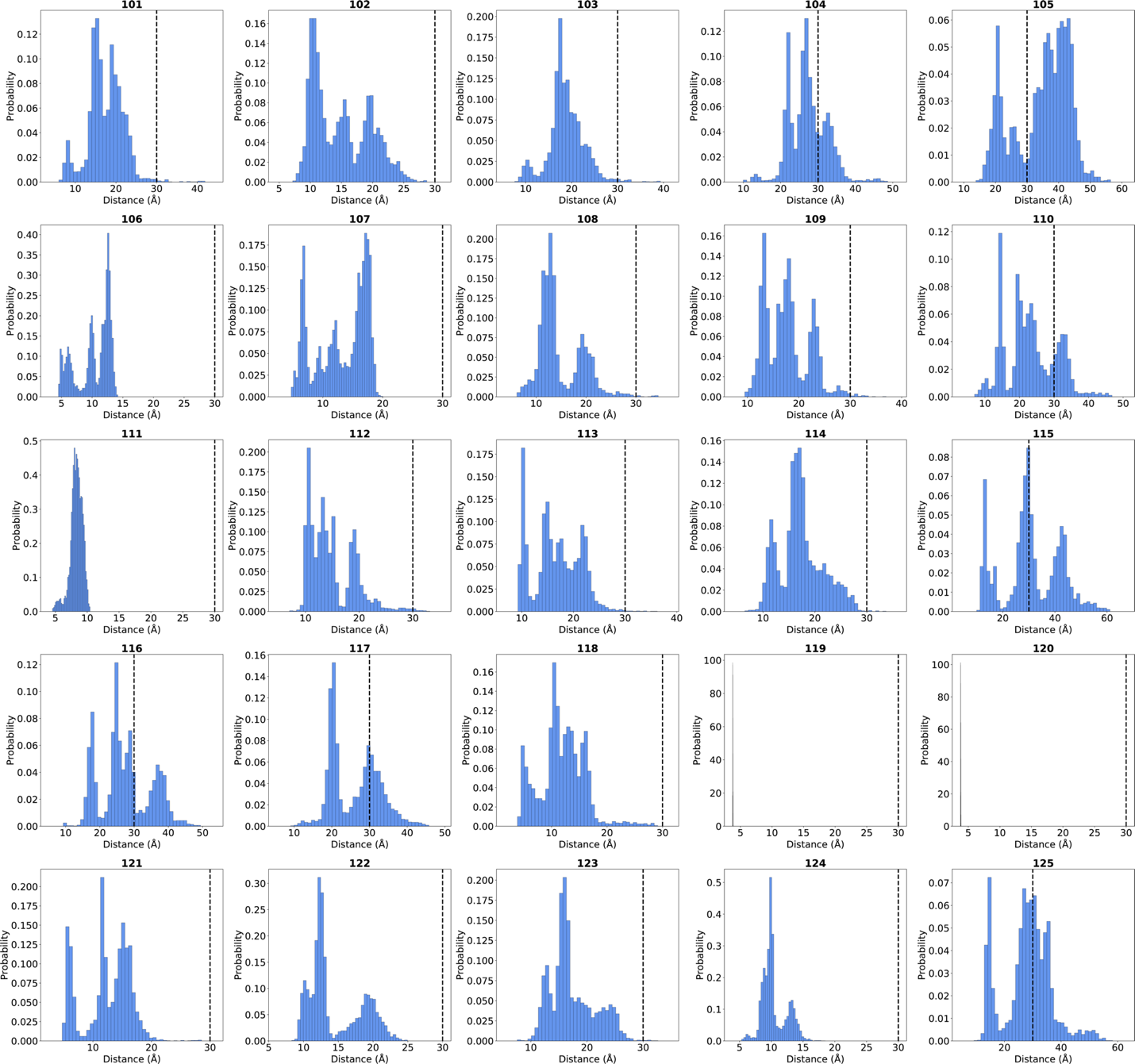

**Fig. S12. Computational distance distributions of the 125 residue pairs in Cluster 6.** The numbers above each subplot are corresponding to the numbers in Table S4 indicating each residue pair. The black dashed line indicates the maximal cross-linking length for each crosslinker: ABAS < 17 Å, EDC < 10 Å, SDA < 15 Å, TATA < 15 Å, CBDPS < 28 Å, and BS3 < 30 Å.

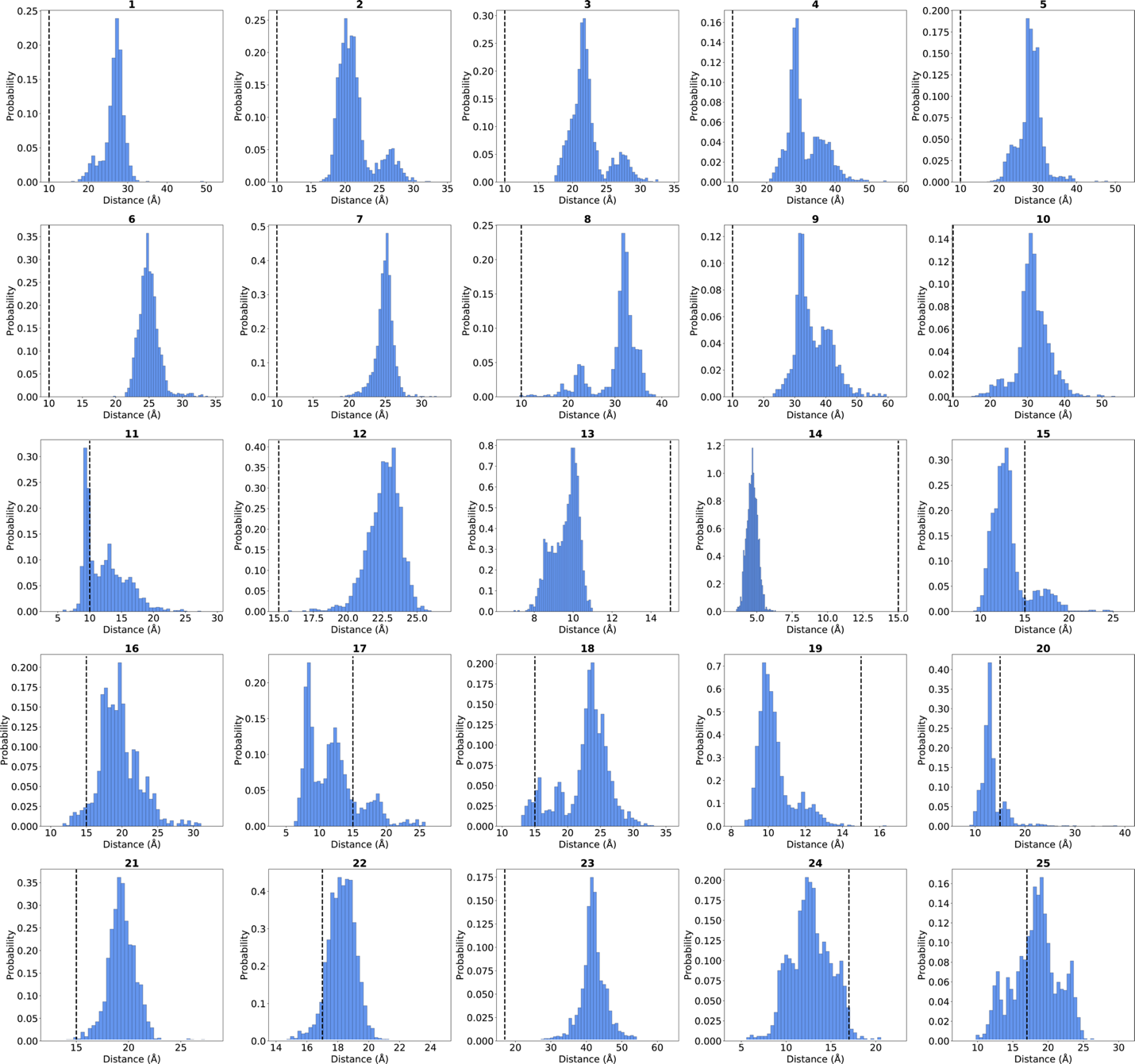

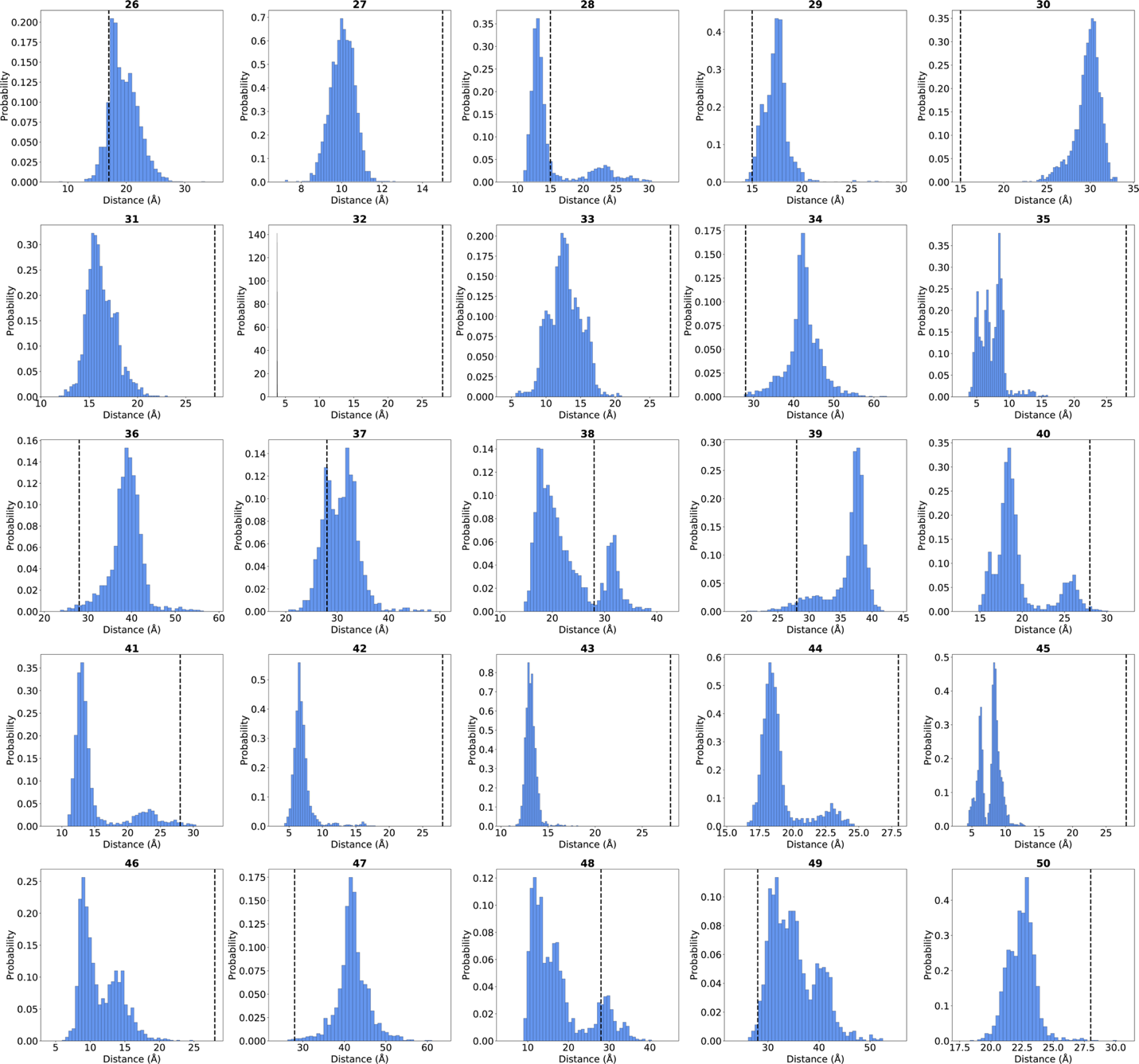

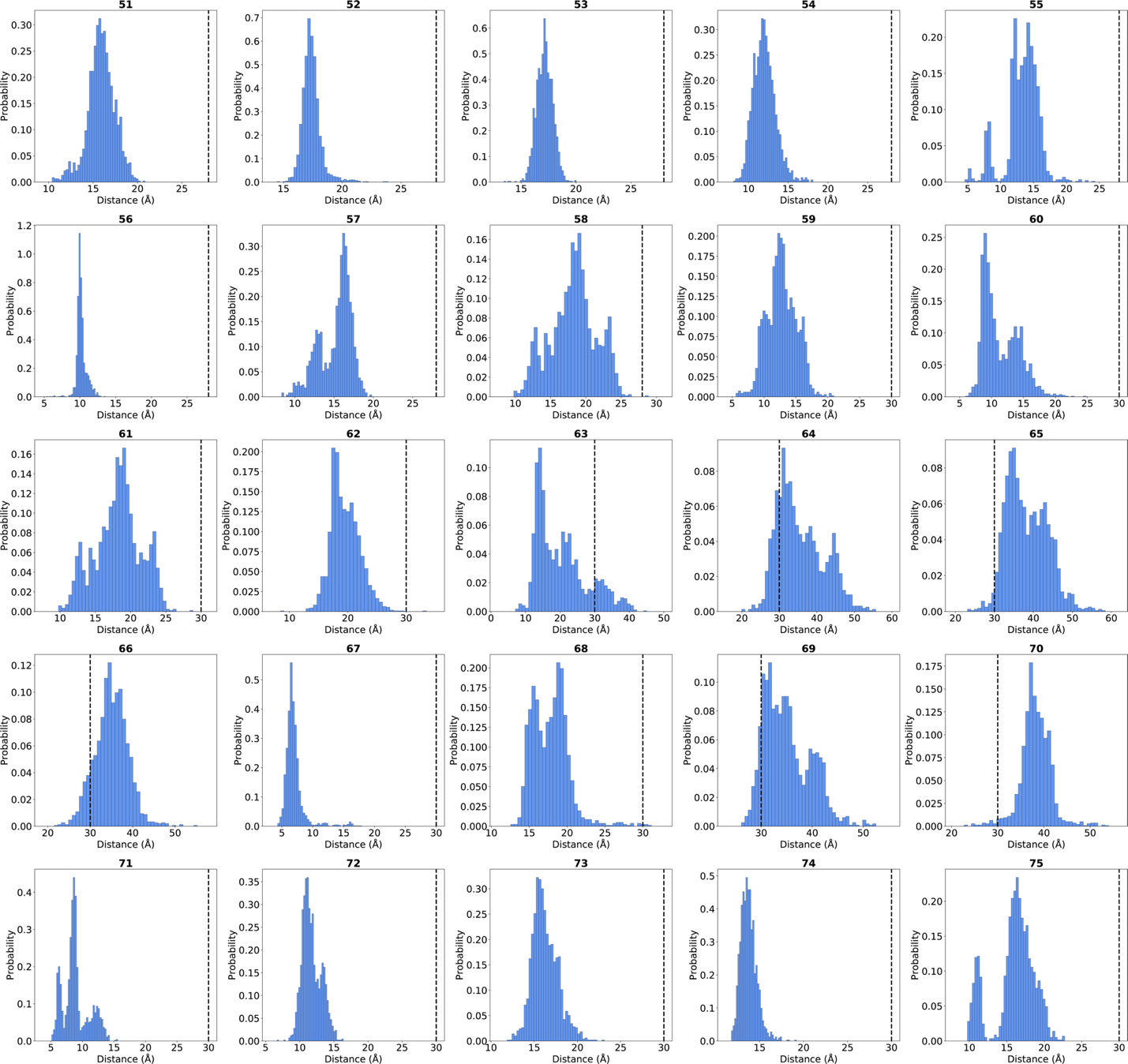

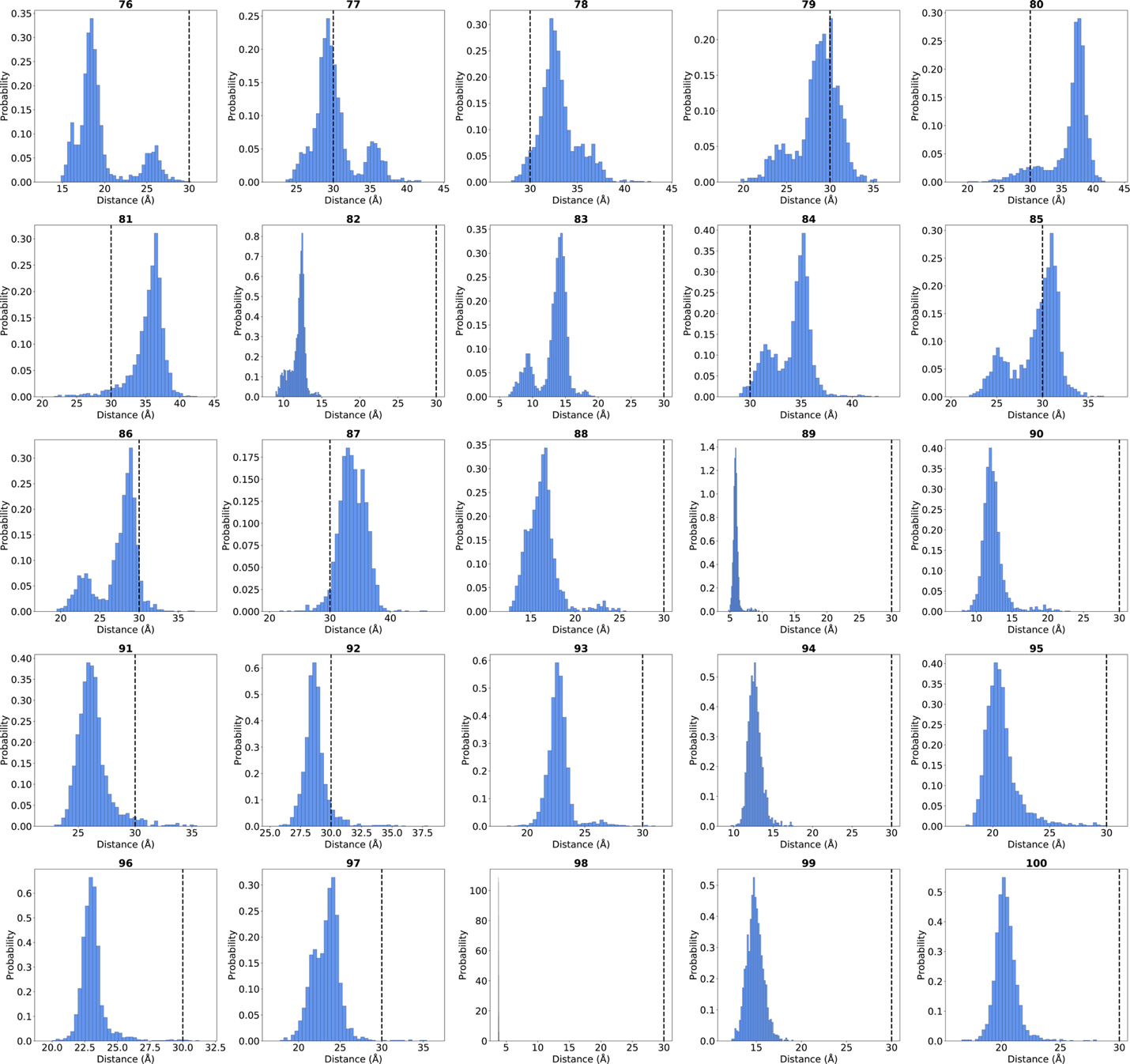

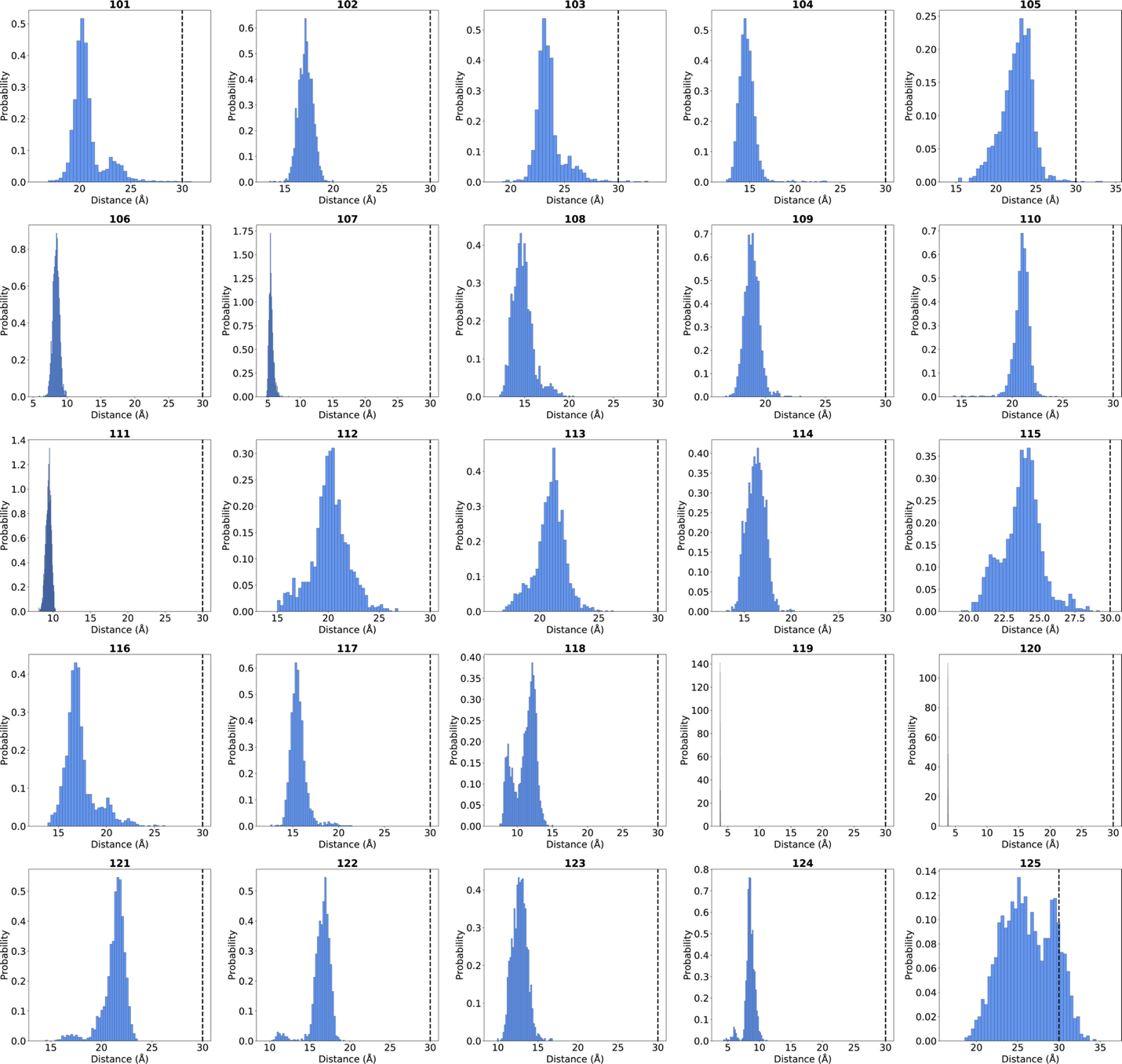

**Fig. S13. Computational distance distributions of the 125 residue pairs in Cluster 7.** The numbers above each subplot are corresponding to the numbers in Table S4 indicating each residue pair. The black dashed line indicates the maximal cross-linking length for each crosslinker: ABAS < 17 Å, EDC < 10 Å, SDA < 15 Å, TATA < 15 Å, CBDPS < 28 Å, and BS3 < 30 Å.

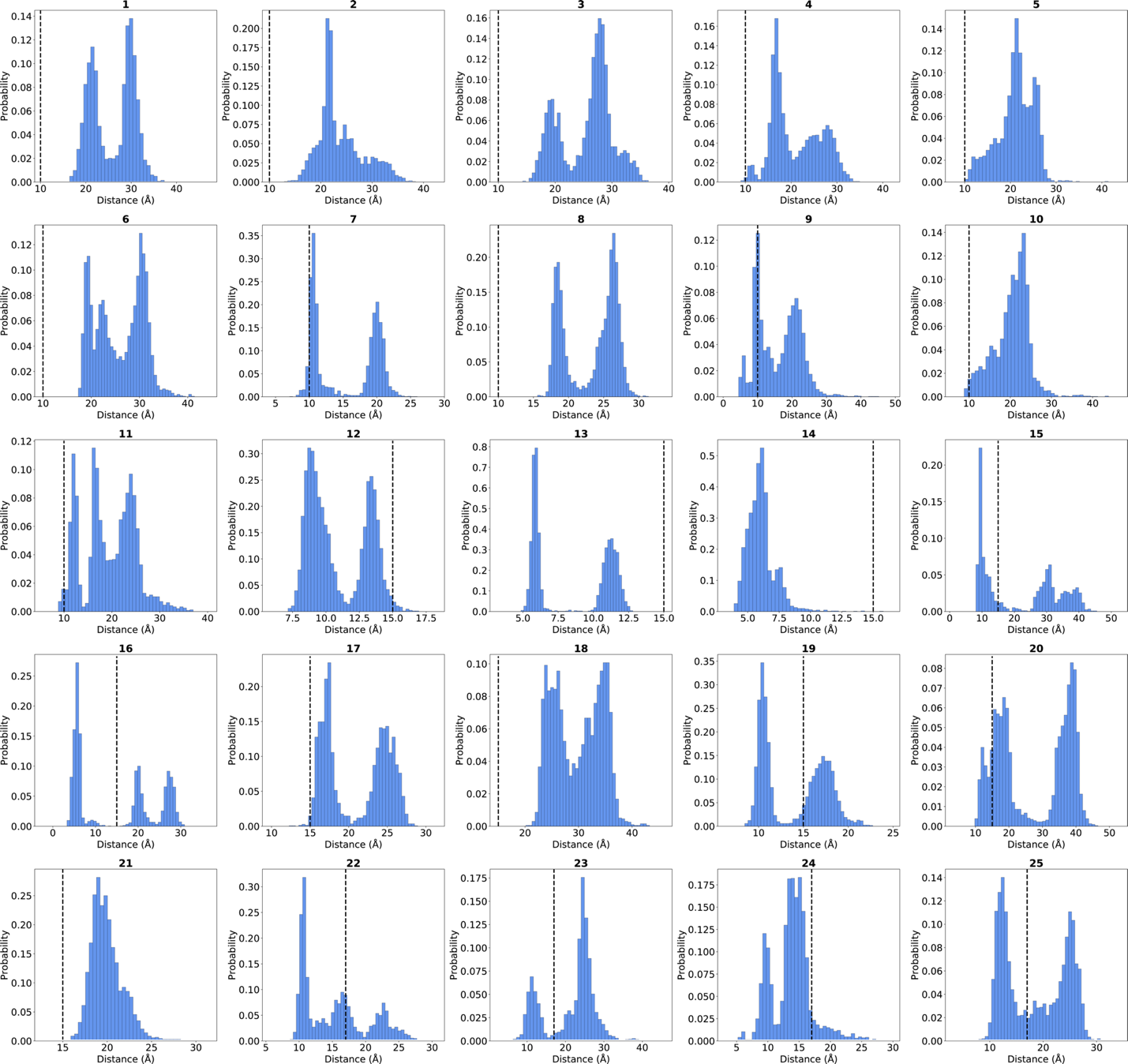

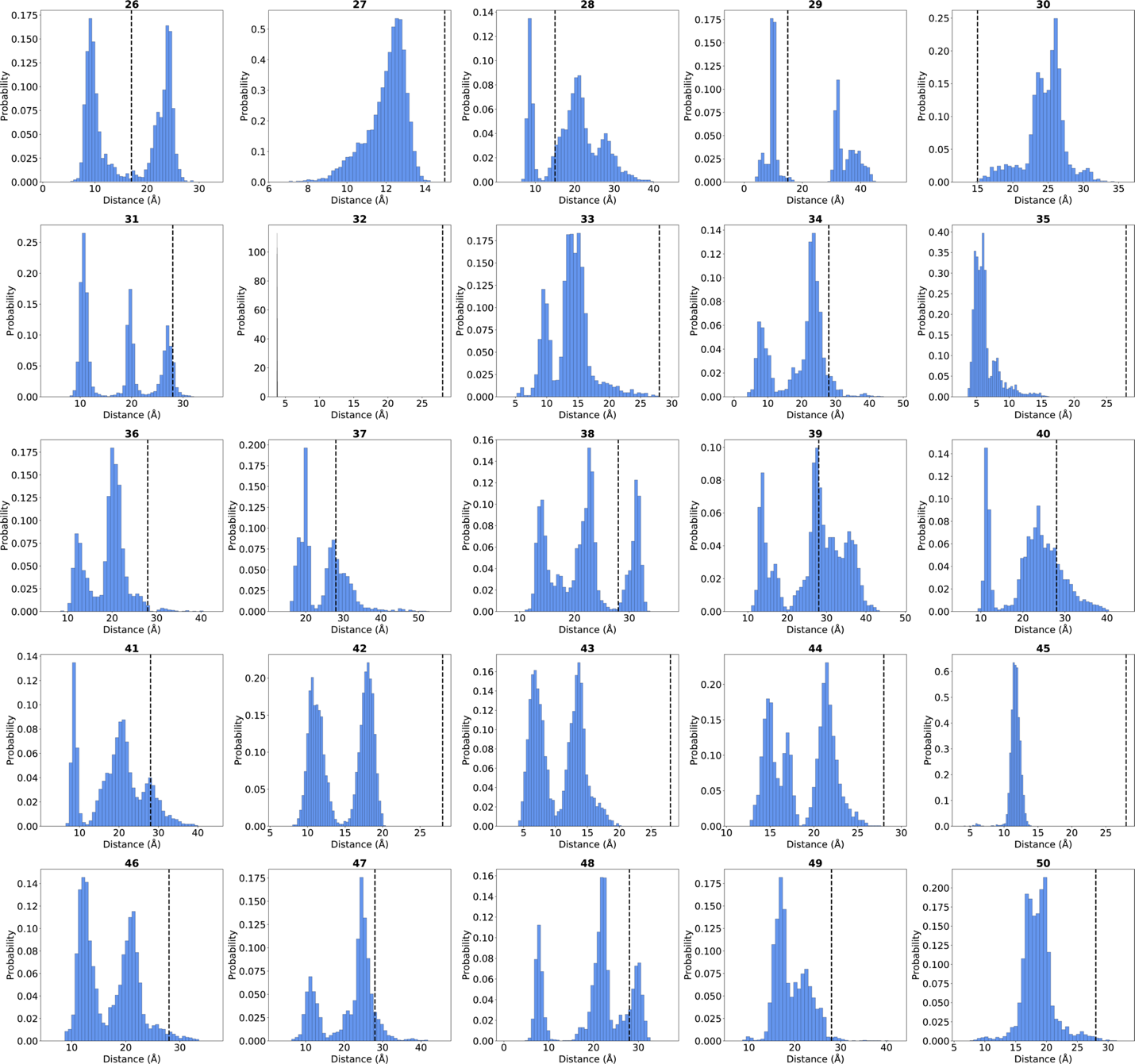

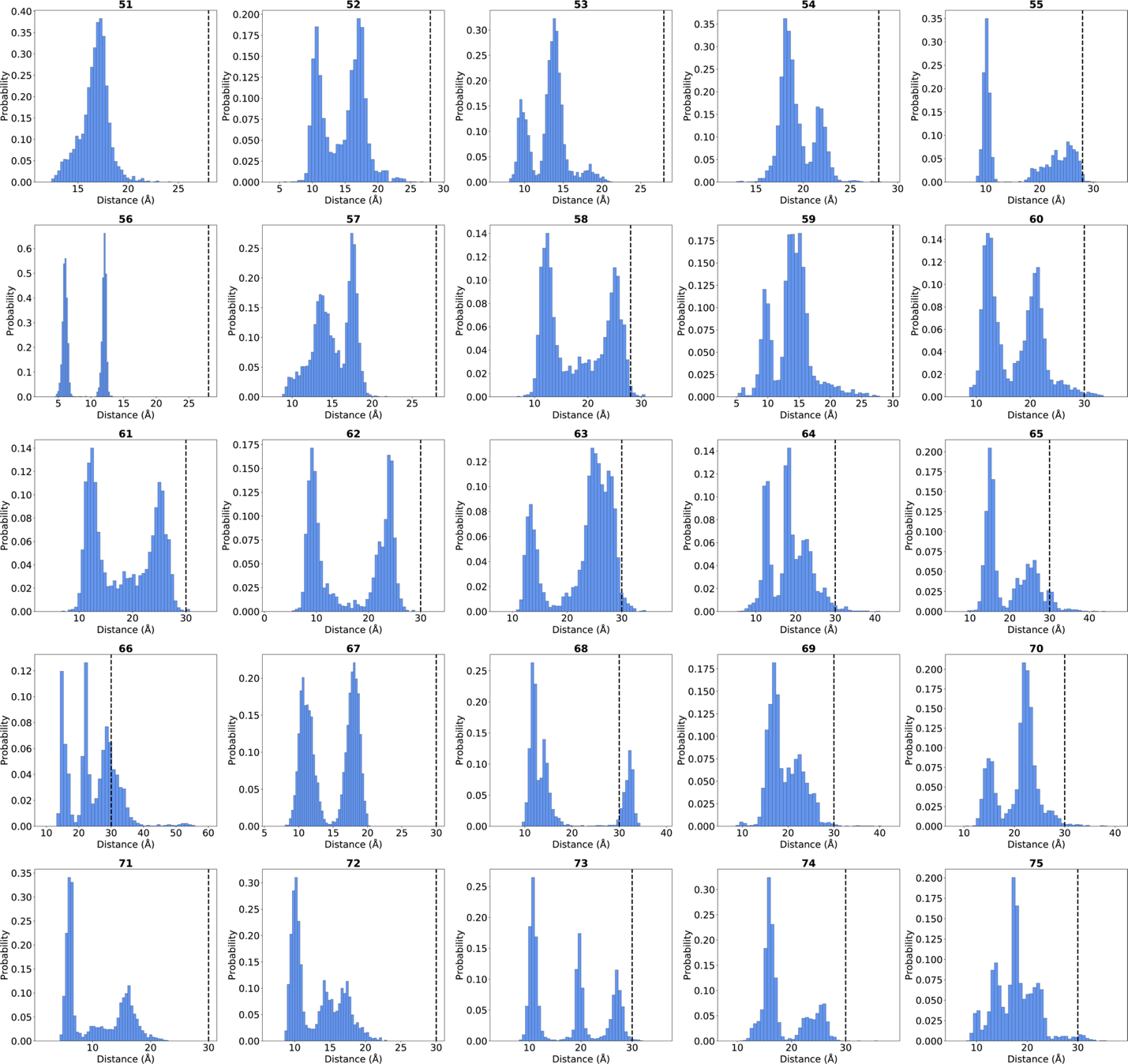

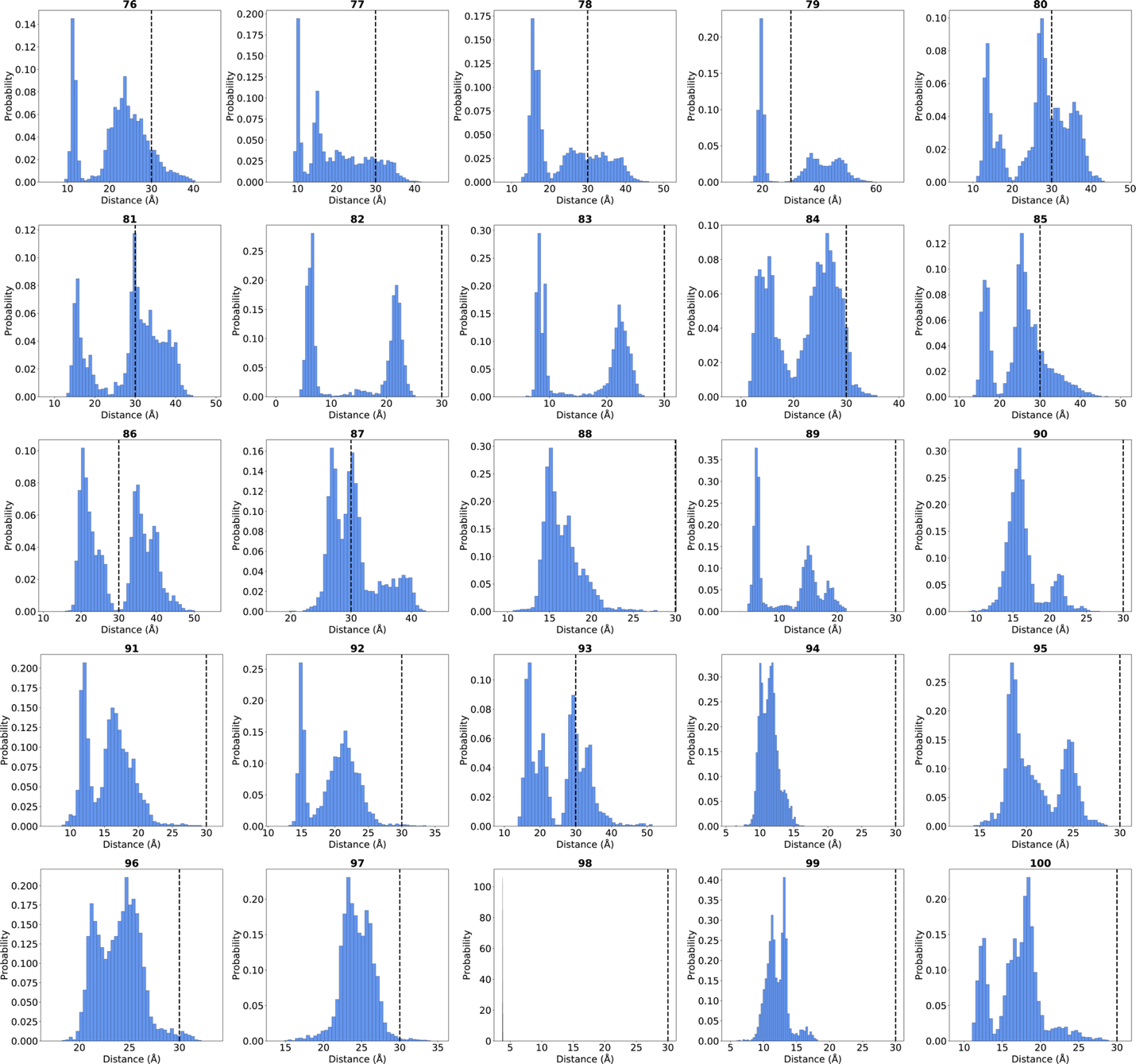

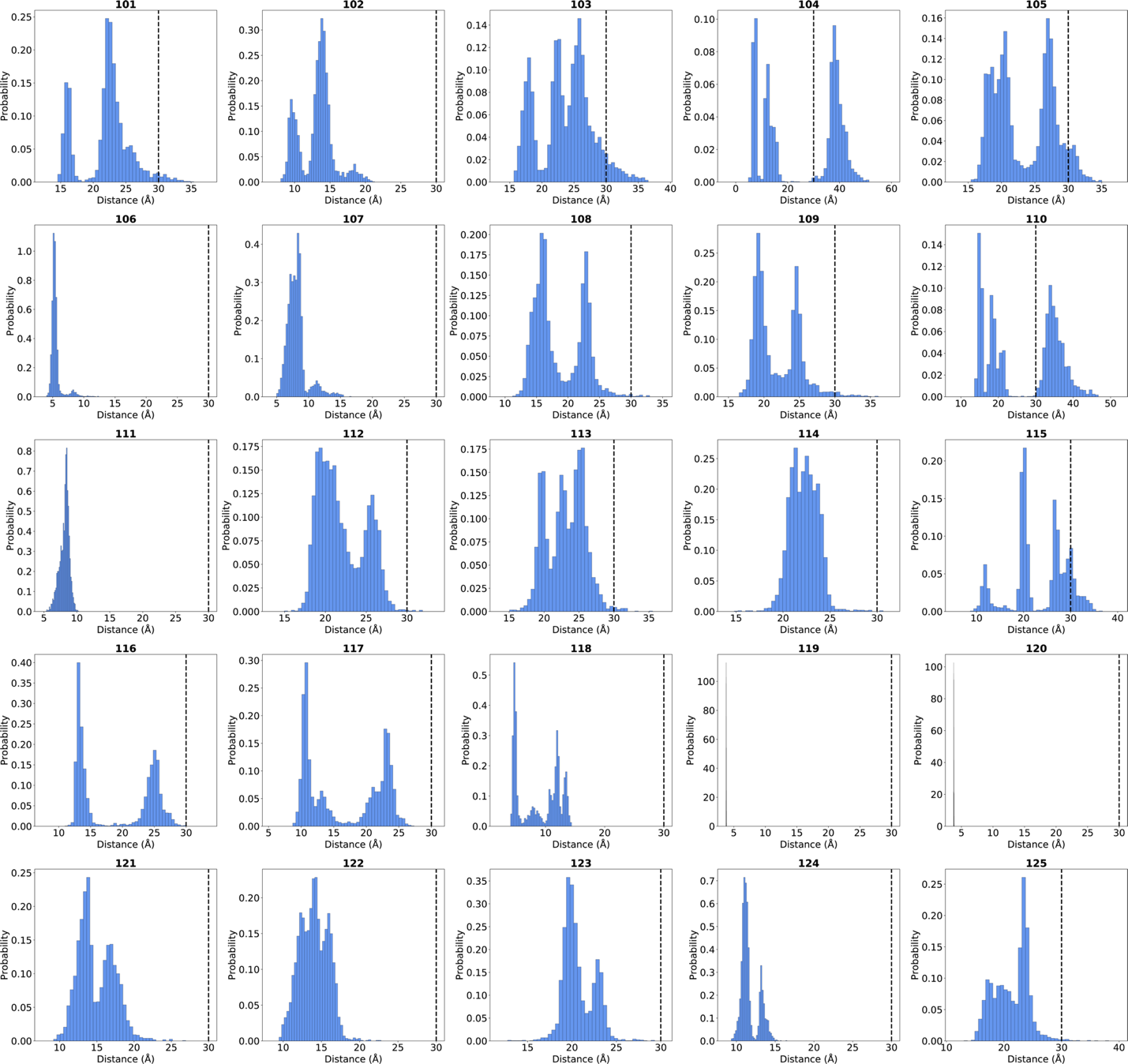

**Fig. S14. Computational distance distributions of the 125 residue pairs in Cluster 8.** The numbers above each subplot are corresponding to the numbers in Table S4 indicating each residue pair. The black dashed line indicates the maximal cross-linking length for each crosslinker: ABAS < 17 Å, EDC < 10 Å, SDA < 15 Å, TATA < 15 Å, CBDPS < 28 Å, and BS3 < 30 Å.

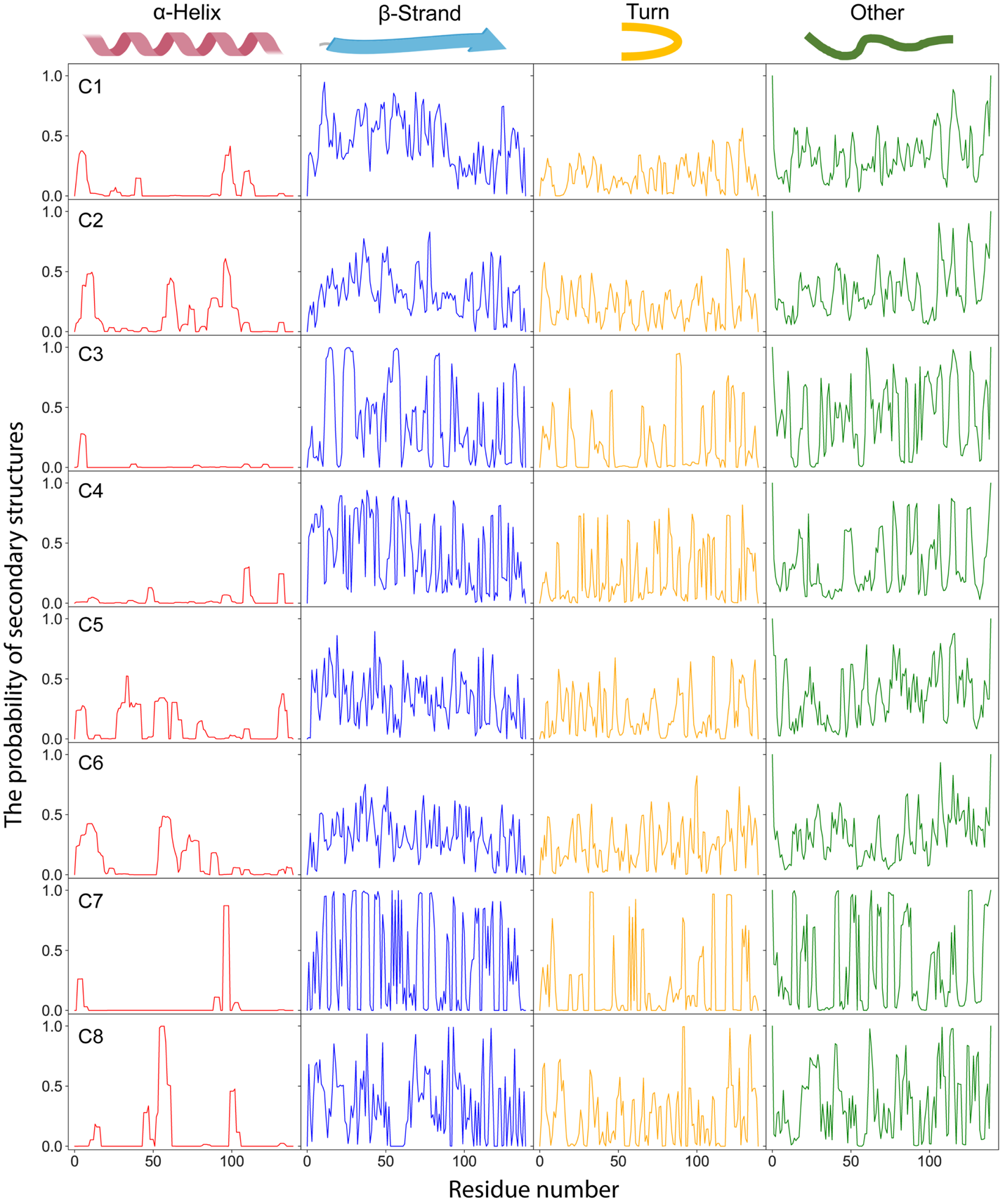

**Fig. S15. The probability of secondary structures for every residue within clusters 1~8 (C1~8).**

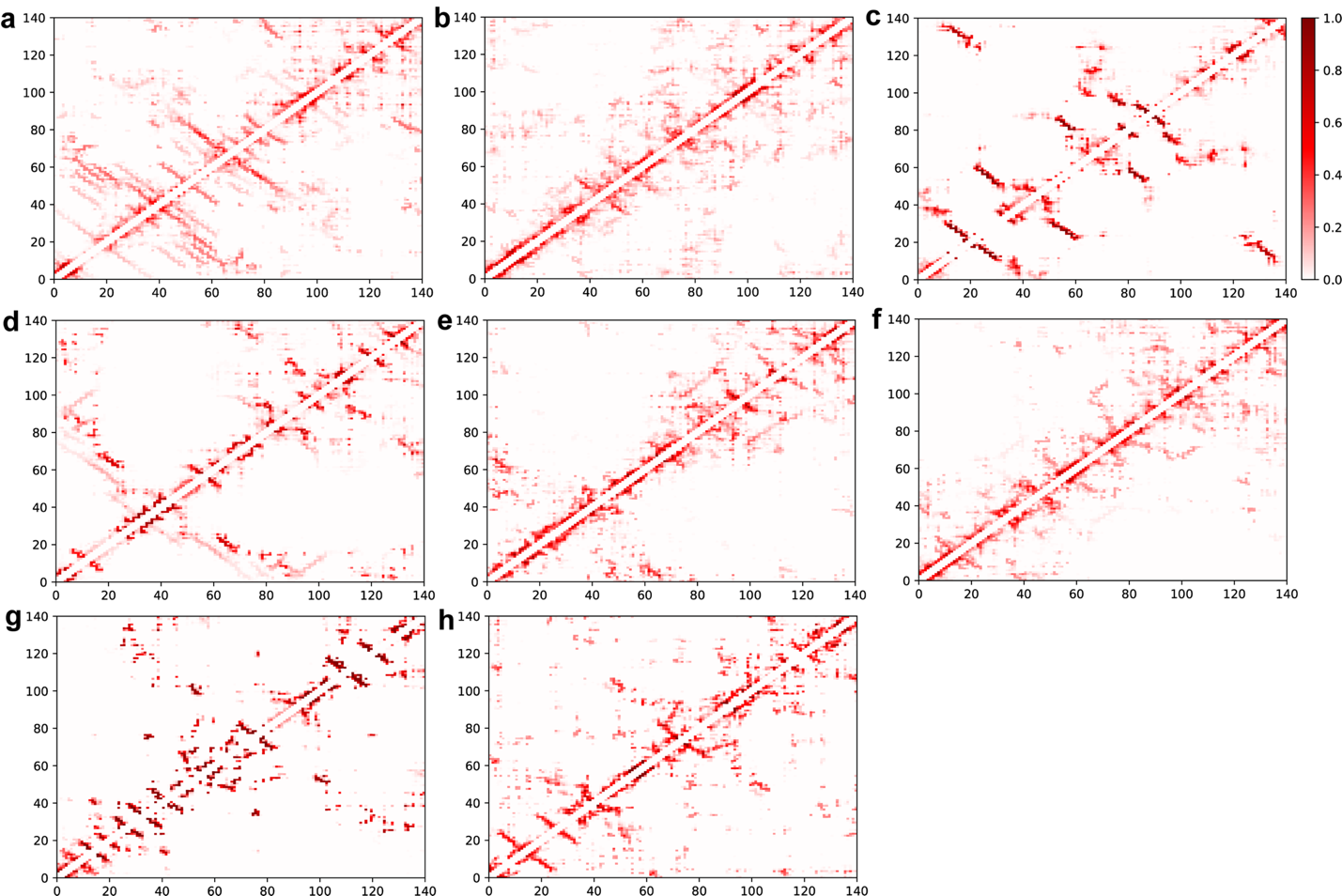

**Fig. S16. The contact frequency maps of clusters 1~8 (panels a~h).** The x- and y-axis indicate residue numbers.

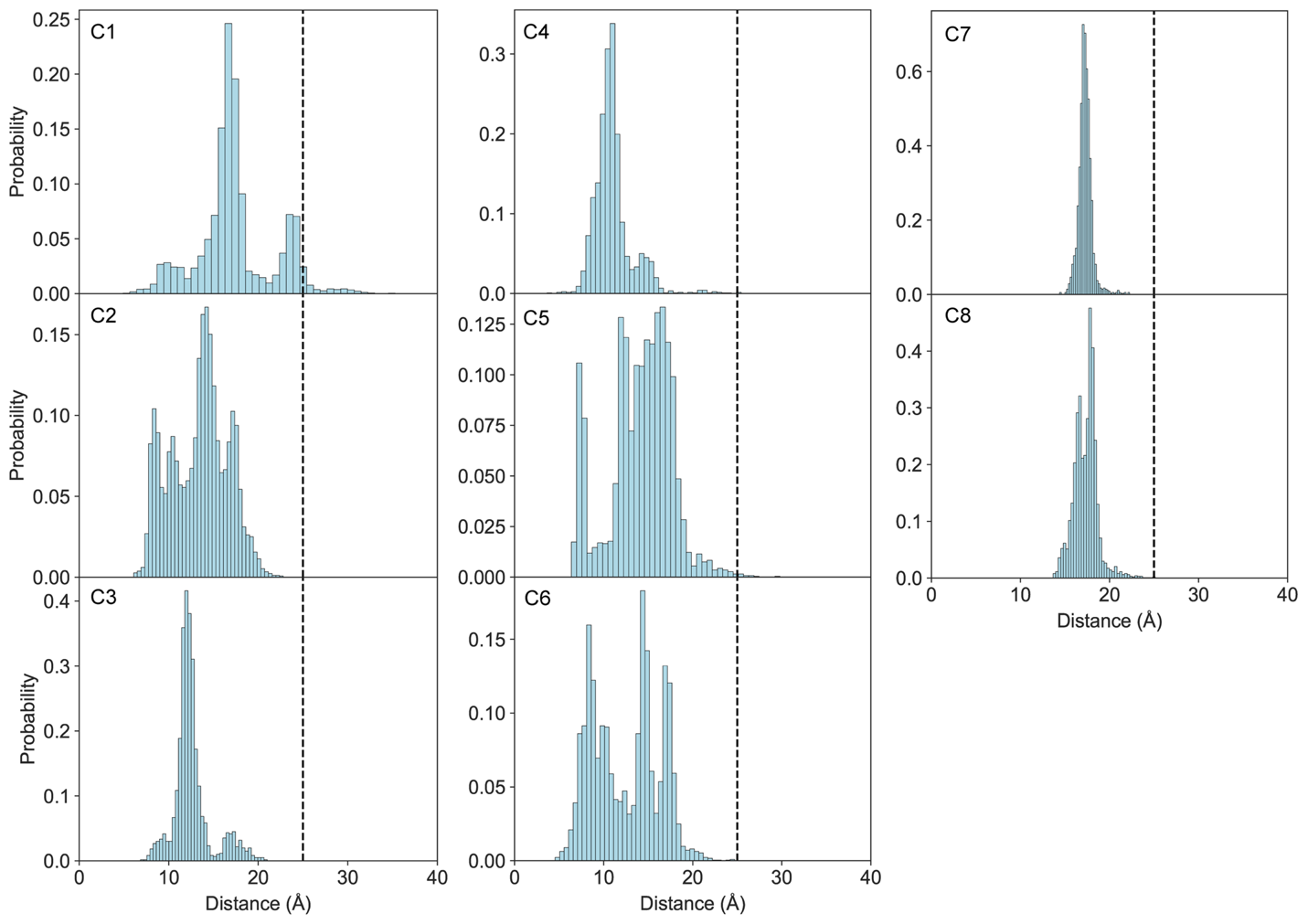

**Fig. S17. The simulated distance distribution between the centers of mass of NAC and C-terminal segments for clusters 1~8 (C1~8).** Structures with distances lower than 25 Å (black dashed vertical lines) are defined to have this long-range interaction.

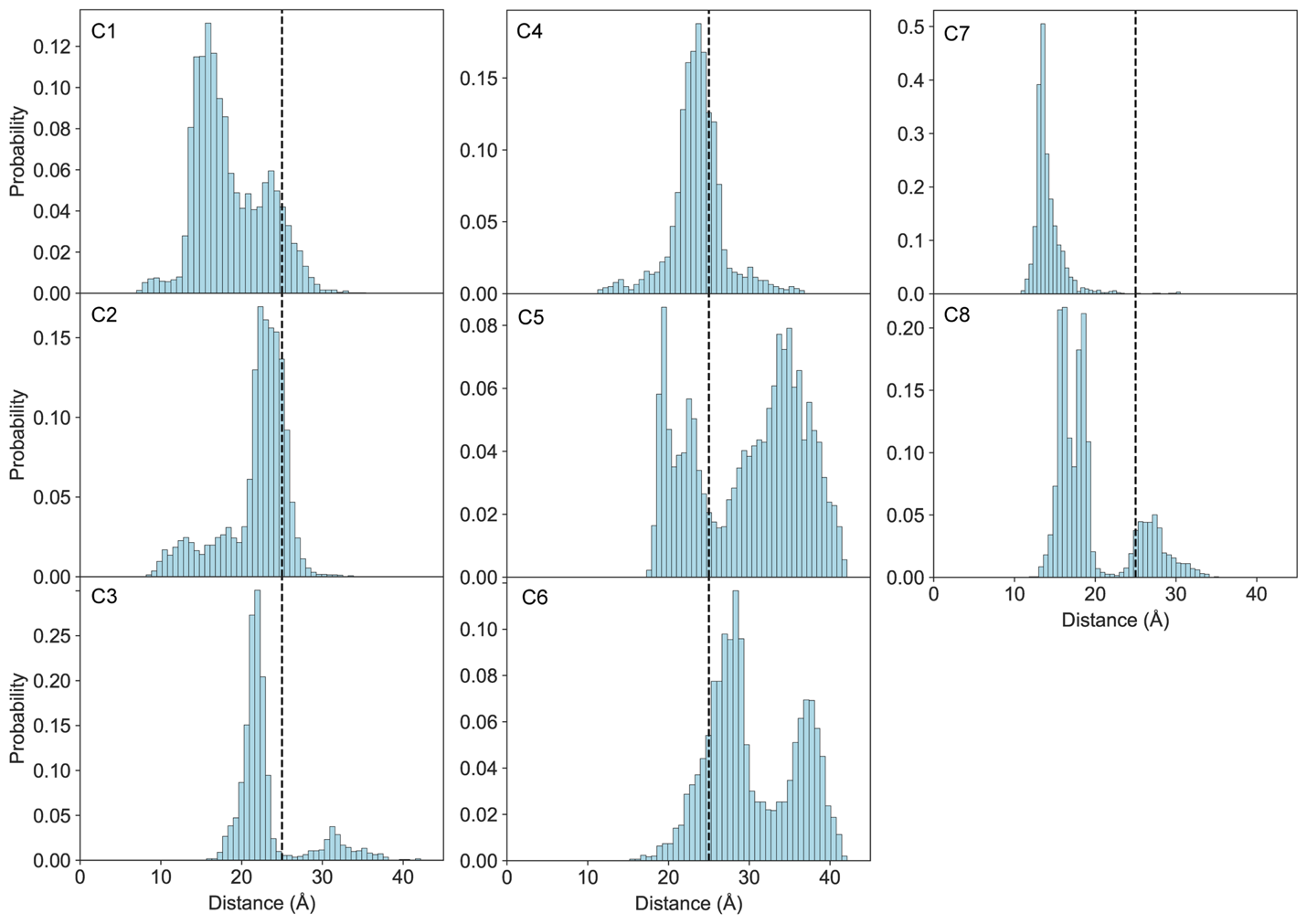

**Fig. S18. The simulated distance distribution between the centers of mass of N- and C- terminal segments for clusters 1~8 (C1~8).** Structures with distances lower than 25 Å (black dashed vertical lines) are defined to have this long-range interaction.

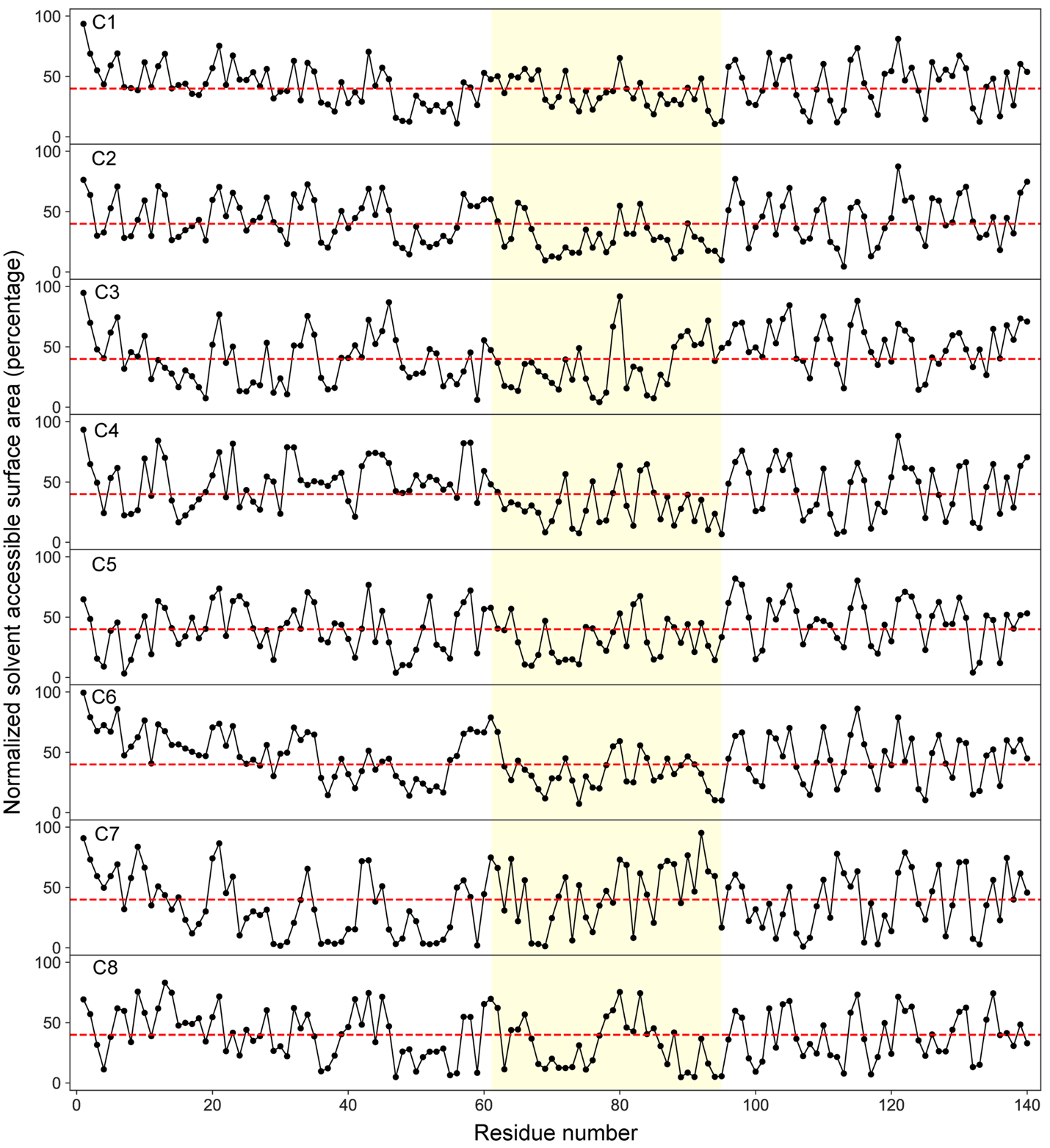

**Fig. S19. The normalized solvent accessible surface area (SASA) of each residue for all the structures that have N- and C- terminal long-range contacts in clusters 1~8 (C1~8).** Residues with SASA < 0.4 are considered buried, while those > 0.4 are considered surface exposed. The NAC segment is highlighted in yellow.

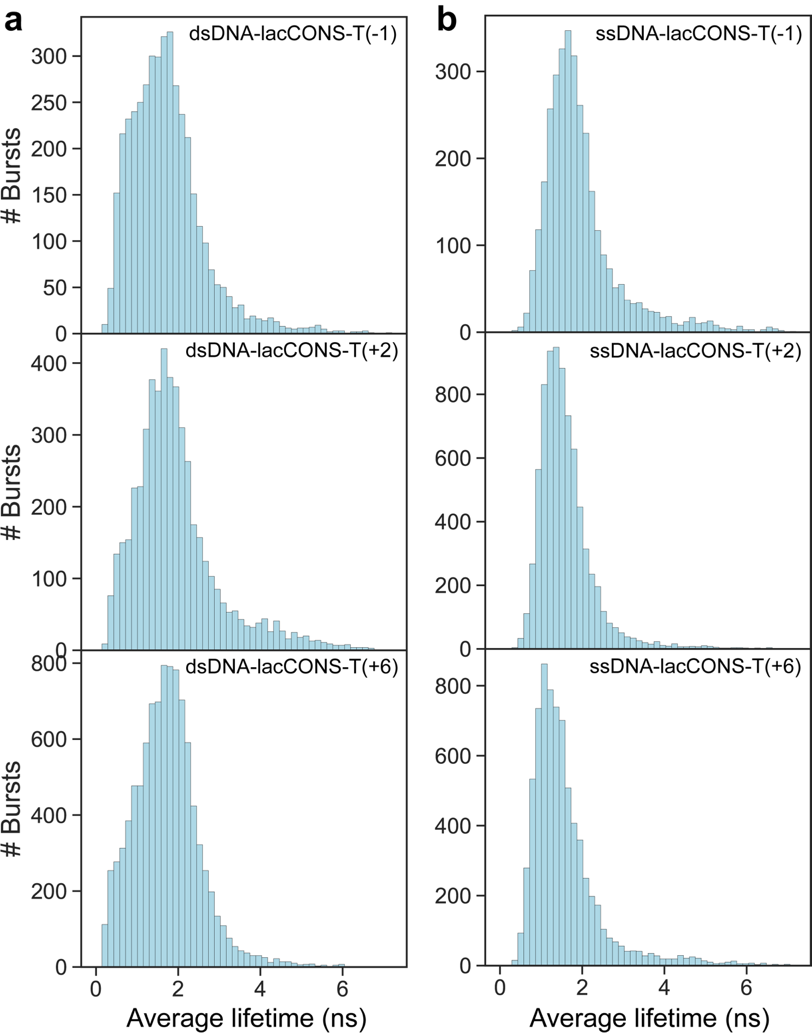

**Fig. S20. The smPIFE experiments of double strand DNA (a) and single strand DNA (b).**

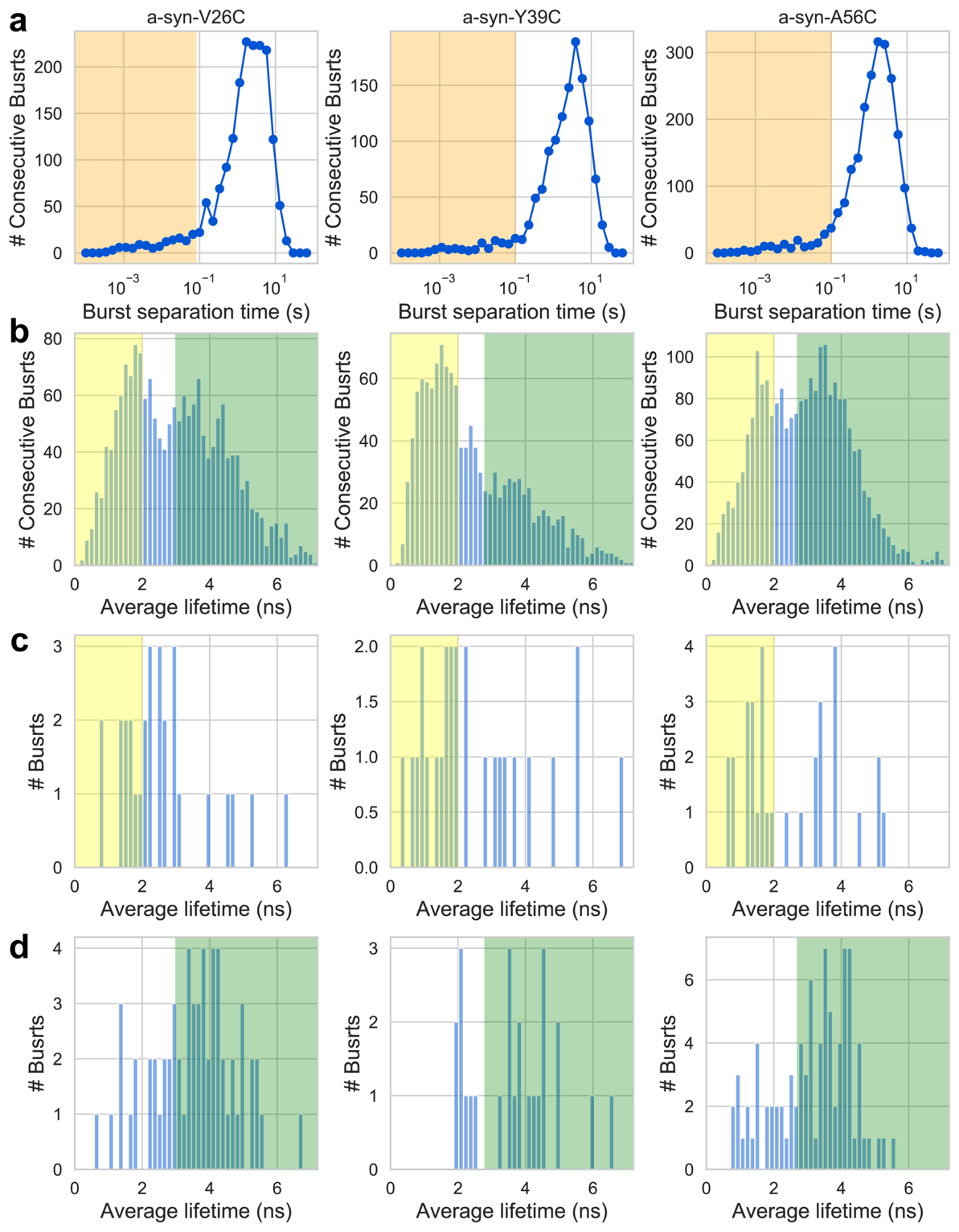

**Fig. S21.** **PIFE burst recurrence analysis shows individual molecules undergo transitions between different average lifetime values within 100 ms.** Each column summarizes the PIFE burst recurrence analysis of Cy3 labeling different residues in α-syn. From top to bottom: a. the histogram of separation times between consecutive single-molecule bursts. The orange shade represents separation times between consecutive bursts of recurring molecules, where the first and second bursts arise from the same molecule; b. the average lifetime histogram of all single-molecule bursts. The yellow and green shades represent the range of average lifetime values chosen to represent values within low and high average lifetime subpopulations, respectively; c or d. the average lifetime histograms of bursts that were separated from a previous burst by a time within the orange-shaded timescale (in panel a), and where the previous burst had an average lifetime within the range represented by the yellow or green shades, respectively.

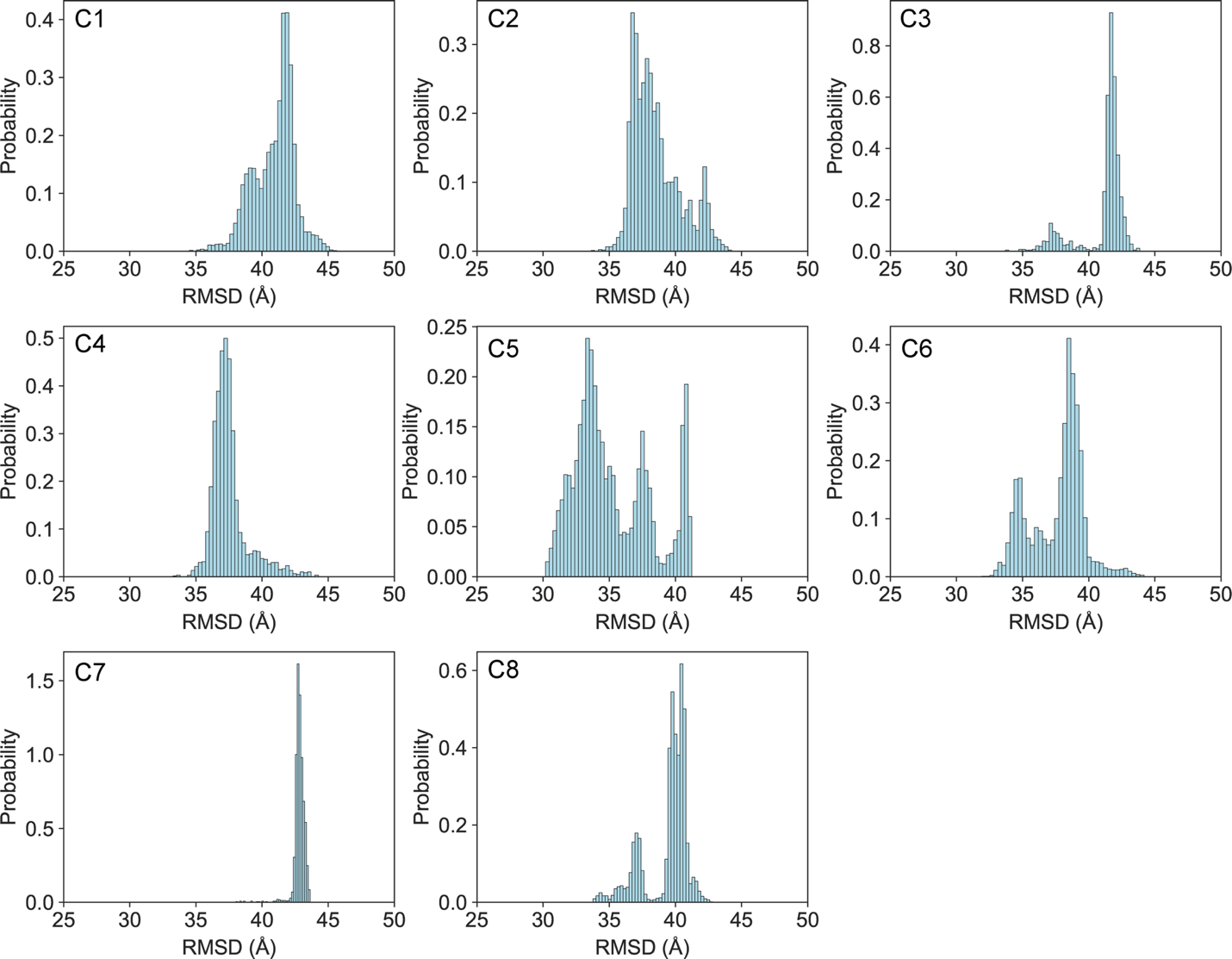

**Fig. S22. The RMSD between the structure of human micelle-bound α-syn (PDB 1XQ8) and all the structures in each cluster.** The RMSD values are calculated by comparing structures of the whole protein.

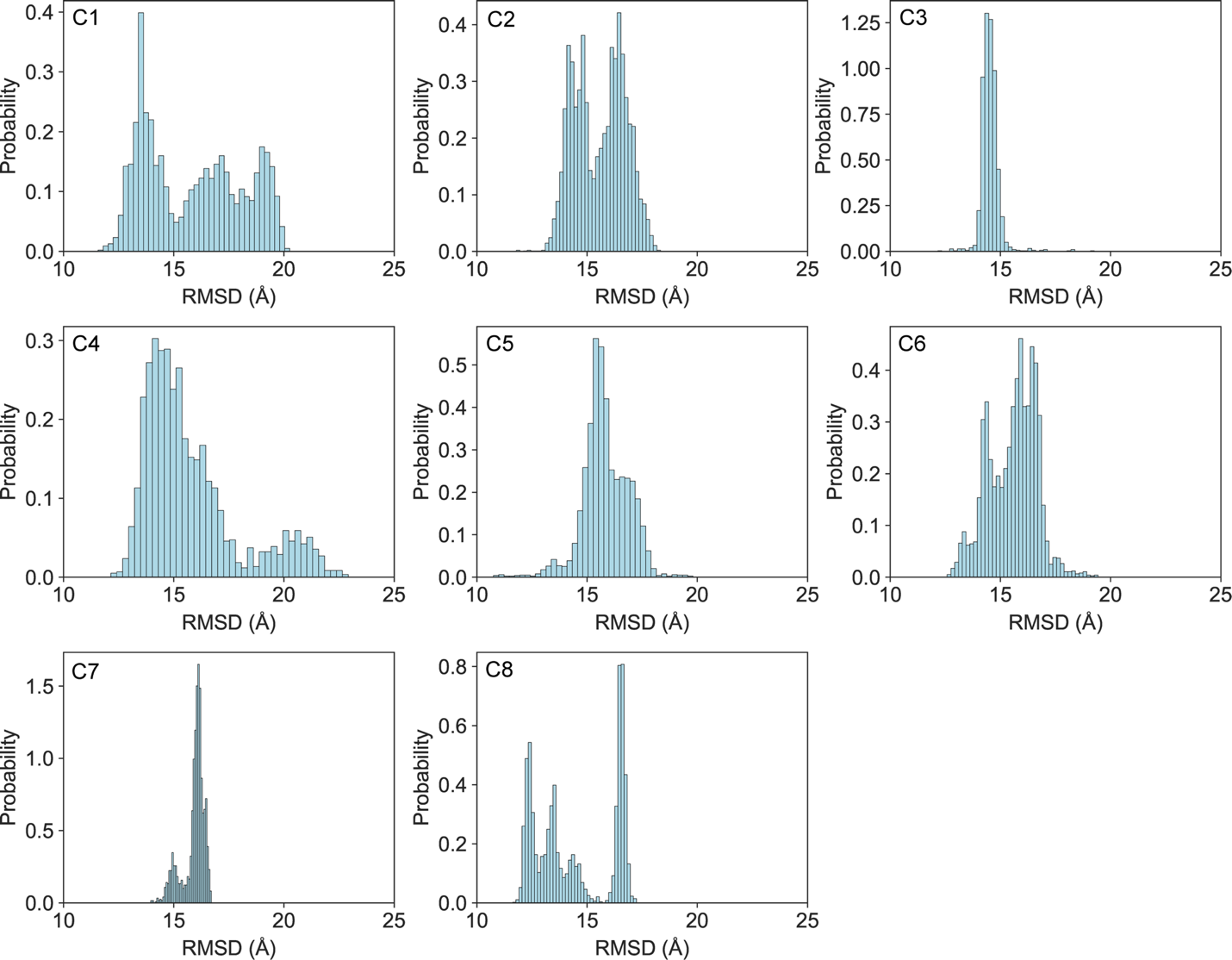

**Fig. S23. The RMSD between the cryo-EM structure of α-syn fiber (PDB 6A6B, residues 37~99) and all the structures in each cluster.** The RMSD values are calculated by comparing structures of the truncated protein from residues 37 to 99.

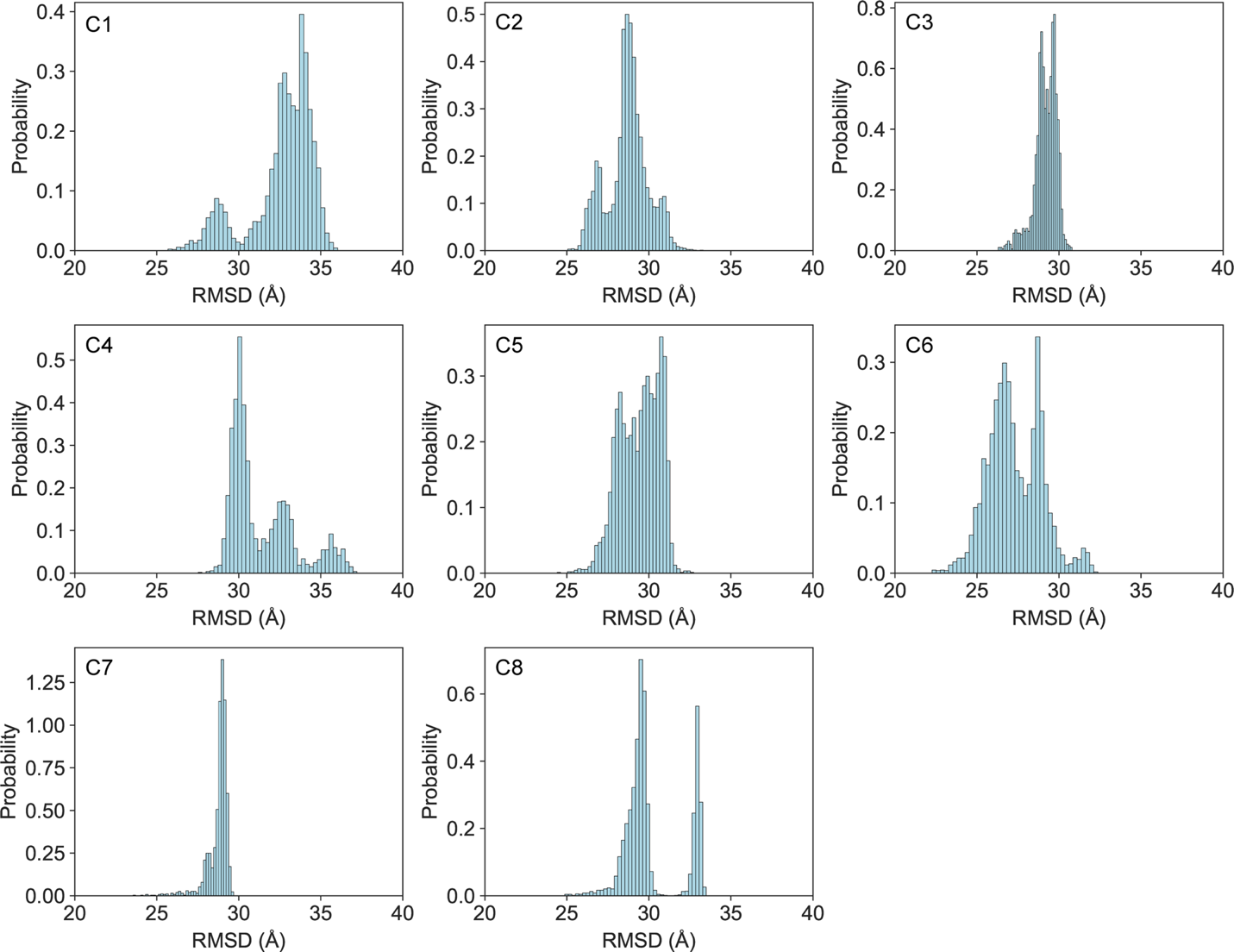

**Fig. S24. The RMSD between the NMR structure of α-syn fiber (PDB 2N0A, 30~98) and all the structures in each cluster.** The RMSD values are calculated by comparing structures of the truncated protein from residues 30 to 98.

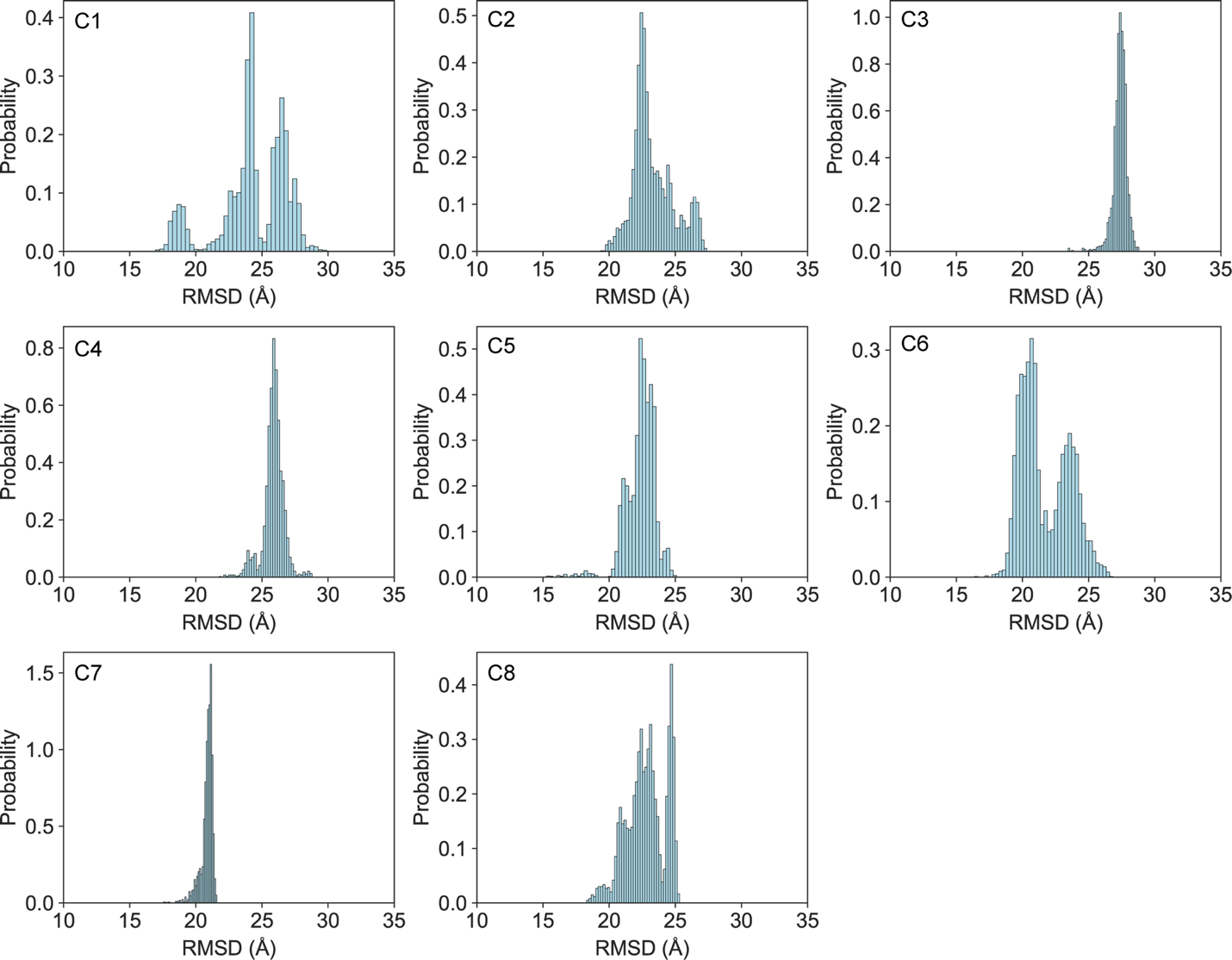

**Fig. S25. The RMSD between the cryo-EM structure of multiple system atrophy Type II-1 α-syn filament (PDB 6XYP, chain A, residues 14~94) and all the structures in each cluster.** The RMSD values are calculated by comparing structures of the truncated protein from residues 14 to 94.

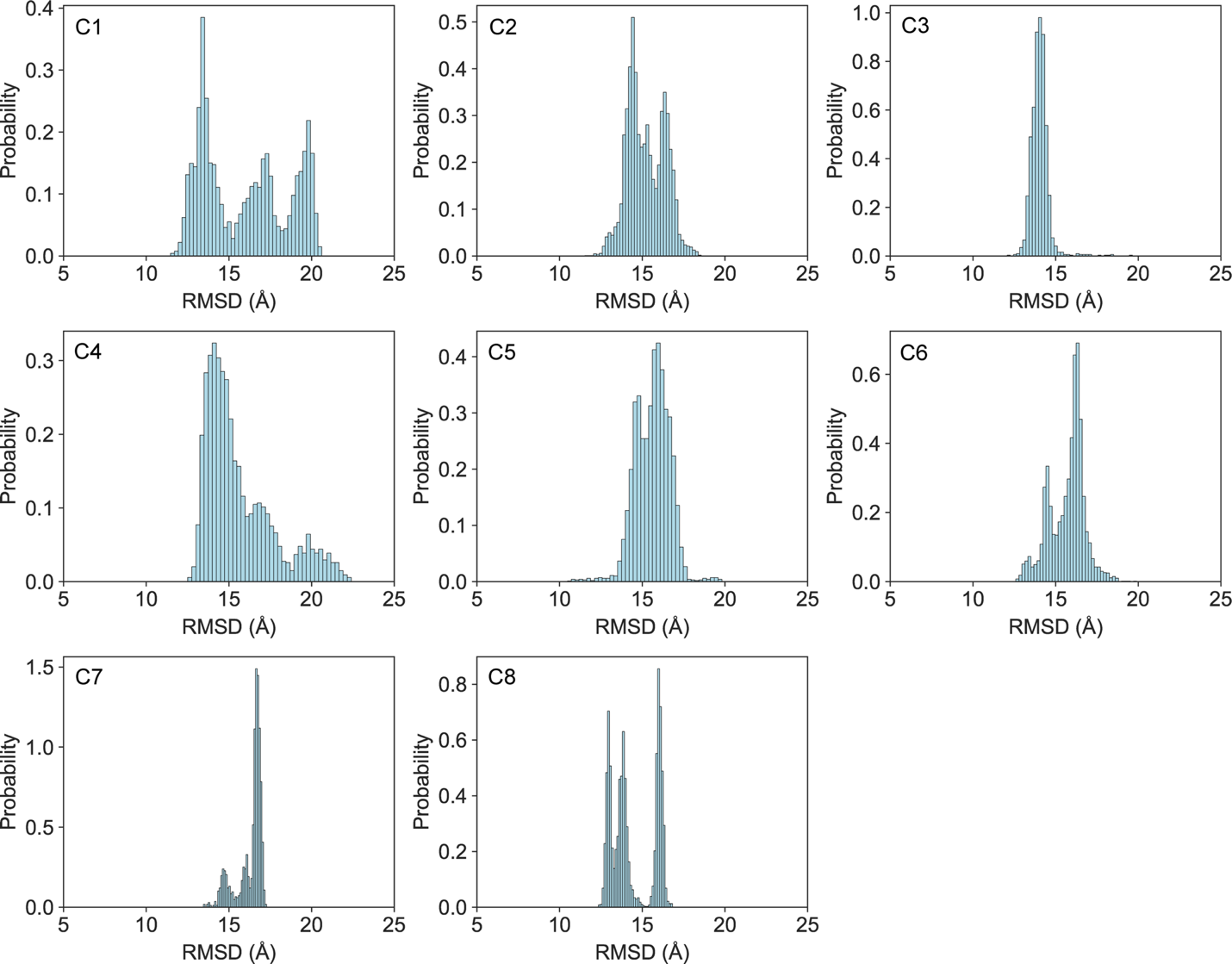

**Fig. S26. The RMSD between the cryo-EM structure of multiple system atrophy Type II-1 α-syn filament (PDB 6XYP, chain B, residues 36~99) and all the structures in each cluster.** The RMSD values are calculated by comparing structures of the truncated protein from residues 36 to 99.

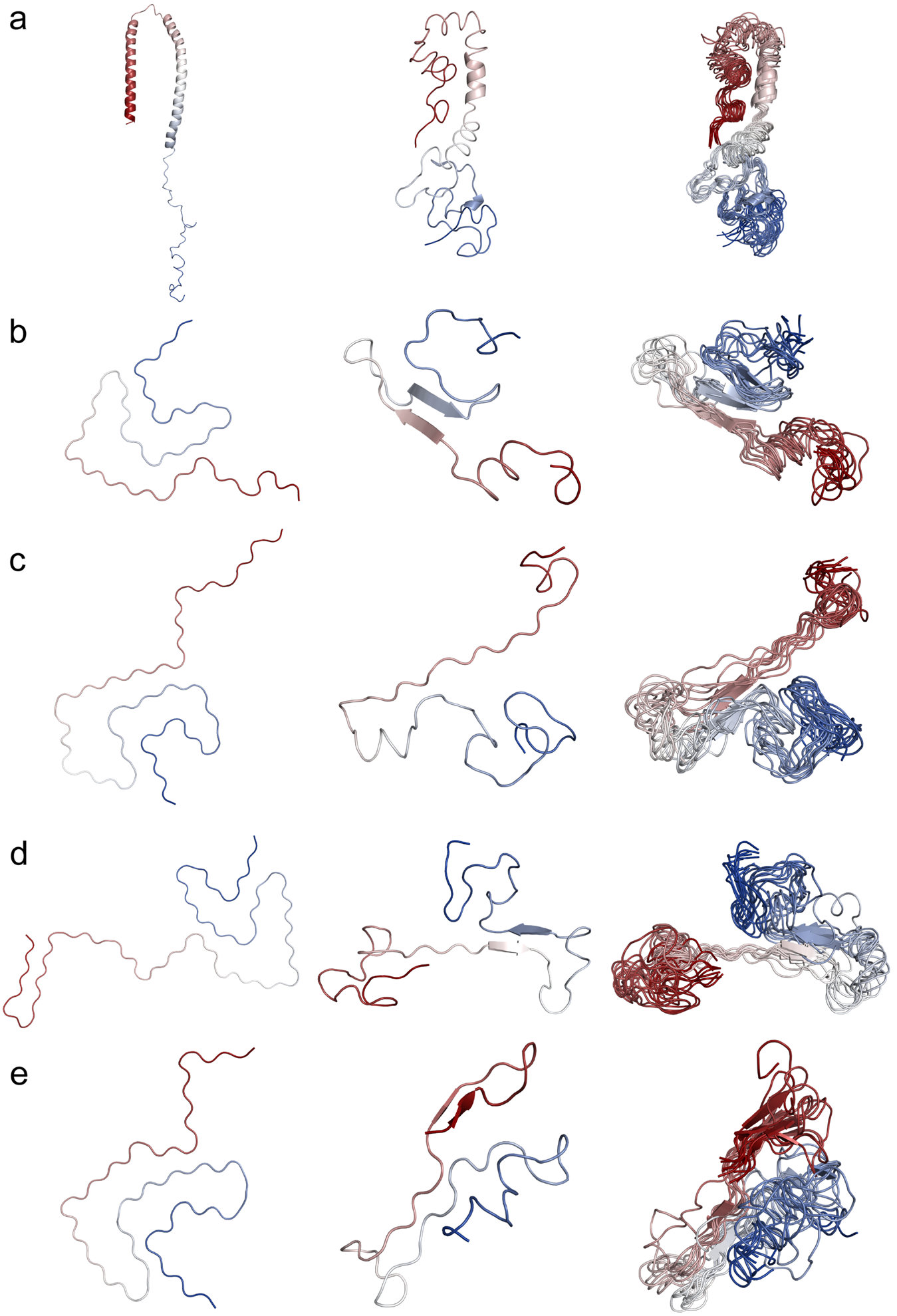

**Fig. S27. Comparison of α-syn structures in PDB with simulated structures.** a, The NMR structure of micelle-bound α-syn (PDB 1XQ8, residues 1~140) on the left panel is compared to the structure in cluster 5 with the lowest RMSD (in the middle). Right panel shows the top 10 structures in cluster 5 with the lowest RMSD. b, The cryo-EM structure of α-syn fiber (PDB 6A6B, residues 37~99) on the left panel is compared to the structure in cluster 1 with the lowest RMSD (in the middle). Right panel shows the top 10 structures in cluster 1 with the lowest RMSD. c, The NMR structure of α-syn fiber (PDB 2N0A, residues 30~98) on the left panel is compared to the structure in cluster 1 with the lowest RMSD (in the middle). Right panel shows the top 10 structures in cluster 1 with the lowest RMSD. d, The cryo-EM structure of multiple system atrophy Type II-1 α-syn filament (PDB 6XYP, chain A, residues 14~94) is compared to the structure in cluster 1 with the lowest RMSD (in the middle). Right panel shows the top 10 structures in cluster 1 with the lowest RMSD. e, The cryo-EM structure of multiple system atrophy Type II-1 α-syn filament (PDB 6XYP, chain B, residues 36~99) is compared to the structure in cluster 1 with the lowest RMSD (in the middle). Right panel shows the top 10 structures in cluster 1 with the lowest RMSD.

**Table S1. The replica exchange rate of DMD simulations.**

| Temperature 1  (kcal/mol•k_B_) | Temperature 2  (kcal/mol•k_B_) | Exchange rate |
| --- | --- | --- |
| 0.35 | 0.36 | 0.551 |
| 0.36 | 0.37 | 0.549 |
| 0.37 | 0.38 | 0.577 |
| 0.38 | 0.39 | 0.577 |
| 0.39 | 0.40 | 0.555 |
| 0.40 | 0.41 | 0.591 |
| 0.41 | 0.42 | 0.605 |
| 0.42 | 0.43 | 0.591 |
| 0.43 | 0.44 | 0.589 |
| 0.44 | 0.45 | 0.599 |
| 0.45 | 0.46 | 0.636 |
| 0.46 | 0.47 | 0.605 |
| 0.47 | 0.48 | 0.605 |
| 0.48 | 0.49 | 0.598 |
| 0.49 | 0.50 | 0.594 |
| 0.50 | 0.51 | 0.594 |
| 0.51 | 0.52 | 0.577 |
| 0.52 | 0.53 | 0.535 |
| 0.53 | 0.54 | 0.540 |
| 0.54 | 0.55 | 0.554 |
| 0.55 | 0.56 | 0.494 |
| 0.56 | 0.57 | 0.469 |
| 0.57 | 0.58 | 0.493 |
| 0.58 | 0.59 | 0.449 |
| 0.59 | 0.60 | 0.550 |

**Table S2. Pairwise RMSD (Å) between the eight clusters.**

| Clusters | C1 | C2 | C3 | C4 | C5 | C6 | C7 | C8 |
| --- | --- | --- | --- | --- | --- | --- | --- | --- |
| C1 | 0.0 | 16.3 | 20.9 | 19.1 | 19.7 | 21.7 | 17.3 | 15.6 |
| C2 | 16.3 | 0.0 | 15.7 | 13.0 | 14.7 | 16.6 | 15.2 | 13.8 |
| C3 | 20.9 | 15.7 | 0.0 | 16.7 | 18.3 | 17.4 | 15.7 | 16.1 |
| C4 | 19.1 | 13.0 | 16.7 | 0.0 | 13.6 | 18.3 | 18.1 | 15.9 |
| C5 | 19.7 | 14.7 | 18.3 | 13.6 | 0.0 | 17.0 | 18.9 | 15.3 |
| C6 | 21.7 | 16.6 | 17.4 | 18.3 | 17.0 | 0.0 | 15.1 | 17.5 |
| C7 | 17.3 | 15.2 | 15.7 | 18.1 | 18.9 | 15.1 | 0.0 | 14.2 |
| C8 | 15.6 | 13.8 | 16.1 | 15.9 | 15.3 | 17.5 | 14.2 | 0.0 |

**Table S3. The average pairwise RMSD within each cluster**

| Clusters | RMSD (Å) |
| --- | --- |
| C1 | 13.5 |
| C2 | 12.2 |
| C3 | 8.0 |
| C4 | 11.2 |
| C5 | 11.7 |
| C6 | 13.6 |
| C7 | 5.3 |
| C8 | 10.0 |

**Table S4. Cross-linking data used to verify predicted conformational ensemble of α-syn**

| No. | Crosslinker | Sequence 1 | Sequence 2 | Crosslink |
| --- | --- | --- | --- | --- |
| 1 | EDC | G_7_LSK_10_ | E_126_MPSEE_131_ | K10-E126 |
| 2 | EDC | E_110_GIL_113_ | K_43_TKEGVVH_50_ | E110-K43 |
| 3 | EDC | S_42_KTK_45_ | E_110_GILE_114_ | K43-E114 |
| 4 | EDC | M_5_K_6_ | E_110_GILE_114_ | K6-E110 |
| 5 | EDC | M_5_K_6_ | E_126_MPSEE_131_ | K6-E126 |
| 6 | EDC | K_80_TVE_83_ | E_126_MPSEE_131_ | K80-E126 |
| 7 | EDC | E_110_GIL_113_ | T_92_GFVK_96_ | E110-K96 |
| 8 | EDC | V_95_KKD_98_ | Y_136_EPEA_140_-Cterm | K96-E137 |
| 9 | EDC | Nterm-M_1_DVF_4_ | E_114_DMPVDPD_121_ | N-term-E114 |
| 10 | EDC | Nterm-M_1_DVF_4_ | E_126_MPSEE_131_ | N-term-E126 |
| 11 | EDC | Nterm-M_1_DVF_4_ | S_9_KAKEG_14_ | N-term-E13 |
| 12 | TATA | D_119_PDNEAY_125_ | E_130_EGYQ_134_ | D121-E130 |
| 13 | TATA | E_114_DMPVDPD_121_ | Y_125_EMP_128_ | D121-Y125 |
| 14 | TATA | G_31_KTKEG_36_ | V_37_LYVG_41_ | G31-V37 |
| 15 | TATA | K_45_EGVVH_50_ | A_107_PQ_109_ | K45-A107 |
| 16 | TATA | M_5_KGLSKA_11_ | Q_24_GVA_27_ | M5-G25 |
| 17 | TATA | M_5_KGL_8_ | V_16_A_17_ | M5-V16 |
| 18 | TATA | N_65_VGGAVVT_72_ | P_138_E_139_ | N65-P138 |
| 19 | TATA | P_117_VDPDNEAY_125_ | P_128_SE_130_ | P117-P128 |
| 20 | TATA | V_26_AE_28_ | A_107_PQ_109_ | V26-A107 |
| 21 | TATA | V_82_EGA_85_ | P_117_V_118_ | V82-P117 |
| 22 | ABAS | G_47_VVH_50_ | V_95_KKD_98_ | H50-K96 |
| 23 | ABAS | V_95_KK_97_ | Nterm-M_1_DVF_4_ | K97-N-term |
| 24 | ABAS | S_9_K_10_ | Nterm-M_1_DVF_4_ | K10-N-term |
| 25 | ABAS | A_19_EKT_22_ | Nterm-M_1_DVF_4_ | K21-N-term |
| 26 | ABAS | Nterm-M_1_DVF_4_ | K_23_QGV_26_ | N-term-K23 |
| 27 | SDA | G_25_VAE_28_ | G_31_KTKEGVLY_39_ | E28-K32 |
| 28 | SDA | E_35_GVLYVGSK_43_ | M_5_KGLSKAKEGVVAAAE_20_ | K43-K12 |
| 29 | SDA | E_105_GAPQ_109_ | V_40_GSKTKEGVVH_50_ | E105-K45 |
| 30 | SDA | A_85_GSIAAATGFVK_96_ | E_35_GVLYVGSKTKEGV_48_ | K96-S42 |
| 31 | CBDPS | S_9_K_10_ | G_31_KTK_34_ | K10-K32 |
| 32 | CBDPS | G_93_FVK_96_ | K_97_D_98_ | K96-K97 |
| 33 | CBDPS | K_10_A_11_ | Nterm-M_1_DVF_4_ | K10-N |
| 34 | CBDPS | V_95_K_96_ | Nterm-M_1_DVF_4_ | K96­-N |
| 35 | CBDPS | M_5_K_6_ | Nterm-M_1_DVF_4_ | K6-N |
| 36 | CBDPS | M_5_K_6_ | T_92_GFVK_96_ | K6-K96 |
| 37 | CBDPS | M_5_K_6_ | K_80_TVEGA_85_ | K6-K80 |
| 38 | CBDPS | M_5_K_6_ | K_45_EGVVH_50_ | K6-K45 |
| 39 | CBDPS | V_95_KK_97_ | G_7_LSKAK_12_ | K96-K12 |
| 40 | CBDPS | A_11_K_12_ | T_44_KEGVVH_50_ | K12-K45 |
| 41 | CBDPS | S_42_KT_44_ | K_45_EGVVH_50_ | K43-K45 |
| 42 | CBDPS | M_5_KG_7_ | A_11_KEGVVA_17_ | K6-K12 |
| 43 | CBDPS | G_31_KT_33_ | T_44_KEGVVH_50_ | K32-K45 |
| 44 | CBDPS | T_54_VAEKT_59_ | A_78_QKT_81_ | K58-K80 |
| 45 | CBDPS | M_5_K_6_ | S_9_KAKEGVVA_17_ | K6-K10 |
| 46 | CBDPS | Nterm-M_1_DVF_4_ | A_11_KEGVVA_17_ | N-K12 |
| 47 | CBDPS | K_97_DQLGK_102_ | Nterm-M_1_DVF_4_ | K97-N |
| 48 | CBDPS | M_5_K_6_ | S_42_KTKEGVVH_50_ | K6-K43 |
| 49 | CBDPS | M_5_K_6_ | K_60_EQVTNVGGA_69_ | K6-K60 |
| 50 | CBDPS | A_18_AEKT_22_ | T_44_KEGVVH_50_ | K21-K45 |
| 51 | CBDPS | K_80_TVEGA_85_ | G_93_FVKKD_98_ | K80-K97 |
| 52 | CBDPS | K_23_QGVAE_28_ | K_45_EGVVH_50_ | K23-K45 |
| 53 | CBDPS | K_34_EGVL_38_ | T_44_KEGVVH_50_ | K34-K45 |
| 54 | CBDPS | T_44_KEGVVH_50_ | V_55_AEKT_59_ | K45-K58 |
| 55 | CBDPS | M_5_KGL_8_ | A_19_EKTKQGVAE_28_ | K6-K21 |
| 56 | CBDPS | V_95_K_96_ | K_97_DQLGKNEEGAPQEGIL_113_ | K96-K102 |
| 57 | CBDPS | K_80_TVEGAGSIAAATGFVK_96_ | K_96_KDQLGK_102_ | K80-K96 |
| 58 | CBDPS | Nterm-M_1_DVF_4_ | A_11_KEGVVAAAEK_21_ | N-K21 |
| 59 | BS^3^ | Nterm-M_1_DVFMK_6_ | G_7_LSKAKEGVVAAAEK_21_ | N-K10 |
| 60 | BS^3^ | Nterm-M_1_DVFMKGLSK_10_ | A_11_KEGVVAAAEK_21_ | N-K12 |
| 61 | BS^3^ | Nterm-M_1_DVFMK_6_ | E_13_GVVAAAEKTKQGVAEAAGK_32_ | N-K21 |
| 62 | BS^3^ | Nterm-M_1_DVFMKGLSKAKEGVVAAAEK_21_ | T_22_KQGVAEAAGKTK_34_ | N-K23 |
| 63 | BS^3^ | Nterm-M_1_DVFMK_6_ | E_35_GVLYVGSKTKEGVVHGVATVAEK_58_ | N-K43 |
| 64 | BS^3^ | Nterm-M_1_DVFMK_6_ | T_44_KEGVVHGVATVAEKTK_60_ | N-K58 |
| 65 | BS^3^ | Nterm-M_1_DVFMK_6_ | T_59_KEQVTNVGGAVVTGVTAVAQK_80_ | N-K60 |
| 66 | BS^3^ | Nterm-M_1_DVFMK_6_ | T_59_KEQVTNVGGAVVTGVTAVAQKTVEGAGSIAAATGFVK_96_ | N-K80 |
| 67 | BS^3^ | Nterm-M_1_DVFMKGLSK_10_ | A_11_KEGVVAAAEK_21_ | K6-K12 |
| 68 | BS^3^ | Nterm-M_1_DVFMKGLSK_10_ | Q_24_GVAEAAGKTK_34_ | K6-K32 |
| 69 | BS^3^ | Nterm-M_1_DVFMKGLSK_10_ | T_59_KEQVTNVGGAVVTGVTAVAQK_80_ | K6-K60 |
| 70 | BS^3^ | Nterm-M_1_DVFMKGLSK_10_ | K_97_DQLGK_102_ | K6-K97 |
| 71 | BS^3^ | Nterm-M_1_DVFMKGLSKAK_12_ | E_13_GVVAAAEKTKQGVAEAAGK_32_ | K10-K21 |
| 72 | BS^3^ | Nterm-M_1_DVFMKGLSKAKEGVVAAAEK_21_ | T_22_KQGVAEAAGK_32_ | K10-K23 |
| 73 | BS^3^ | G_7_LSKAK_12_ | T_22_KQGVAEAAGKTK_34_ | K10-K32 |
| 74 | BS^3^ | A_11_KEGVVAAAEK_21_ | T_22_KQGVAEAAGKTK_34_ | K12-K32 |
| 75 | BS^3^ | A_11_KEGVVAAAEK_21_ | T_33_KEGVLYVGSK_43_ | K12-K34 |
| 76 | BS^3^ | A_11_KEGVVAAAEK_21_ | T_44_KEGVVHGVATVAEK_58_ | K12-K45 |
| 77 | BS^3^ | A_11_KEGVVAAAEK_21_ | T_44_KEGVVHGVATVAEKTKEQVTNVGGAVVTGVTAVAQK_80_ | K12-K58 |
| 78 | BS^3^ | A_11_KEGVVAAAEK_21_ | T_59_KEQVTNVGGAVVTGVTAVAQK_80_ | K12-K60 |
| 79 | BS^3^ | A_11_KEGVVAAAEK_21_ | T_59_KEQVTNVGGAVVTGVTAVAQKTVEGAGSIAAATGFVK_96_ | K12-K80 |
| 80 | BS^3^ | A_11_KEGVVAAAEK_21_ | T_81_VEGAGSIAAATGFVKK_97_ | K12-K96 |
| 81 | BS^3^ | G_7_LSKAKEGVVAAAEK_21_ | K_97_DQLGK_102_ | K12-K97 |
| 82 | BS^3^ | A_11_KEGVVAAAEKTK_23_ | Q_24_GVAEAAGKTK_34_ | K21-K32 |
| 83 | BS^3^ | E_13_GVVAAAEKTKQGVAEAAGK_32_ | T_33_KEGVLYVGSK_43_ | K21-K34 |
| 84 | BS^3^ | E_13_GVVAAAEKTK_23_ | T_59_KEQVTNVGGAVVTGVTAVAQK_80_ | K21-K60 |
| 85 | BS^3^ | A_11_KEGVVAAAEKTKQGVAEAAGK_32_ | T_59_KEQVTNVGGAVVTGVTAVAQK_80_ | K21-T72 |
| 86 | BS^3^ | E_13_GVVAAAEKTK_23_ | E_61_QVTNVGGAVVTGVTAVAQKTVEGAGSIAAATGFVK_96_ | K21-K80 |
| 87 | BS^3^ | A_11_KEGVVAAAEKTK_23_ | K_97_DQLGK_102_ | K21-K97 |
| 88 | BS^3^ | T_22_KQGVAEAAGK_32_ | T_33_KEGVLYVGSKTKEGVVHGVATVAEK_58_ | T22-K43 |
| 89 | BS^3^ | E_13_GVVAAAEKTKQGVAEAAGK_32_ | Q_24_GVAEAAGKTKEGVLYVGSK_43_ | K23-K32 |
| 90 | BS^3^ | T_22_KQGVAEAAG KTK_34_ | E_35_GVLYVGSK_43_ | K23-S42 |
| 91 | BS^3^ | T_22_KQGVAEAAGK_32_ | E_46_GVVHGVATVAEKTK_60_ | K23-K58 |
| 92 | BS^3^ | T_22_KQGVAEAAGK_32_ | T_59_KEQVTNVGGAVVTGVTAVAQK_80_ | K23-K60 |
| 93 | BS^3^ | T_22_KQGVAEAAGK_32_ | E_61_QVTNVGGAVVTGVTAVAQKTVEGAGSIAAATGFVK_96_ | K23-K80 |
| 94 | BS^3^ | Q_24_GVAEAAGKTK_34_ | E_35_GVLYVGSKTKEGVVHGVATVAEK_58_ | K32-K43 |
| 95 | BS^3^ | T_22_KQGVAEAAGKTK_34_ | E_46_GVVHGVATVAEKTK_60_ | K32-K58 |
| 96 | BS^3^ | Q_24_GVAEAAGKTK_34_ | T_59_KEQVTNVGGAVVTGVTAVAQK_80_ | K32-K60 |
| 97 | BS^3^ | T_22_KQGVAEAAGKTK_34_ | K_97_DQLGK_102_ | K32-K97 |
| 98 | BS^3^ | T_33_KEGVLYVGSK_43_ | T_33_KEGVLYVGSK_43_ | T33-K34 |
| 99 | BS^3^ | Q_24_GVAEAAGKTKEGVLYVGSK_43_ | T_44_KEGVVHGVATVAEK_58_ | K34-T44 |
| 100 | BS^3^ | T_33_KEGVLYVGSK_43_ | T_44_KEGVVHGVATVAEK_58_ | K34-T54 |
| 101 | BS^3^ | T_33_KEGVLYVGSK_43_ | T_44_KEGVVHGVATVAEK_58_ | K34-K45 |
| 102 | BS^3^ | T_33_KEGVLYVGSK_43_ | T_44_KEGVVHGVATVAEKTK_60_ | K34-K58 |
| 103 | BS^3^ | T_33_KEGVLYVGSK_43_ | T_59_KEQVTNVGGAVVTGVTAVAQK_80_ | K34-K60 |
| 104 | BS^3^ | T_33_KEGVLYVGSK_43_ | E_61_QVTNVGGAVVTGVTAVAQKTVEGAGSIAAATGFVK_96_ | K34-K80 |
| 105 | BS^3^ | T_33_KEGVLYVGSK_43_ | K_97_DQLGK_102_ | K34-K97 |
| 106 | BS^3^ | T_33_ KEGVLYVGSKTK_45_ | E_35_GVLYVGSKTKEGVVHGVATVAEK_58_ | T39-K43 |
| 107 | BS^3^ | T_33_KEGVLYVGSK_43_ | T_44_KEGVVHGVATVAEK_58_ | T39-K45 |
| 108 | BS^3^ | E_35_GVLYVGSK_43_ | E_46_GVVHGVATVAEKTK_60_ | T39-K58 |
| 109 | BS^3^ | E_35_GVLYVGSK_43_ | T_59_KEQVTNVGGAVVTGVTAVAQK_80_ | T39-K60 |
| 110 | BS^3^ | T_33_KEGVLYVGSKTK_45_ | E_61_QVTNVGGAVVTGVTAVAQKTVEGAGSIAAATGFVK_96_ | T39-K80 |
| 111 | BS^3^ | E_35_GVLYVGSK TK_45_ | T_44_KEGVVHGVATVAEKTK_60_ | S42-K45 |
| 112 | BS^3^ | E_35_GVLYVGSK_43_ | T_44_KEGVVHGVATVAEKTK_60_ | S42-K58 |
| 113 | BS^3^ | E_35_GVLYVGSKTKEGVVHGVATVAEK_58_ | T_59_KEQVTNVGGAVVTGVTAVAQK_80_ | K43-K60 |
| 114 | BS^3^ | T_33_KEGVLYVGSKTK_45_ | K_97_DQLGK_102_ | K43-K97 |
| 115 | BS^3^ | T_44_KEGVVHGVATVAEK_58_ | T_59_KEQVTNVGGAVVTGVTAVAQK_80_ | K45-K60 |
| 116 | BS^3^ | T_44_KEGVVHGVATVAEK_58_ | K_97_DQLGK_102_ | K45-K97 |
| 117 | BS^3^ | T_44_KEGVVHGVATVAEKTK_60_ | E_61_QVTNVGGAVVTGVTAVAQKTVEGAGSIAAATGFVK_96_ | K58-K80 |
| 118 | BS^3^ | T_44_KEGVVHGVATVAEKTK_60_ | K_97_DQLGK_102_ | K58-K97 |
| 119 | BS^3^ | T_59_KEQVTNVGGAVVTGVTAVAQK_80_ | K_97_DQLGK_102_ | K60-K97 |
| 120 | BS^3^ | T_59_KEQVTNVGGAVVTGVTAVAQKTVEGAGSIAAATGFVK_96_ | K_97_DQLGK_102_ | K80-K97 |
| 121 | BS^3^ | T_81_VEGAGSIAAATGFVK_96_ | K_97_DQLGK_102_ | S87-K97 |
| 122 | BS^3^ | E_35_GVLYVGSKTKEGVVHGVATVAEK_58_ | E_46_GVVHGVATVAEK_58_ | K43-T54 |
| 123 | BS^3^ | T_44_KEGVVHGVATVAEK_58_ | T_59_KEQVTNVGGAVVTGVTAVAQK_80_ | K45-T59 |
| 124 | BS^3^ | T_44_KEGVVHGVATVAEKTK_60_ | E_61_QVTNVGGAVVTGVTAVAQK_80_ | K58-T64 |
| 125 | BS^3^ | T_59_KEQVTNVGGAVVTGVTAVAQK_80_ | T_81_VEGAGSIAAATGFVK_96_ | K60-S87 |

**Table S5. The average secondary structure content (in percentage) of each cluster and the conformational ensemble.**

| Cluster | α-Helix | β-Strand | Turn | Other |
| --- | --- | --- | --- | --- |
| C1 | 5.1 | 44.2 | 16.5 | 34.2 |
| C2 | 11.4 | 32.4 | 21.4 | 34.8 |
| C3 | 1.2 | 38.0 | 18.3 | 42.5 |
| C4 | 2.8 | 41.6 | 23.1 | 32.5 |
| C5 | 10.5 | 32.7 | 20.6 | 36.2 |
| C6 | 11.1 | 31.7 | 24.6 | 32.6 |
| C7 | 3.8 | 40.2 | 19.7 | 36.3 |
| C8 | 8.2 | 33.1 | 23.4 | 35.3 |
| Ensemble average | 7.9 | 36.3 | 20.8 | 34.9 |
